# An oxytocin-gated circuit from the hypothalamus silences olfactory tubercle neurons to drive prosocial grooming

**DOI:** 10.64898/2026.03.27.714758

**Authors:** Yanbiao Zhong, Jian Yang, Yize Qi, Lulu Guo, Haiping Wang, Ding Wang, Xinghan Lin, Maoyuan Wang, Haishui Shi, Xinyu Nan, Haixuan Xu, Guanqing Li, Dawei Wang, Minghong Ma, Jian Mao, Yiqun Yu, Chanyi Lu, Yun-Feng Zhang

**Affiliations:** State Key Laboratory of Animal Biodiversity Conservation and Integrated Pest Management, Institute of Zoology, Chinese Academy of Sciences, Beijing 100101, China; Department of Rehabilitation Medicine, The First Affiliated Hospital of Gannan Medical University, Ganzhou 341000, China; Ganzhou Intelligent Rehabilitation Technology Innovation Center, Ganzhou 341000, China; Key Laboratory of Prevention and Treatment of Cardiovascular and Cerebrovascular Diseases (Gannan Medical University), Ministry of Education, Ganzhou 341000, China; Gannan Medical University, GanZhou 341000, China; University of Chinese Academy of Sciences, Beijing 100101, China; Flavor Science Laboratory, Beijing Life Science Academy (BLSA), Beijing, 102209, China, The Hebei Key Laboratory of Early Life Health Promotion, Nursing School, Hebei Medical University, Shijiazhuang 050031, China; Postdoctoral Research Station of Hebei Medical University, Shijiazhuang 050017, China; School of Nursing, Hebei Medical University, Shijiazhuang 050031, China; School of Life Sciences, Hebei University, Baoding 071002, China; State Key Laboratory for Biology of Plant Diseases and Insect Pests, Institute of Plant Protection, Chinese Academy of Agricultural Sciences, Beijing 100193, China; Institute of Western Agricultural of CAAS, Changji 831100, Xinjiang, China; Department of Neuroscience, Perelman School of Medicine, University of Pennsylvania, Philadelphia, PA 19104, USA; ENT Institute and Department of Otorhinolaryngology, Eye & ENT Hospital, Fudan University, Shanghai, 200031, China; Olfactory Disorder Diagnosis and Treatment Center, Eye & ENT Hospital, Fudan University, Shanghai 200031, China; State Key Laboratory of Molecular Developmental Biology, Institute of Genetics and Developmental Biology, Chinese Academy of Sciences, Beijing 100101, China

**Keywords:** Allogrooming, Oxytocin, Paraventricular nucleus, Olfactory tubercle, GIRK channels

## Abstract

Spontaneous helping behaviors such as allogrooming are vital for survival in social species, yet their underlying neural mechanisms remain largely unknown. Although oxytocin (OXT) is known to modulate allogrooming, the precise neural circuits, molecular pathways, and sensory drivers remain unclear. Here, using a social emergency paradigm in mice, we show that observer mice selectively allogroom distressed, anesthetized conspecifics, a behavior that is guided by main olfactory system cues and mitigates the demonstrator’s anxiety. We identify an oxytocinergic circuit from the paraventricular hypothalamus to the medial olfactory tubercle (PVN^OXT^→mOT) that is necessary and sufficient for this helping behavior. In vivo fiber photometry reveals that PVN^OXT^→mOT activity and OXT release are temporally locked to allogrooming initiation. This behavioral effect requires oxytocin receptor (OXTR) signaling specifically on mOT dopamine D1 receptor-expressing neurons, where OXTRs suppress neuronal excitability via G protein-gated inwardly rectifying K^+^ (GIRK) channels. Disruption of this inhibitory mechanism induces neuronal hyperexcitability and impairs allogrooming, a deficit rescued by restoring GIRK function. These findings define a PVN^OXT^-mOT circuit for prosocial helping, revealing an oxytocinergic pathway with potential therapeutic relevance for neuropsychiatric disorders characterized by social deficits.

## Introduction

Human life is characterized by complex social environments that necessitate reciprocal assistance, particularly in emergencies or life-threatening situations^1–3^. Spontaneous helping behaviors, such as rescue-like actions, are vital for the well-being and survival of social species^4–7^. Allogrooming, grooming directed at another individual, is a key prosocial behavior observed in rodents^4,8–12^ and primates, including humans^13,14^. It serves multiple physiological functions, such as maintaining hygiene^15,16^, orchestrating social dynamics^11,15–19^, and regulating emotion^4,13,20,21^. The effective performance of allogrooming, which underpins these functions, relies on specific neural substrates, including the neuropeptide oxytocin (OXT)^4,8,12,22^.

The OXT, primarily synthesized in the paraventricular (PVN) and supraoptic (SON) nuclei of the hypothalamus ^23,24^, is evolutionarily conserved across vertebrate species^25–29^ and acts as a regulator of various types of social behaviors^30–40^, including allogrooming^4,8,19,22^. OXT signaling is critical for diverse prosocial interactions, including allogrooming, the act of grooming a conspecific^41^. Recent work in mice indicates that PVN^OXT^ neurons coordinate the emotional and motor components of rescue-like behavior, including allogrooming, via distinct projections to the central amygdala (CeA) and dorsal bed nucleus of the stria terminalis (dBNST)^4^. Furthermore, these neurons respond differentially to social context, and their activity is necessary for reviving-like allogrooming behavior^8^. While these findings underscore a key role for OXT in organizing allogrooming, potentially through the integration of sensory social cues^42^, the specific sensory modalities (e.g., olfactory, visual, auditory) required for this regulation remain unclear. The precise neural and molecular mechanisms within PVN^OXT^ circuits that underlie allogrooming are thus still not fully defined.

PVN OXT (PVN^OXT^) neurons project broadly to brain regions including the thalamus, cortex, amygdala, hippocampus, and striatum^24,31,36,43–46^. Within the ventral striatum, which comprises the nucleus accumbens (NAc) and the olfactory tubercle (OT; also called tubular striatum^47^), the OT is populated primarily by GABAergic spiny projection neurons expressing dopamine D1 or D2 receptors (D1 or D2 SPNs), alongside sparse interneurons such as the D3 receptor-expressing neurons (D3 neurons) of the Islands of Calleja^47–50^. Like other olfactory regions [e.g., anterior olfactory nucleus (AON), piriform cortex (PIR)], the OT receives direct oxytocinergic input from the PVN^26,44^. Although OXT signaling is known to modulate social behavior through several olfactory structures, including the olfactory bulb^51^, AON^52,53^ and PIR^42^, its functional role in the OT remains unexplored. Anatomical studies confirm PVN projections to the OT^54,55^, and oxytocinergic fibers^24,56^, and oxytocin receptors (OXTRs)^57,58^ have been identified within this region, particularly in the Islands of Calleja. Together, these observations suggest that a functional PVN→OT oxytocinergic pathway may regulate allogrooming behavior in mice.

Herein, we combined optogenetic, chemogenetic, pharmacological, and genetic approaches with in vivo fiber photometry and behavioral assays to investigate the role of the PVN→OT oxytocinergic pathway in allogrooming. Our study aimed to determine: 1) whether mice actively perform allogrooming toward a distressed conspecific; 2) which sensory system guides this behavior; and 3) the underlying neural and molecular mechanism mediated by the PVN^OXT^–OT pathway. Using a sodium pentobarbital-induced anesthesia model to simulate an unconscious partner, we found that observer mice displayed a significant preference to allogroom anesthetized demonstrators, and that this allogrooming reduced anesthesia-induced anxiety-like behavior in the recipients/demonstrators. Pharmacological blockade revealed that the main olfactory system is necessary for this selective allogrooming. In vivo fiber photometry recorded time-locked increases in both PVN^OXT^-OT axonal activity and OT oxytocin release during an observer’s social investigation and allogrooming of an anesthetized demonstrator. Circuit manipulations showed that inhibiting the PVN^OXT^-OT pathway decreased allogrooming, while activating it increased this behavior. This regulation requires OXTR signaling, which suppresses the excitability of OT D1 neurons via G protein-gated inwardly rectifying K□ (GIRK/Kir3) channels. Collectively, our findings establish a critically facilitatory role for the PVN→OT oxytocinergic circuit in modulating prosocial allogrooming, thereby delineating a specific neural and molecular mechanism for oxytocin action within the central olfactory system.

## Materials and Methods

### Animals

The following mouse lines were used. C57BL/6 mice were purchased from Beijing Vital River Laboratory Animal Technology Co., Ltd., Beijing, China. Oxytocin-Ires Cre mice (strain # 024234; OXT-Cre) and Oxtr^flox/flox^ mice (B6.129(SJL)-*Oxtr^tm1.1Wsy^*/J; strain #008471; originally donated by Dr. W. Scott Young III -National Institute of Mental Health^59^) were from were obtained from the Jackson Laboratory. Bacterial artificial chromosome (BAC)-transgenic D1-Cre [MMRRC Tg(Drd1a-cre)EY262Gsat] and D2-Cre [MMRRC Tg(Drd2-cre)ER44Gsat] mice were purchased from MMRRC (Davis, California)^60^ and backcrossed to a C57BL/6 background. The D1-Cre line was crossed with the Cre-dependent tdTomato reporter line (JAX stock no. 007909 or Ai9 line B6.Cg-Gt(ROSA)26Sor^tm9(CAG-tdTomato)^ ^Hze/^). BAC D3-Cre line (stock B6.FVB(Cg)-Tg(Drd3-cre)KI198Gsat/Mmucd; RRID: MMRRC_031741-UCD) line was obtained from the Mutant Mouse Resource and Research Centers (MMRRC) at University of California at Davis, a National Institutes of Health (NIH)-funded strain repository, and was donated to the MMRRC by N. Heintz (The Rockefeller University, GENSAT) and C. Gerfen (NIH, National Institute of Mental Health). Mice were group-housed (5 per cage; 20-26 °C, 40-70% humidity, 12 h light/12 h dark cycle) with food and water available *ad libitum*, until the surgery of receiving virus injection and intracranial implantation and housed afterwards with 3-4 mice in one cage. All animal procedures complied with governmental regulations in China and were approved by the Institutional Animal Care and Use Committee of the Institute of Zoology, Chinese Academy of Sciences (IOZ-IACUC-2022-230). Male mice (8-12 weeks old) were recruited in the study.

### Viral injection and optical fiber/cannula implantation

Mice were anesthetized with isoflurane (induction: 5%, maintenance: 1-2%; RWD, cat# R510-22-10) and secured on a stereotaxic apparatus (RWD, Shenzhen, China; Model 68018). Ophthalmic ointment were applied to the eyes to protect the corneas from drying, and a heating pad was placed under the abdomen to maintain body temperature. Following treated with iodophor (Shandong Lircon Medical Technology Co., Ltd., Shandong, China), a 2-3 cm midline incision was made in the scalp using sterile surgical scissors (RWD, cat# S14014-10) to expose the cranial bones. The skull was leveled horizontally, and a handheld skull drill (RWD, Model 78001) was used to create a hole targeting the desired brain region. For viral injection (250 nl each side), rAAV2/9-hSyn-DIO-stGtACR2-EGFP (2.00×10^12^ vg/ml; Brain Case, cat# BC-1128), rAAV2/9-hSyn-DIO-hChR2-EGFP (2.00×10^12^ vg/ml; Brain Case, cat# BC-1397), rAAV2/9-hSyn-DIO-hM4D(Gi)-mCherry (2.00×10^12^ vg/ml; Brain Case, cat# BC-0153), rAAV2/9-hSyn-DIO-hM3D(Gq)-mCherry (2.50×10^12^ vg/ml; Brain Case, cat# BC-0143), rAAV2/9-EF1α-DIO-Axon-GCaMP6s (2.00×10^12^ vg/ml; Brain Case, cat# BC-0197), rAAV2/9-hSyn-OXT1.0 (2.00×10^12^ vg/ml; Brain Case, cat# BC-0293), rAAV2/9-hSyn-Cre-EGFP (2.00×10^12^ vg/ml; Brain Case, cat# BC-0160), rAAV2/9-D1-Cre-mCherry (4.98**×**10^12^ vg/ml; Brain VTA, cat# PT-0960), rAAV2/9-D2-Cre-EGFP (4.59**×**10^12^ vg/ml; Brain VTA, cat# PT-0813), rAAV2/9-EF1α-DIO-taCasp3-EGFP (5.16×10^12^ vg/ml; Brain Case, cat# BC-1142), rAAV2/9-hSyn-EGFP (2.00×10^12^ vg/ml; Brain VTA, cat# PT-0241), rAAV2/9-hSyn-DIO-mCherry (2.00×10^12^ vg/ml; Brain VTA, cat# PT-0115), rAAV2/9-hSyn-DIO-EGFP (5.00 × 10^12^ vg/ml; Brain Case, cat# BC-0244), rAAV-EF1α-DIO-EGFP (2.00 × 10^12^ vg/ml; Brain Case, cat# BC-0015), or rAAV2/9-hSyn-DIO-hKir3.2-EGFP (5.00 x 10^12^ vg/ml; Brain case, cat# BC-5517), was injected into the PVN or mOT, unilaterally or bilaterally where as appropriate, of OXT-Cre, Oxtr^flox/flox^ or C57BL/6 mice. Viral solutions were injected into the target brain region at a rate of 50 nl/min using a microinjection pump (Harvard Apparatus, cat# 70-4507) connected to a 10 µl microsyringe (Hamilton Company, USA) with a glass needle (RWD, cat# GC-3.5). The syringe tip was left in place for 10-15 min post-injection, to permit viral diffusion. As appropriate, an optical fiber (Ø200 μm, 0.37 NA; Inper, cat# 295N8; unilateral) or cannula (Ø410 µm core, 6 mm length; RWD, cat# 62004; bilateral) was then implanted into the targeted brain region. To stabilize the implant, two small screws (Xinghua Tianzhuo metal products Co., Ltd., China) were attached to the skull and secured with biotissue glue (3M, cat# 1469SB) and dental cement (Yamahachi Dental Industry (Changshu) Co., Ltd, China). Antibacterial agents were applied to the incision site to prevent infection. For pharmacologyical drug administration (200 nl each side), as appropriate, the OTXR selective antagonist L-368899 (1 μg/μl; Saitong, cat# S10501, agonist oxytocin (1 μg/μl; Saitong, cat# S10501), or saline (0.9% NaCl; as the control) was bilaterally administered into the mOT, followed by behavioral testing 30 minutes post-injection. Behavioral tests were conducted at least three weeks after viral injection. Stereotaxic coordinates (relative to bregma): the PVN (AP: -0.72 mm; ML: ±0.25 mm; DV: -4.95 mm), the mOT (AP: +1.54 mm; ML: ±1.15 mm; DV: -5.25 mm), and the AON: (AP: +2.68; ML: ±1.15; DV: -3.05 mm). Viral injection/expression sites and optical fiber/cannula locations were verified postmortem (**Fig. S19-20**), and only mice with correct targeting were included in the analysis.

### Optogenetic and chemogenetic manipulations

For optogenetic experiments, blue light at 20 Hz (470 nm, 10-15 mW/mm², 10 ms pulse; Optogenetic activation) or shining continuously (optogenetic inhibition) was applied to activate ChR2-expressing or inhibit stGtACR2-expressing PVN^OXT^ neuronal fibers in the mOT, respectively, during behavioral tests. EGFP-expressing mice were used as the controls. OXT-Cre mice with bilateral injection of excitatory/inhibitory designer receptors exclusively activated by designer drugs rAAV-hSyn-DIO-hM3D(Gq)-mCherry (excitatory DREADDs), rAAV-hSyn-DIO-hM4D(Gi)-mCherry (inhibitory DREADDs), or rAAV-hSyn-DIO-mCherry (control virus) in the PVN and bilateral cannula implantation in the mOT were adminstered with the chemogenetic ligand clozapine N-oxide (CNO, 3 μg/μl; Brain Case, cat# CNO-25 mg) or saline (0.9% NaCl; as a control) into the mOT, 30 min before behavioral tests.

### Behavioral assays

All behavioral experiments were performed during the light cycle (9:00 am-12:00 pm). Prior to test, mice were pair-housed and cohabited for two weeks except for the unfamiliar partner test in which the two mice used were novel reciprocally. All mice tested were habituated to both the experimenter and testing environment for three consecutive days. Mice were transferred to the testing room 1 h before the test for habituation. Optogenetic manipulations and local CNO administration (bilateral infusion; 200 nL) were applied as appropriate. Behaviors were videotaped for post-hoc analysis using EthoVision XT 16 software (Noldus) or PotPlayer64-bit software or analyzing manually as appropriate. To minimize bias, experimenters were blinded to group assignments during behavioral scoring.

### Allogrooming test

During testing, one mouse was randomly designated as the observer, while the other served as the demonstrator. The demonstrator was anesthetized via intraperitoneal injection of pentobarbital sodium (0.5%, 75 mg/kg). At 10-15 minutes post-injection, the anesthetized demonstrator was placed at one side of the home cage, and then the observer was re-introduced into the cage. To determine whether the main olfactory system is involved in allogrooming, observers received intraperitoneal injections of either methimazole (75 mg/kg; Harveybio, cat# SR2795; behavioral tests conducted 4 days post-injection) or saline (as the control). Observer’s allogrooming and social investigation toward the anesthetized demonstrator, as well as non-social behaviors (self-grooming and rearing) were videotaped for 10 minutes. The test of the observer’s behaviors was conducted in the home cage or a novel cage with familiar or unfamiliar demonstrators as appropriate. Recordings were post-analyzed to quantify the observer’s allogrooming and social investigation toward the anesthetized demonstrator, as well as non-social behaviors (self-grooming and rearing).

As appropriate, in the home cage, an observer was physically isolated with a toy mouse or an anesthetized demonstrator by an opaque divider containing twelve evenly distributed holes (diameter: 0.5) for 10 minutes (Non-contact). The holes on the divider allowed the disperse of olfactory cues from the toy mouse or anesthetized demonstrator while preventing visual cues. In another context, the divider was taken away, and the observer was allowed to socially contact with the anesthetized demonstrator for 10 minutes (social contact). After the recovery of the demonstrators, behaviors of these observers and demonstrators in the open field test (OFT) and light-dark box transition test (LDT) were monitored to check their emotional-state.

For the odorant transfer induced-allogrooming test, five mice were group-housed for two weeks prior to experimentation. One mouse was randomly designated as the observer, the other one as the demonstrator, and another two as the odorant donor mice (one awake and the other anesthetized). Odorants were collected by gently swabbing the donor mouse’s body (orofacial and perianal regions, head, tail, and body trunk) with a mineral oil-moistened cotton swab from the awake or anesthetized donor mouse, respectively. These odorants sourced-cotton swabs were immediately used to swab the orofacial part of the awake demonstrator, and observer’s allogrooming toward the demonstrator was videotaped for 10 minutes. Recordings were analyzed post-hoc to quantify latency to the first allogrooming onset, number of allogrooming bouts, and duration of allogrooming. Allogrooming, is defined as a mouse (observer) using its mouth licking the body surface (orofacial part, head, body trunk, tail, and anogenital area) of a conspecific (demonstrator/recipient), which triggers movements in the recipient that can be detected by the frame-by frame differentiation.

### Two-choice social interaction test

The test was conducted in the home cage in two distinct contexts. In context one (contact condition), one anesthetized demonstrator and one toy mouse were diagonally positioned in the cage with the observer in between. Observer’s behaviors were videotaped for 10 minutes. Recordings were analyzed post-hoc to quantify observer-directed social behaviors (allogrooming and investigation) toward the demonstrator and non-social behaviors (self-grooming and rearing) versus the toy mouse. In context two (non-contact condition), five mice was group-housed with for two weeks. One mouse was randomly designated as the observer, the other two as the demonstrators, and the remaining two as the odorant donor mice (one awake and the other anesthetized). Odorants were collected from the awake or anesthetized donor mouse, respectively, as aforementioned, and these odorants sourced-cotton swabs were immediately used to swab the orofacial part of the awake demonstrator. The two awake demonstrators with donor mouse’s odorants were separately put in a medical gauze-covered wire cup, diagonally located in the home cage. Observer’s behaviors were videotaped for 10 minutes. Recordings were analyzed post-hoc to quantify time spent and distance travelled around the cup.

### Three-chamber social interaction test

The test was conducted in an apparatus (60 × 40 × 20 cm) comprising two side chambers (20 × 40 × 20 cm each) and a central chamber (20 × 40 × 20 cm). The experiment contains three phases. Phase 1 (habituation): Subject mouse freely explored the empty apparatus for 10 minutes. Phase 2 (sociability test): A familiar mouse (S1) within a wire cup and an empty wire cup (E) were diagonally placed in the two side chambers. The subject mouse introduced and allowed to freely explore for 10 minutes. Phase 3: (social novelty test): A novel mouse (S2) was introduced into the empty wire cup that in phase 2. The subject mouse was allowed to freely explore for another 10 minutes. Behavioral metrics were calculated as following: Sociability preference index = [investigation time (S1) - investigation time (E)] / [investigation time (S1) + investigation time (E)]; Social novelty preference index = [investigation time (S2) - investigation time (S1)] / [investigation time (S2) + investigation time (S1)]. The locations of S1 and E, as well as S1 and S2, were counterbalanced between different subject mice tested.

### Buried food-seeking test

The hM4Di-, hM3Dq-, ChR2-, or stGtACR2-injected, and mice with D3 neurons ablation, OXTR KO or antagonism (L-368899; 1 μg/μl; bilateral infusion, 200 nL) in the OT were used. The experiment was conducted as previously described^61,62^. Briefly, mice were food-deprived for 24 h with ad libitum access to water prior to testing. During the test, a small piece of food pellet (∼2 g) was randomly buried in the corner of the cage (30 × 20 × 12 cm), 1.5 cm beneath the bedding (6 cm deep). The mouse was placed diagonally from the food pellet. The test lasted 10 minutes. Latency to food finding was defined as the time required for the mouse to locate the buried food pellet and initiate burrowing. Fresh cages and bedding were used for each test. Mice were released from an identical starting position, while pellet placement was randomized across trials.

### Open field test (OFT)

The OFT was performed in an open field arena (40 × 40 cm), comprising a central zone (20 × 20 cm) and a peripheral zone. The experimental mouse was promptly placed in the central zone and allowed to freely explore the arena for 5 minutes. The total distance travelled, distance travelled in the center zone, time spent in the center zone, and frequency of entries into the center zone were calculated.

### Light-dark box transition test (LDT)

The light-dark box test was conducted in a 45 × 28 × 30 cm apparatus, comprising a dark chamber (15 × 28 × 30 cm) and a light chamber (30 × 28 × 30 cm), with a dark-to-light area ratio of 1:2. Mice were placed in the center of the light box with the head oppositely facing the dark compartment, and were allowed to freely explore the two compartments for 10 min. The total time spent in the dark compartment was calculated.

### In vivo fiber photometry

To monitor dynamic activities of PVN^OXT^ neuronal fibers and OXT changes in the mOT, mice were anesthetized with 1-2% isoflurane and immobilized on a stereotaxic apparatus. A total of 250 nl of rAAV2/9-EF1α-DIO-Axon-GCaMP6s or rAAV2/9-hSyn-DIO-EGFP, and rAAV2/9-hSyn-OXT1.0 or rAAV2/9-hSyn-EGFP was unilaterally injected into the mOT (AP: +1.54 mm; ML: ±1.15 mm; DV: -5.25 mm) of OXT-Cre mice or C57BL/6 mice as appropriate. An optical fiber (Ø200 μm, 0.37 NA; Inper, cat# 295N8) was implanted ∼100 μm above the injection site and fixed with dental cement. After three weeks to allow for viral expression, the optical fiber was connected to a fiber photometry system (RWD, Model R821) via a patch cable. To minimize motion artifacts, the system simultaneously recorded signals stimulated by a 410 nm light source. The 470 nm laser power at the fiber tip was adjusted to 50 µW to avoid photobleaching.

For recordings in the allogrooming test, GCaMP6s or OXT sensor-expressing observer was introduced into a cage with an anesthetized demonstrator. GCaMP6s fluorescence signals or OTX sensor-dependent fluorescence signals in the OT were monitored for 10 minutes during observer socially interacting (allogrooming and investigation) with the demonstrator or performing non-social behaviors (rearing and walking). For recordings in the two-choice social interaction test, GCaMP6s or OXT sensor-expressing observer was introduced into a cage with an anesthetized demonstrator and a toy mouse located diagonally. GCaMP6s fluorescence signals or OTX sensor-dependent calcium signals in the OT were monitored for 10 minutes during observer interacting (allogrooming and investigation) with the demonstrator/toy mouse or performing non-social behaviors (rearing and walking). Videos were recorded to synchronize calcium fluorescence signals with specific behaviors as appropriate. Allogroomimg initiation was defined as the time point observer contacting and frequently scratching any body region of the demonstrator. Investigation initiation was defined as the time point observer positioning themselves within 2 cm of the demonstrator or toy mouse while facing it and displaying sustained sniffing behavior. Rearing initiation was defined as the time point observer’s forelegs lifting away from the cage floor and keeping upright. Walking initiation was defined as the time point observer switching from other behavioral states with its four legs completely touching the cage floor and keeping moving.

GCaMP6s fluorescence signals or OTX sensor-dependent calcium signals were excited by a 470 nm LED, and calcium-independent signals were excited by the 410 nm LED, with both light sources alternating at 30 Hz. Calcium-dependent fluorescence changes were quantified as ΔF/F using multichannel fiber-photometry software (v2.0.0.32640). ΔF/F was calculated as (baseline corrected 470 nm-fitted 410 nm (baseline corrected 410 nm))/median (baseline). The median (baseline) denoted the median value of the raw 470 nm signal during the 5-second period before investigation. The area under the curve (AUC) was calculated based on ΔF/F during the specific behavior period (-3-0 and 0-3 seconds) to compare responses of pre- versus post-initiation.

### Patch-clamp recording

Whole-cell patch-clamp recordings were performed as we previously described^54,63,64^. Briefly, the dissected brain was immediately placed in ice-cold cutting solution containing (in mM) 110 choline chloride, 2.5 KCl, 0.5 CaCl_2_, 7 MgCl_2_, 1.3 NaH_2_PO_4_, 25 NaHCO_3_, 10 glucose, 1.3 Na-ascorbate, and 0.6 Na-pyruvate (osmolality ∼300 mOsm, pH ∼7.4), bubbled with 95% O_2_ and 5% CO_2_. Coronal slices (250 µm thick) containing the OT were prepared using a Leica VT 1200S vibratome. Slices were incubated in oxygenated artificial cerebrospinal fluid (ACSF; in mM: 125 NaCl, 2.5 KCl, 2 CaCl_2_, 1.3 MgCl_2_, 1.3 NaH_2_PO_4_, 25 NaHCO_3_, 10 glucose, 1.3 Na-ascorbate, and 0.6 Na-pyruvate; osmolality ∼300 mOsm, pH ∼7.4), bubbled with 95% O_2_ and 5% CO_2_ for ∼60 min at 33 °C and then at room temperature for at least 40 min before recording.

Slices were visualized under an upright microscope (Eclipse FN1, Nikon, Japan) with a 40× water-immersion objective and equipped with wide-field fluorescence to identify fluorescently labelled neurons. Whole-cell recordings were obtained with a Multiclamp 700B amplifier and a Digidata 1550B analog-to-digital converter (Molecular Devices, San Jose, CA, USA). Recording pipettes were made from borosilicate glass with a Flaming-Brown puller (P-97, Sutter Instruments; tip resistance 6-8 MΩ). Whole-cell recordings were obtained to measure action potentials (APs) using depolarizing current steps in 10-pA increments. APs and spontaneous excitatory postsynaptic currents (sEPSCs) were recorded with a potassium-based internal solution containing (in mM): 135 K-gluconate, 5 NaCl, 0.5 CaCl□·2H□O, 5 EGTA, 10 HEPES, 2 Mg-ATP, and 0.1 Na-GTP (osmolality ∼300 mOsm, pH ∼7.3). During these recordings, the membrane potential was held at −70 mV, and the GABA_A_ receptor antagonist bicuculline (10 μM) was applied in the ACSF. Recordings of spontaneous inhibitory postsynaptic currents (sIPSCs) were performed with the membrane potential held at 0 mV, and CNQX (10 μM) and AP5 (50 μM) were added to block AMPA receptors and NMDA receptors, respectively. A cesium-based internal solution with a high-Cl internal solution was used, containing (in mM): 125 CsMeSO□, 7 CsCl, 5 QX-314·Cl, 10 HEPES, 0.5 EGTA, 0.3 Na□GTP, 4 Mg-ATP, and 10 phosphocreatine (osmolality ∼300 mOsm, pH ∼7.3). Viral infection in the OT was confirmed in brain slices during recording. As appropriate, oxytocin (1 μM) was bath perfused at 1-2 ml/min. Compounds used were purchased from Sigma (USA).

### Quantitative real-time PCR (qPCR)

Mice were immediately decapitated under anesthesia with isoflurane and OT samples were immediately collected on ice. Total RNA was extracted and quantified. Briefly, the RNA extraction was performed with Cell Total RNA Isolation Kit (Vazyme, cat# R433, Beijing, China) and RNA samples were treated with DNAse (DNase I, Amplification Grade; Life Technologies) to avoid genomic DNA contamination. The cDNA synthesis was performed with cDNA Reverse Transcription kit (Vazyme, cat # Q713, Beijing, China), according to the manufacturer’s instructions. The relative abundance of transcripts was analyzed using SYBR Green PCR Master Mix for amplification reaction (Applied Biosystems®, Life Technologies Corporation). Data were analyzed using the fold change (2^−ΔΔCt^) relative to the control group. The level of OXTR mRNA expression was normalized to the reference gene (Gapdh) mRNA expression. The primers used are as follows: Oxtr F: TCGTACTGGCCTTCATCGTG; Oxtr R: TGAAGGCAGAAGCTTCCTTGG. Gapdh F: CATCAAGAAGGTGGTGAAGCA; Gapdh R: CTGTTGAAGTCACAGGAGACA.

### Statistical analysis

Shapiro-Wilk tests were used to assess data normality. Parametric tests (Student’s t-test or one-way repeated-measures ANOVA) were applied to normally distributed data; otherwise, non-parametric tests (Mann-Whitney or Wilcoxon matched-pairs signed rank test) were used. Statistical analyses were performed in GraphPad Prism 8, and figures were assembled in Adobe Illustrator.

## Results

### Mice preferentially allogroom anesthetized conspecifics

To investigate whether mice exhibit rescue-like behaviors such as allogrooming, we utilized a sodium pentobarbital-induced anesthesia model to simulate an unconscious, unresponsive conspecific (demonstrator). We then quantified the behavior of a partner (observer) toward them. In a two-choice social interaction test (**Fig. 1a**), observers interacted more with an anesthetized demonstrator than with a toy mouse (**Fig. 1b**), which made the demonstrator recover faster, as reflected by the shorter latency to initial movement upon the presence versus absence of the observer (**Fig. 1c**). The observer spent a significantly higher percentage of time in the demonstrator’s chamber (**Fig. 1d**), further indicating a preference toward anesthetized demonstrator. This preference was characterized by increased allogrooming and social investigation towards the demonstrator, as well as more non-social behaviors (self-grooming and rearing) in that chamber compared to the side with the toy mouse (**Fig. 1b, e-h**).

**Figure 1.**
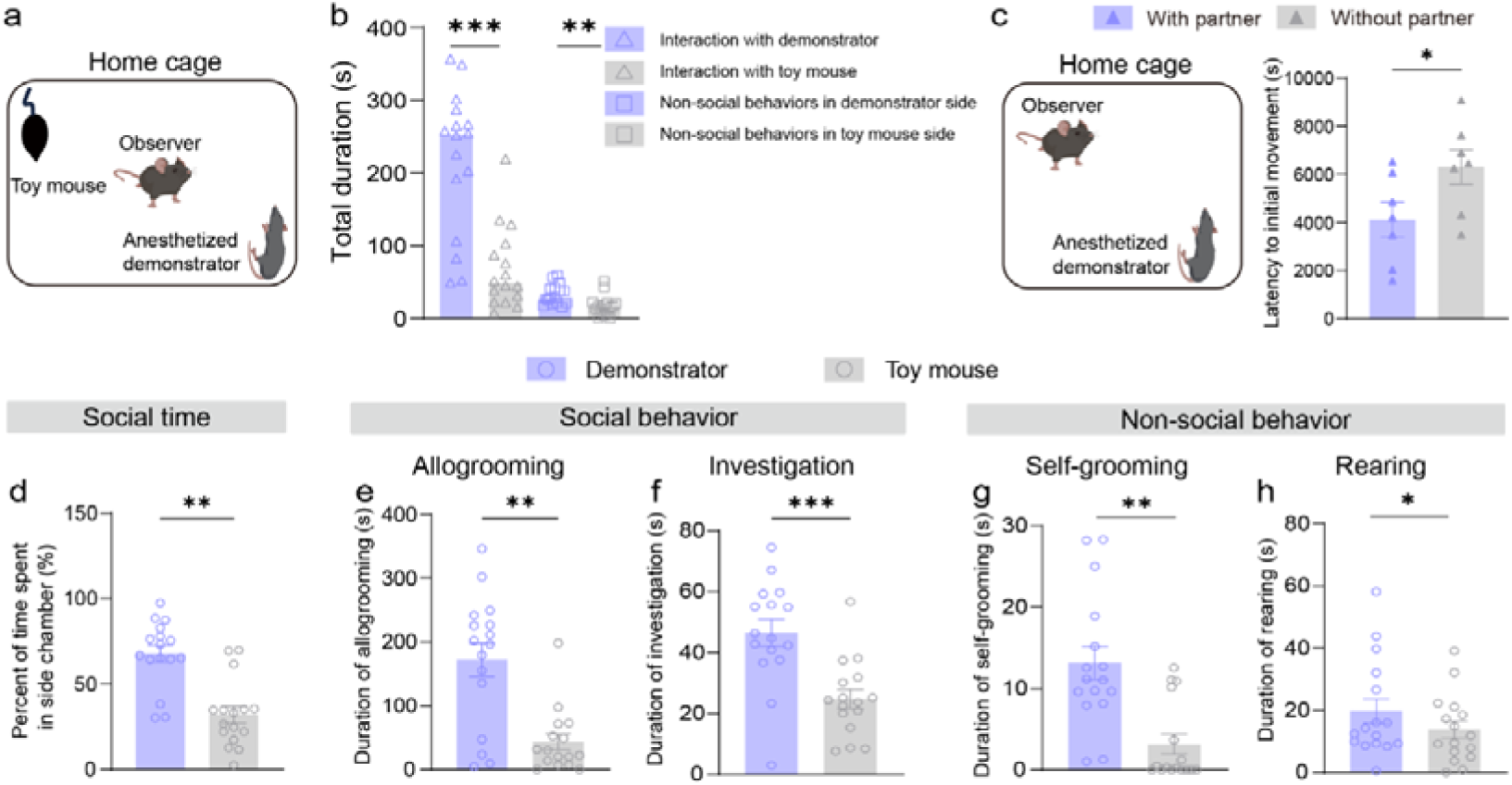
Mice preferentially allogroom anesthetized conspecifics in a two-choice social interaction test. **a** Schematic illustrating the two-choice social interaction test. **b** Total duration of social and non-social behaviors directed towards a conspecific (demonstrator) or an inanimate object (toy). **c** Left, Schematic showing the behavioral paradigm. Right, Latency to initial movement of demonstrator to recover from anesthetized state with/without partner (observer). **d** Fraction of time spent exploring the demonstrator versus the toy. **e, f** Duration of social behaviors, including allogrooming (**e**) and investigation (**f**). **g, h** Duration of non-social behaviors, including self-grooming (**g**) and rearing (**h**). Data are from the same observer and demonstrator cohorts (n = 16 mice per group) and are presented as mean ± SEM. \**p*□<□0.05, \*\**p*<□0.01, \*\*\**p*□<□0.001. Statistical details see Table S1.

Given the importance of emotional contagion in social animals including rodents^65–68^, we next assessed the bidirectional influence between the distressed demonstrator and the observer. Observers became hyperactive, but not more anxious, when in the presence of an anesthetized demonstrator, even without direct physical contact (**Fig. S1a-h**). Notably, the observer’s allogrooming significantly reduced the demonstrator’s anesthesia-induced anxiety-like behaviors. This anxiolytic effect was absent under non-physical contact conditions (**Fig. S1i-o**), indicating that allogrooming itself is essential for normalizing the demonstrator’s anxiolytic state.

Since familiarity influences social behaviors including allogrooming in rodents^8,10,69^, we tested its effect on the observer’s response. Observers displayed significantly more allogrooming and social investigation toward a cagemate demonstrator in a familiar home cage environment than in an unfamiliar new cage (**Fig. S2a-b**). In contrast, the levels of allogrooming and social investigation were similar toward a cagemate versus a stranger demonstrator (**Fig. S2e-f**). This pattern was also observed for non-social behaviors (self-grooming and rearing; **Fig. S2c-d, g-h**). Together, these results indicate that familiarity with the environment, but not with the partner, modulates the observer’s allogrooming and non-social behaviors toward an anesthetized conspecific.

To determine whether a demonstrator’s anesthetized state influences allogrooming preference, we conducted a two-choice test using awake versus anesthetized demonstrators. Observers displayed a significant spatial preference for anesthetized demonstrators, spending more time and travelling a greater distance near their cup (**Fig. 2a-b**). To identify the responsible sensory cues, we first assessed the role of vision using gauze-wrapped cups that blocked visual and physical access but permitted odor dispersion (**Fig. 2c**). Observers maintained their preference under these conditions (**Fig. 2d**), indicating that visual cues are dispensable. Given the importance of olfaction in rodent social behavior, we next investigated its role. Saline-treated control observers exhibited a strong preference for the anesthetized demonstrator, reflected in their locomotion tracks (**Fig. 2e**), increased time spent, and greater distance travelled near its cup **(Fig. 2f)**. However, this preference was abolished following ablation of the main olfactory epithelium (MOE) with methimazole^70,71^ (**Fig. 2e-f**). Furthermore, consistent with reduced perception of volatile odors^70^, methimazole-treated observers showed drastically reduced allogrooming, characterized by increased latency to onset, fewer bouts, and shorter duration (**Fig. 2 g-j**). Since methimazole ablates the MOE but spares the vomeronasal organ^71^, we conclude that the main olfactory system is necessary for the biased allogrooming toward anesthetized demonstrators.

**Figure 2.**
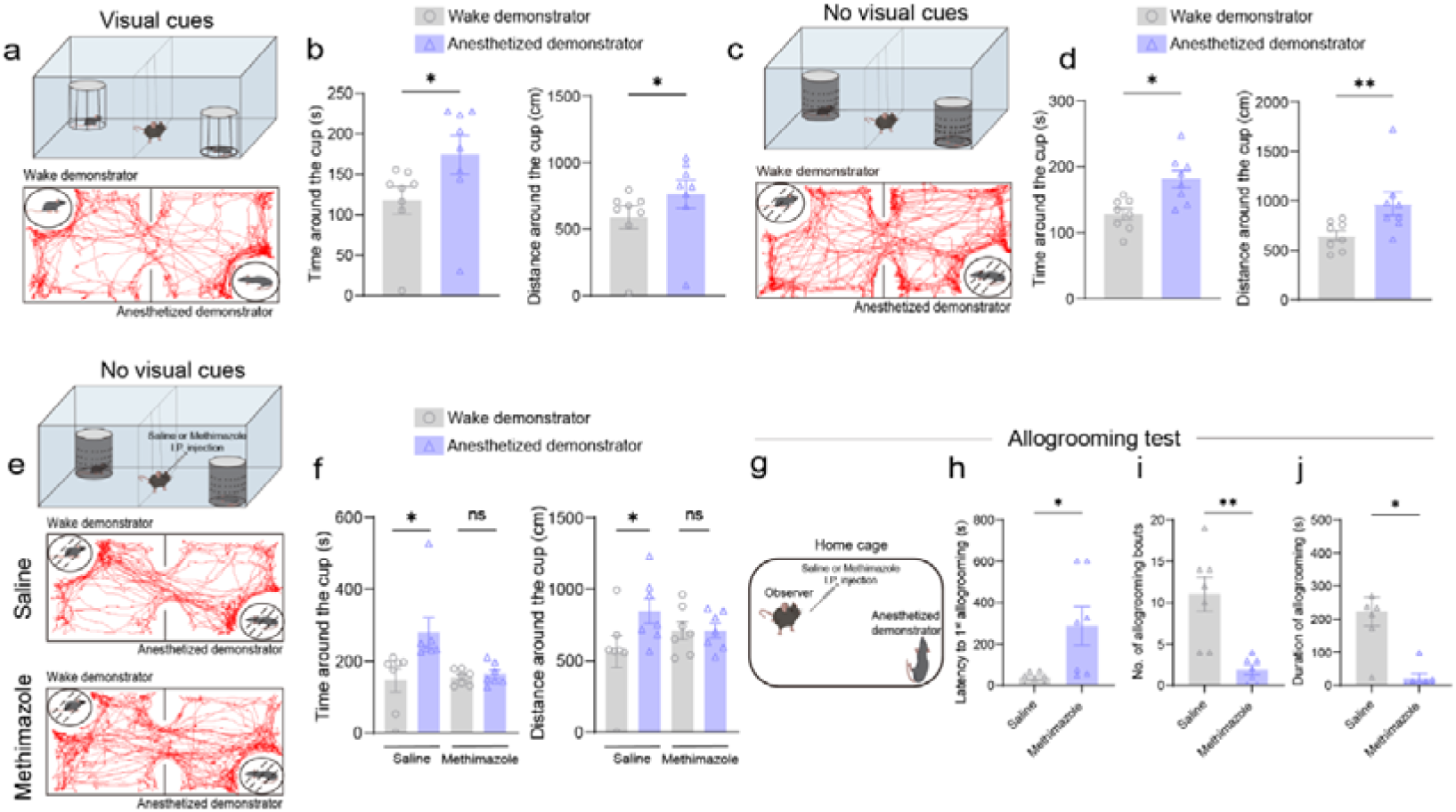
The main olfactory system is necessary for observer’s preferential allogrooming toward anesthetized demonstrator. **a** Behavioral paradigm schematic (top) and representative observer locomotion tracks (bottom): anesthetized versus awake demonstrator with visual cues. **b** Duration (left) and travelling distance (right) of observer exploration near stimulus cup (visual cues present). **c-d** Parallel tests without visual cues: paradigm schematic (top) and locomotion traces (bottom) (**c**) and cup exploration duration and travelling distance (**d**). **e-f** Observer responses after pharmacological treatment (saline versus methimazole): paradigm schematic (top) and locomotion traces (bottom) (**e**) and cup exploration duration and travelling distance (**f**). **g-j** Allogrooming test. **g** Behavioral schematic. **h-j** Allogrooming by treated observers toward anesthetized demonstrators: latency to first event (**h**), event frequency (**i**) and total duration (**j**). Observers: n = 8 (**b, d**), demonstrators: n = 16 (**b, d**). Pharmacologically treated observers (n = 7; **f**, **h-j**), demonstrators: n = 14 (**f**) and a subcohort n = 7 (**h-j**). Data are presented as mean ± SEM. \**p*□<□0.05, \*\**p*□<□0.01; ns, no significance. Statistical details see Table S1.

Noting that allogrooming was primarily directed at the orofacial region, a site rich in secretory glands in rodents^71,72^, we hypothesized that odorants from anesthetized demonstrators drive this behavior. To test this, we transferred odorants sampled from the orofacial region of anesthetized or awake demonstrators onto naïve conspecifics. In a non-social context (no direct contact), observers investigated both stimulus animals equally. However, in a social contact context, observers displayed significantly enhanced allogrooming toward conspecifics bearing anesthetized-demonstrator odorants compared to those bearing awake-demonstrator odorants (**Fig. S3**). This indicates that both olfactory cues and social contact are necessary for orchestrating allogrooming behavior via the detection of demonstrator-derived odorants.

### The PVN^OXT^-OT pathway modulates allogrooming toward anesthetized conspecifics

Oxytocin (OXT) is known to regulate allogrooming^4,8,19,22^, and the olfactory tubercle (OT), a key component of the main olfactory system, receives oxytocinergic projections from the paraventricular nucleus of the hypothalamus (PVN)^26,44^. We therefore hypothesized that the PVN^OXT^-OT pathway modulates allogrooming directed toward anesthetized conspecifics. To test this, we first inhibited the pathway chemogenetically. We bilaterally expressed Cre-dependent inhibitory DREADDs (hM4Di) or control virus in the PVN and implanted cannula in the medial OT (mOT) of OXT-Cre mice (**Fig. 3a**). Following a three-week expression period, behavioral assays were conducted (**Fig. 3 b-c**). In hM4Di-expressing observers, CNO administration significantly suppressed allogrooming during a direct social test (**Fig. 3d**). In a two-choice paradigm, chemogenetic inhibition selectively diminished time spent allogrooming an anesthetized demonstrator, but not a toy (**Fig. 3e**). This inhibition also significantly reduced sociability preference, while social novelty preference remained unaffected (**Fig. 3f–g)**.

**Figure 3.**
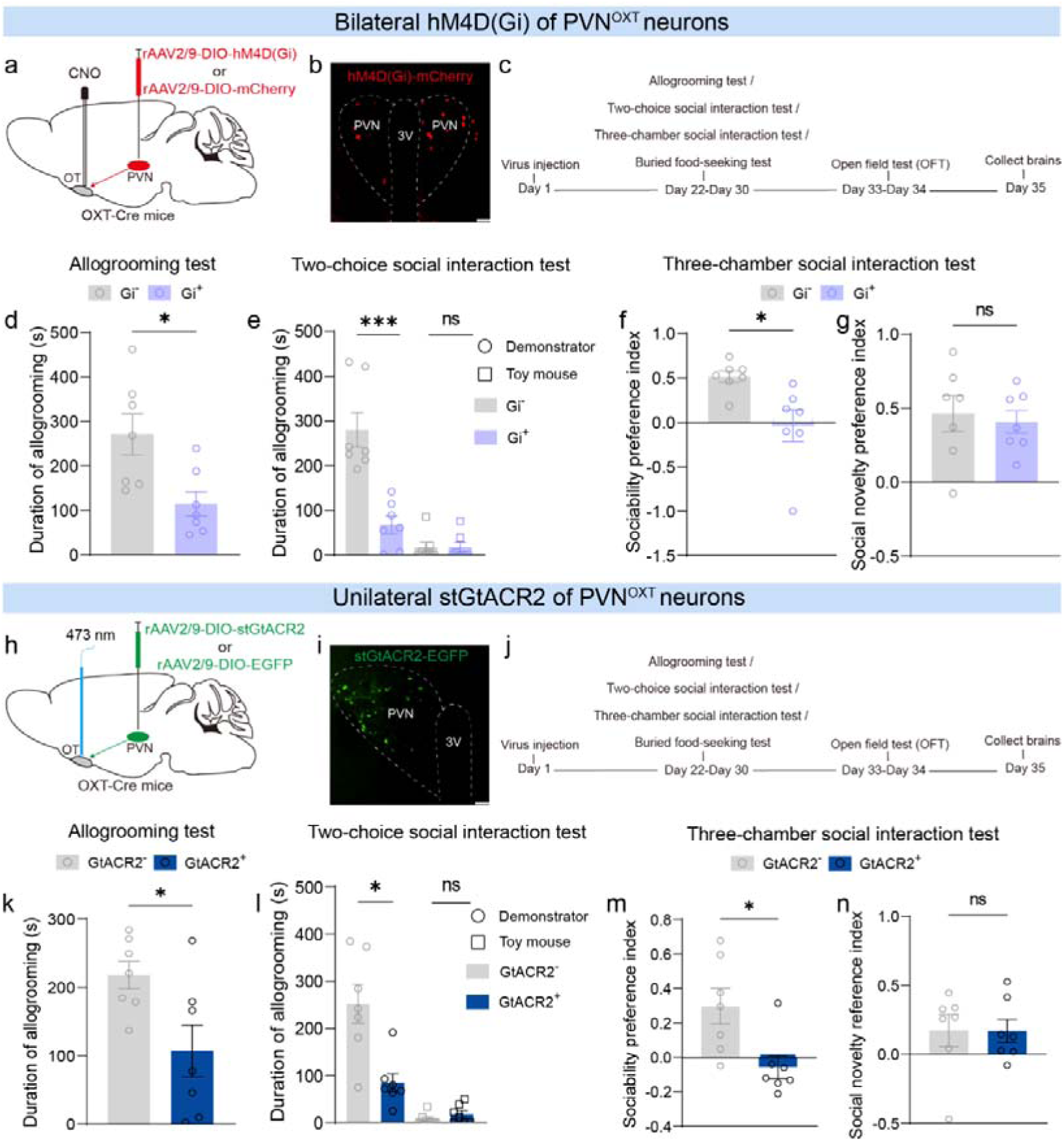
Inhibition of the PVN^OXT^-OT pathway reduces observer’s allogrooming toward anesthetized demonstrator. **a** Chemogenetic viral strategy schematic. **b** Representative image showing hM4Di-expressing OXT neurons in the PVN. **c** Behavioral assay timeline. **d-g** Chemogenetic inhibition of PVN^OXT^ neuronal fibers in the OT ameliorated allogrooming (**d**; allogrooming test), decreased biased allogrooming toward demonstrator (**e**; two-choice test), impaired social recognition and had no effect on social novelty (**f-g**; three-chamber test). **h** Optogenetic viral strategy schematic. **i** Representative image showing stGtACR2-expressing OXT neurons in the PVN. **j** Behavioral assay timeline. **k-n** Optogenetic inhibition of PVN^OXT^ neuronal fibers in OT ameliorated allogrooming (**k**; allogrooming test), decreased biased allogrooming toward demonstrator (**l**; two-choice test), impaired sociability preference but had no effect on social novelty preference (**m-n**; three-chamber test). Chemogenetic: observers (n = 7; **d-g**) and demonstrators (n = 7; **d-e**). Optogenetic: observers (n = 7; **k-n**) and demonstrators (n = 7; **k-l**). 3V, third ventricle. Data are presented as mean ± SEM. \**p*□<□0.05; ns, no significance. Statistical details see Table S1.

We next used optogenetics to recapitulate these findings. We bilaterally expressed the Cre-dependent stGtACR2 or control virus in the PVN and unilaterally implanted (due to spatial limit) an optic fiber in the mOT of OXT-Cre mice (**Fig. 3h-i**). In the same behavioral assays (**Fig. 3j**), inhibition of the PVN^OXT^-OT pathway via stGtACR2 similarly decreased allogrooming (**Fig. 3k-l**) and, interestingly, also reduced sociability preference without affecting social novelty (Fig. **3m-n**). Inhibition did not alter non-social behaviors such as locomotion or olfactory investigation (**Fig. S4**). To determine pathway specificity, we inhibited the PVN^OXT^ projections to the anterior olfactory nucleus (AON), an OXTR-expressing region^57,58,73^ and anatomically receiving projections from PVN^OXT^ neurons^44^. This manipulation did not affect allogrooming, social novelty, olfactory perception, or locomotion, though it reduced sociability, confirming a specific role for the PVN^OXT^-OT pathway in allogrooming (**Fig. S5**).

We reasoned that PVN^OXT^-OT pathway activation increases allogrooming via OXT release in the mOT. Supporting this, bilateral OXT infusion into the mOT significantly increased allogrooming time (**Fig. S6**). We then chemogenetically activated the pathway by bilaterally expressing the Cre-dependent excitatory DREADDs (hM3Dq) or control virus in the PVN and implanting cannula in the mOT (**Fig. 4a-b**). In behavioral assays (**Fig. 4c**), chemogenetic activation via CNO in hM3Dq-expressing observers significantly increased allogrooming during a direct social test (**Fig. 4d**). In a two-choice test, this activation selectively enhanced time spent allogrooming an anesthetized demonstrator versus a toy (**Fig. 4e**), while sociability and social novelty preference remained unchanged (**Fig. 4f-g**). Parallel optogenetic activation experiments yielded similar results (**Fig. 4h-n**), and no changes in non-social behaviors were observed (**Fig. S7**). In conclusion, the PVN^OXT^-OT pathway plays a facilitatory role in modulating allogrooming toward anesthetized demonstrators.

**Figure 4.**
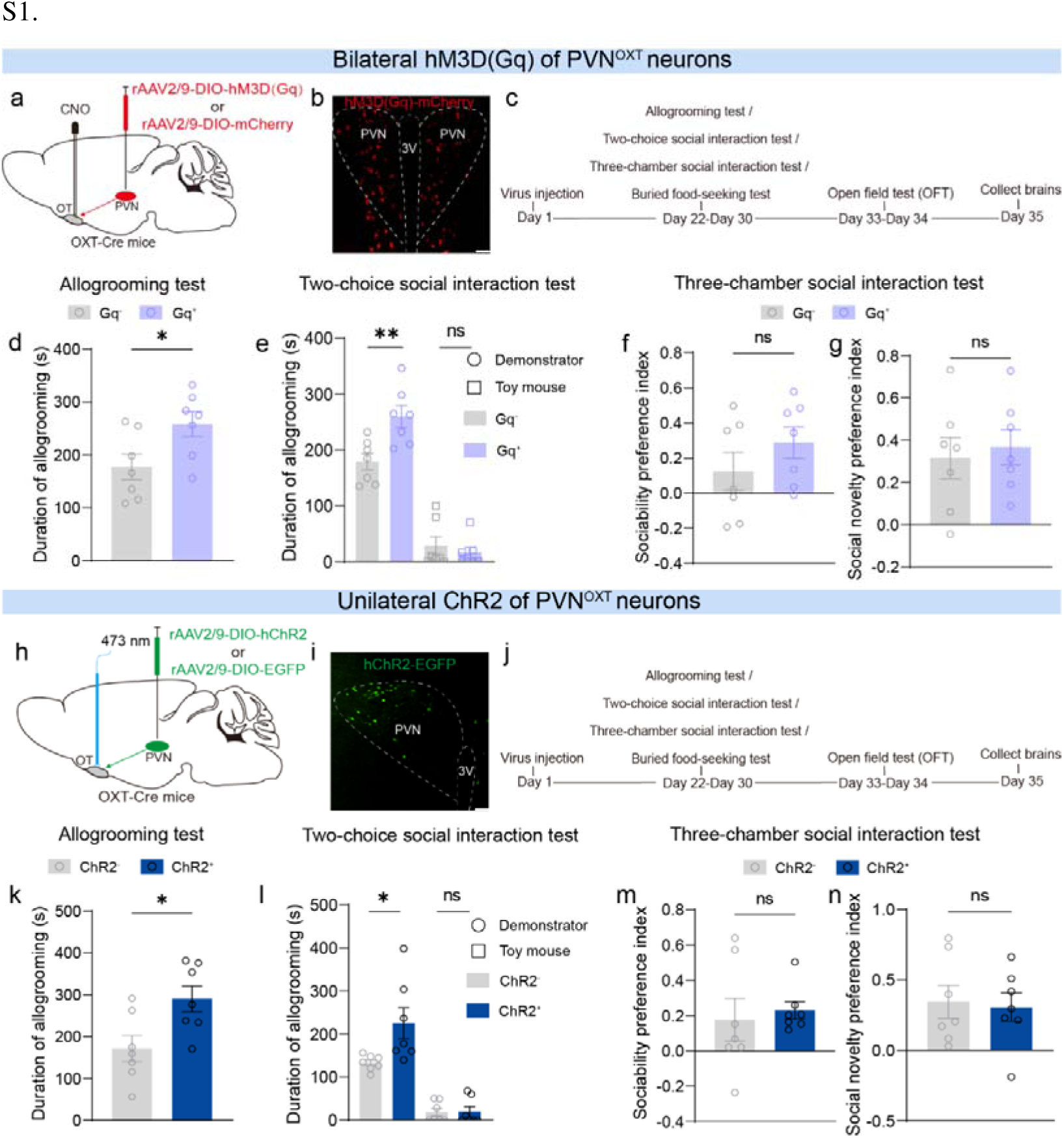
Activation of the PVN^OXT^-OT pathway increases observer’s allogrooming toward anesthetized demonstrator. **a** Chemogenetic viral strategy schematic. **b** Representative image showing hM3Dq-expressing OXT neurons in the PVN. **c** Behavioral assay timeline. **d-g** Chemogenetic activation of PVN^OXT^ neuronal fibers in OT increased allogrooming (**d**; allogrooming test), enhanced biased allogrooming toward demonstrator (**e**; two-choice test), but had no effect on social recognition and novelty (**f-g**; three-chamber test). **h** Optogenetic viral strategy schematic. **i** Representative image showing ChR2-expressing OXT neurons in the PVN. **j** Behavioral assay timeline. **k-n** Optogenetic activation of PVN^OXT^ neuronal fibers in OT increased allogrooming (**k**; allogrooming test), enhanced biased allogrooming toward demonstrator (**l**; two-choice test), but had no effect on sociability and social novelty preference (**m-n**; three-chamber test). Chemogenetic: observers (n = 7; **d-g**) and demonstrators (n = 7; **d-e**). Optogenetic: observers (n = 7; **k-n**) and demonstrators (n = 7; **k-l**). 3V, third ventricle. Data are presented as mean ± SEM. \**p*□<□0.05; ns, no significance. Statistical details see Table S1.

### PVN^OXT^-OT pathway activity and oxytocin release are time-locked to allogrooming

To examine how the PVN^OXT^-OT pathway responds to social interaction, particularly allogrooming directed toward anesthetized demonstrators, we performed fiber photometry of calcium dynamics in axon terminals. We expressed the axonal calcium indicator GCaMP6s or control EGFP in the mOT of OXT-Cre mice (**Fig. S8a-b**), enabling monitoring of activity in PVN^OXT^ axon terminals during behavioral interactions (**Fig. S8c**). GCaMP6s signals in PVN^OXT^ terminals increased robustly and time-locked to the onset of allogrooming and social investigation of an anesthetized demonstrator, reflected in increased area under the curve (AUC) and peak ΔF/F following behavior onset (**Fig. S8d-e**). No significant responses occurred during non-social behaviors (rearing and walking; **Fig. S8f-g**) or during any interaction with a toy mouse (**Fig. S8h-k**). Control EGFP recordings showed no fluorescence changes (**Fig. S9**). These results demonstrate that mOT-projecting PVN^OXT^ neurons are specifically activated during social behaviors, with the strongest responses during allogrooming.

We next asked whether this activity drives oxytocin release in the mOT. We expressed the OXT sensor OXT1.0 or EGFP control in the mOT of wild-type mice (**Fig. 5a-b**). After three weeks, we recorded OXT-dependent sensor dynamics during behaviors (**Fig. 5c**). We observed a rapid, time-locked increase in OXT sensor signal, reflected by the larger AUC and higher peak Δ F/F upon initiation of allogrooming or investigation of anesthetized demonstrators (**Fig. 5d-e**). No significant release was detected during non-social behaviors (rearing and walking), interactions with a toy mouse (**Fig. 5f-k**), or non-contact investigation (**Fig. S10**). Control EGFP mice showed no signal changes (**Fig. S11**). Together, these results show that PVN^OXT^ terminal activity and subsequent OXT release in the mOT are specifically and temporally coupled to active social interaction, most strongly during allogrooming.

**Figure 5.**
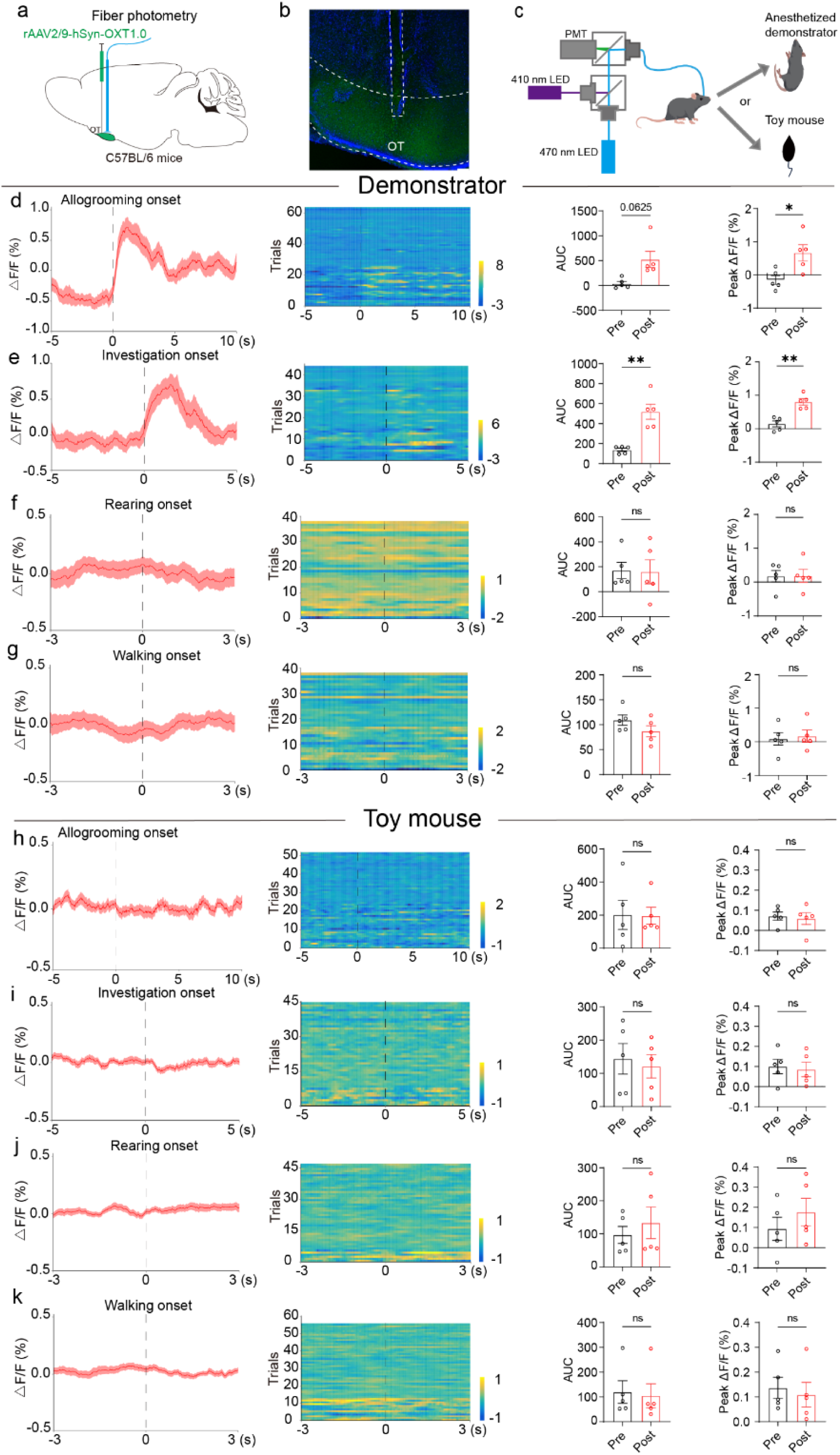
OXT release in the OT is temporally locked to observer’s allogrooming or social investigation initiation of anesthetized demonstrator. **a** Schematic illustrating virus injection strategy. **b** Represent image showing OXT sensor expression and fiber implantation sites in the OT. Scale bar: 100□μm. **c** Schematic of fiber photometry recordings from an observer during interacting with demonstrator or toy mouse. **d-g** Peri-event averaged OXT changes traces (left), heatmaps of individual trial OXT changes in the OT (middle), and quantifications of OXT-dependent fluorescence changes (far right panels; AUC, area under the curve; peak ΔF/F; pre, -3-0 s; post, 0-3 s) during observer’s allogrooming (**d**) and investigation (**e**) toward demonstrator, as well as rearing (**f**) and walking (**g**). **h-k** Corresponding analyses for observer interaction with a toy mouse (**h-k** mirror **d-g**, respectively). Total trials: 63, 44, 39, 38, 53, 44, 47, and 55 for **d-k**, respectively (7-15 trials/mouse from five C57BL/6 observer mice). Dashed lines indicate behavior onset, and shaded areas denote SEM. The same cohort of demonstrators (C57BL/6 mice, n = 5; **d-g**) were used. Data are presented as mean ± SEM. \**p*□<□0.05, \*\**p*□<□0.01; ns, no significance. Statistical details see Table S1.

### OXTR signaling in the OT is necessary for allogrooming toward anesthetized conspecifics

The OXTR signaling is essential for social behaviors^31^. We therefore hypothesized that the PVN^OXT^-OT pathway modulates alloogroming via OXTR activation in the OT. To test this, we first infused the OXTR antagonist L□368,899 or saline (control) bilaterally into the mOT of wild□type mice prior to behavioral testing (**Fig. S12a-b**). Antagonizing OXTR significantly reduced the time observers spent allogrooming anesthetized demonstrators in both a direct social test (**Fig. S12c**) and a two-choice test, without affecting interactions with a toy mouse (**Fig. S12d**). This treatment did not alter sociability or social novelty preference (**Fig. S12e-f**).

To validate these findings genetically, we next generated OT□specific Oxtr conditional knockout (Oxtr□cKO) mice by bilaterally expressing Cre□EGFP or EGFP control in the mOT of Oxtr□floxed mice (**Fig. S12g**). After three weeks for viral expression and recombination (**Fig. S13m**), behavioral assays were performed (**Fig. S12h**). Mirroring the pharmacological results, compared to Oxtr-floxed observers, Oxtr□cKO observers showed markedly reduced allogrooming of anesthetized demonstrators in a direct test (**Fig. S12i**). In the two□choice test, allogrooming was shifted from the demonstrator toward the toy mouse (**Fig. S12j**). Oxtr□cKO also impaired sociability but spared social novelty preference (**Fig. S12k-l**). Both pharmacological and genetic blockade of Oxtr in the OT did not affect olfactory perception, although genetic knockout induced anxiety□like behaviors (**Fig. S13a-l**). Together, these loss□of□function experiments demonstrate that OXTR signaling within the OT is necessary for allogrooming directed toward anesthetized conspecifics, effectively recapitulating the behavioral phenotype observed after inhibition of the upstream PVN^OXT^-OT pathway.

### OXTR signaling in OT D1 neurons is required for allogrooming toward anesthetized conspecifics

The OT consists largely of GABAergic neurons expressing dopamine D1 or D2 receptors (D1 or D2 neurons), with sparse interneurons including D3□receptor-expressing neurons (D3 neurons) in the islands of Calleja (IC)^47–50^. Given that the OT receives oxytocinergic input from the PVN^26,44,54,55^, and OXT fibers and OXTRs are present in the IC^24,56^ and broader OT^57,58^, we examined whether OXTR signaling in specific dopaminergic cell types regulates allogrooming.

We conditionally deleted Oxtr in D1- or D2-expressing neurons by injecting D1-or D2-Cre-expressing AAVs into the mOT of Oxtr-floxed mice (generating D1^OxtrDcKO^ or D2^OxtrDcKO^; **Fig. 6a, i**). Behavioral tests three weeks post-injection (**Fig. 6b, j**) revealed that D1^Oxtr-cKO^ observers showed markedly reduced allogrooming of anesthetized demonstrators (**Fig. 6c**), without significant changes in sociability, social novelty, olfaction, or locomotion (**Fig. 6d-h**). In contrast, D2^OxtrDcKO^ observers displayed no behavioral alterations (**Fig. 6k-p**). Because D3 neurons co-express D1 receptors particularly in the IC of the OT in rodents^74,75^, we also selectively ablated D1□expressing neurons within the D3 population; this manipulation did not impair allogrooming or social preference, though it reduced locomotion (**Fig. S14**). Thus, OXTR signaling specifically in D1 neurons, but not in D2 or D3 neurons, is necessary for allogrooming toward anesthetized conspecifics.

**Figure 6.**
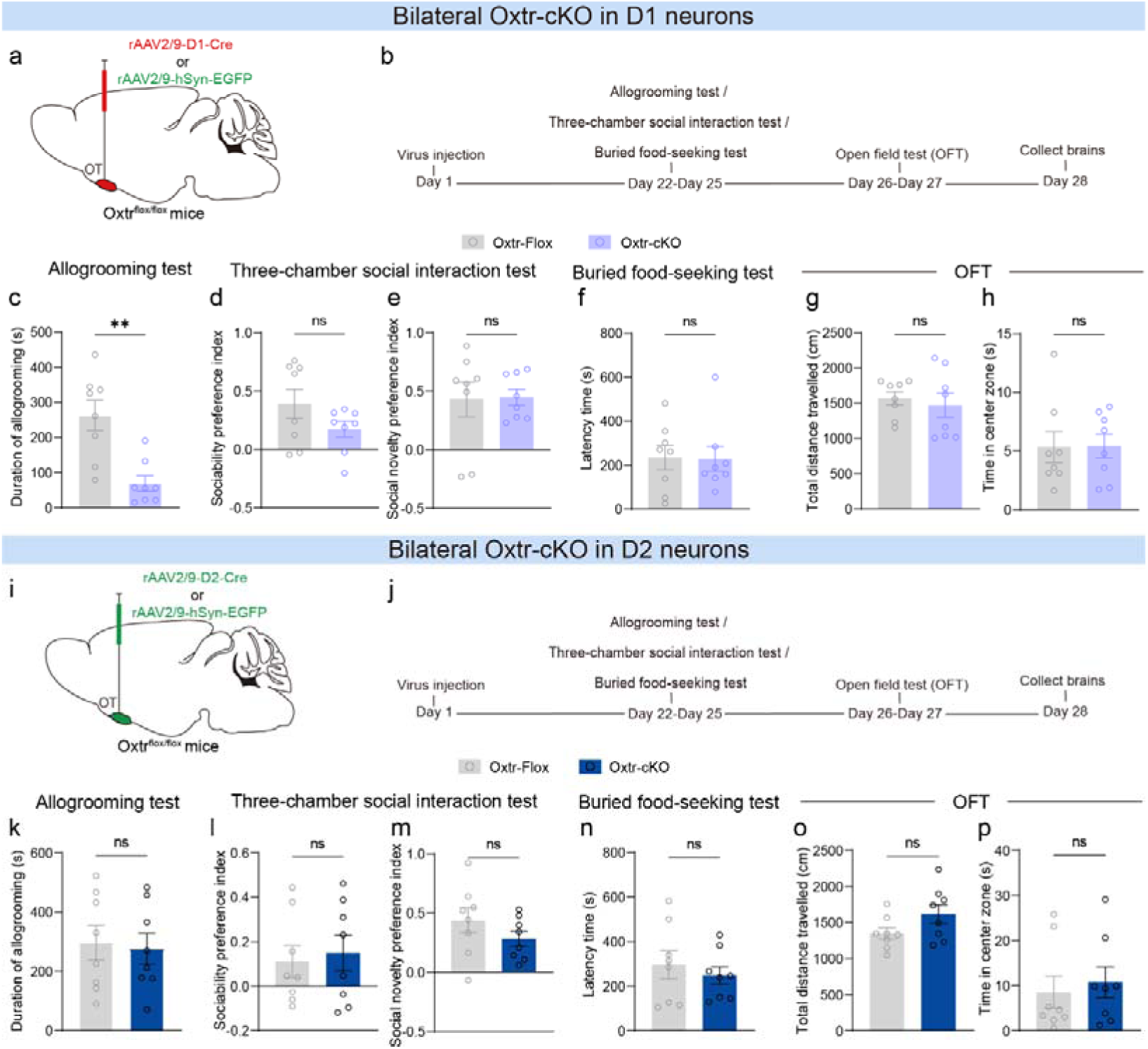
Oxtr-expressing D1 neurons are required for observer’s allogrooming toward anesthetized demonstrator. **a** Schematic of viral injection strategy. **b** Behavioral assay timeline. **c-h** Bilateral Oxtr-cKO in D1 neurons ameliorated allogrooming (**c**; allogrooming test), but did not affect social recognition and social novelty (**d-e**; three-chamber test), and had no effect on olfactory perception (**f**; buried food-seeking test) and locomotion (**g-h**; open field test, OFT). **i** Schematic of viral injection strategy. **j** Behavioral assay timeline. **k-p** Bilateral Oxtr-cKO in D2 neurons had no effect on allogrooming (**k**; allogrooming test), social recognition and novelty (**l-m**; three-chamber test), olfactory perception (**n**; buried food-seeking test), and locomotion (**o-p**; OFT). In **c-h**, observers: Oxtr-Flox controls (n = 8), Oxtr-Flox controls with AAV-D1-Cre (n = 8); Demonstrators C57BL/6 (n = 8). In **k-p**, Oxtr-Flox controls (n = 8), Oxtr-Flox controls with AAV-D2-Cre (n = 8); Demonstrators C57BL/6 (n = 8). Data are presented as mean ± SEM. \**p*□<□0.05, \*\**p*□<□0.01; ns, no significance. Statistical details see Table S1.

To understand how OXTR signaling regulates D1 neuron activity, we performed patch-clamp recordings in acute OT slices from D1□tdTomato mice (**Fig. S15a**). OXT application suppressed D1 neuron intrinsic excitability (reduced firing frequency) without altering resting membrane potential (RMP) (**Fig. S15b-d**). OXT also increased the frequency of spontaneous excitatory postsynaptic currents (sEPSCs) and the amplitude of spontaneous inhibitory postsynaptic currents (sIPSCs) (**Fig. S15e-j**), suggesting enhanced excitatory input and strengthened inhibitory postsynaptic responses. Recordings from D1^OxtrDcKO^ neurons revealed hyperexcitability, including depolarized resting potential, increased firing, and reduced sIPSC amplitude (**Fig. S16**). By contrast, neither OXT application nor Oxtr deletion in D2 neurons significantly altered their electrophysiological properties (**Fig. S15k-, S16k-t**). Additionally, in vivo fiber photometry confirmed that D1 neurons in the OT showed increased activity before the initiation of allogrooming and social investigation of an anesthetized demonstrator but not other non-social behaviors such as walking (**Fig. S17-18**). Together, these data indicate that OXT acts through OXTRs on OT D1 neurons to suppress their excitability, providing a cellular mechanism by which PVN-derived oxytocinergic signaling modulates allogrooming behavior.

### Overexpression of GIRK channels normalizes increased neuronal excitability and rescues allogrooming deficit in D1^Oxtr-cKO^ mice

OXTR signaling can modulate G protein-gated inwardly rectifying K□ (GIRK/Kir3) channels^76^, which are abundant in the rodent OT^77^ and inhibit neurons via potassium efflux and membrane hyperpolarization. To test whether augmenting GIRK activity could counteract the effects of OXTR loss, we bilaterally overexpressed D1-Cre-dependent GIRK channels (Kir3.2) in OT D1 neurons of Oxtr-floxed mice using a D1-Cre driver virus (**Fig. 7a-b**). Three weeks later, GIRK overexpression (GIRK□OE) in D1^Oxtr-cKO^ neurons significantly increased allogrooming toward anesthetized conspecifics (**Fig. 7c**) without affecting social recognition or novelty preference (**Fig. 7d-e**), indicating a specific role for GIRK in rescuing D1^OxtrDcKO^-induced allogrooming deficit. Electrophysiological recordings confirmed that this behavioral rescue corresponded to normalized neuronal excitability, as GIRK□OE reduced both resting membrane potential and firing frequency (**Fig. 7f-h**). These results demonstrate that enhancing GIRK-mediated inhibition directly reverses the neuronal hyperexcitability and allogrooming deficit in D1^Oxtr-cKO^ mice, suggesting a potentially causal link between OXTR signaling, GIRK channel function, and this prosocial behavior.

**Figure 7.**
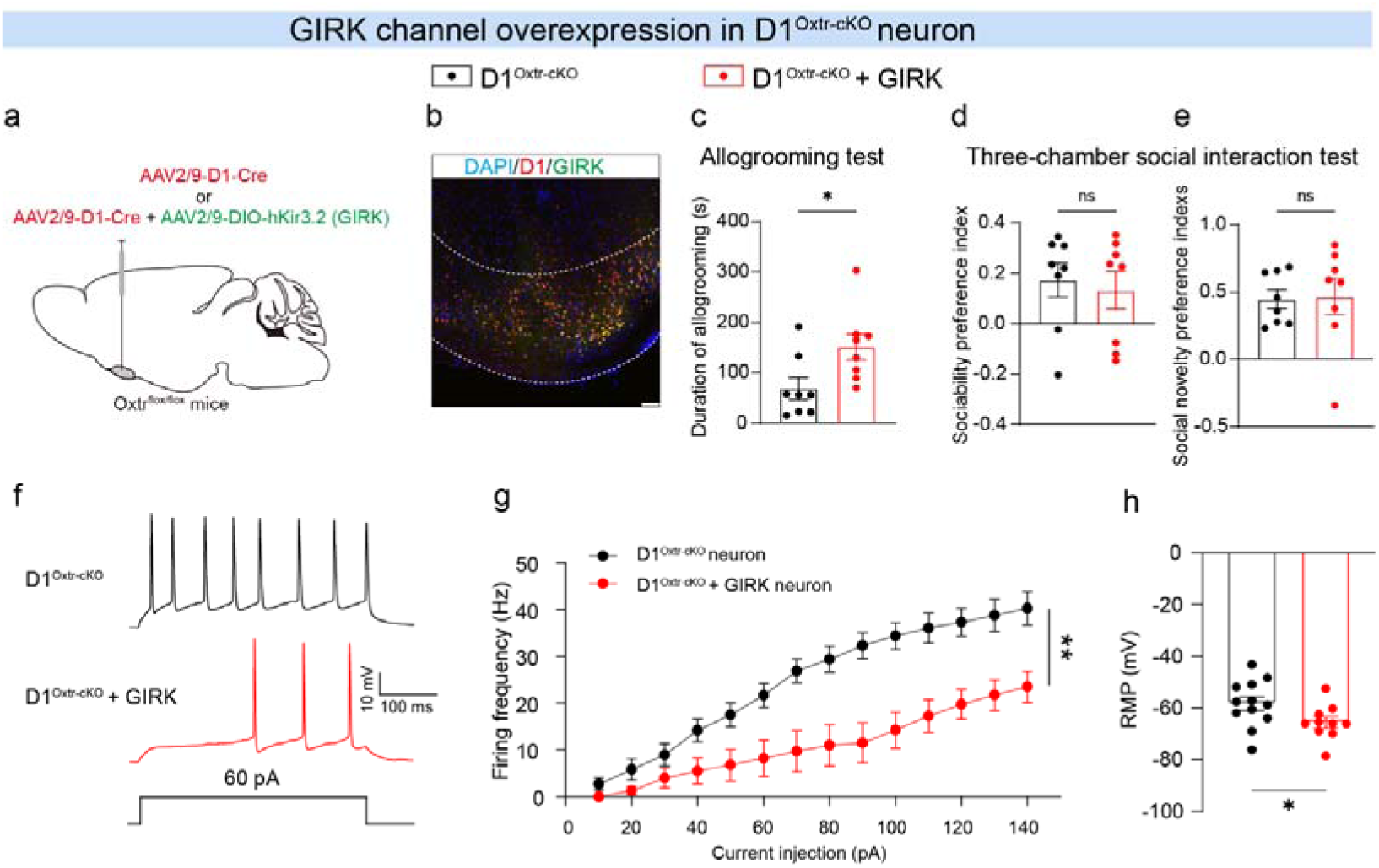
Overexpression of GIRK (GIRK OE) rescues neuronal hyperactivity and allogrooming deficits in D1^Oxtr-cKO^ mice. **a** Schematic of viral injection strategy. **b** Representative image showing GIRK expression in OT D1 neurons. Scale bar, 200 μm. **c-e** Bilateral GIRK OE in D1^Oxtr-cKO^ neurons increased allogrooming (**c**) but did not alter performance in the social recognition or social novelty phases of the three-chamber test (**d-e**). **f-h** Representative action potential traces from a D1^Oxtr-cKO^ neuron and a D1^Oxtr-cKO^ neuron with GIRK OE (**f**). GIRK OE normalized the elevated firing frequency (**g**) and hyperpolarized the resting membrane potential (RMP; **h**) in D1^Oxtr-cKO^ neurons. For behavior, D1^Oxtr-cKO^: n = 8 mice; D1^Oxtr-cKO^ + GIRK OE: n = 7 mice. For patch clamp recording, D1^Oxtr-cKO^ neurons: n = 12 cells; D1^Oxtr-cKO^ + GIRK OE neurons: n = 10 cells (from 3 mice per group). The D1^Oxtr-cKO^ datasets in **c-e** and **g** are reproduced from Figure 6 and Figure S16, respectively. Data are presented as mean ± SEM. \**p*□<□0.05, \*\**p*□<□0.01; ns, no significance. Statistical details see Table S1.

**Figure 8.**
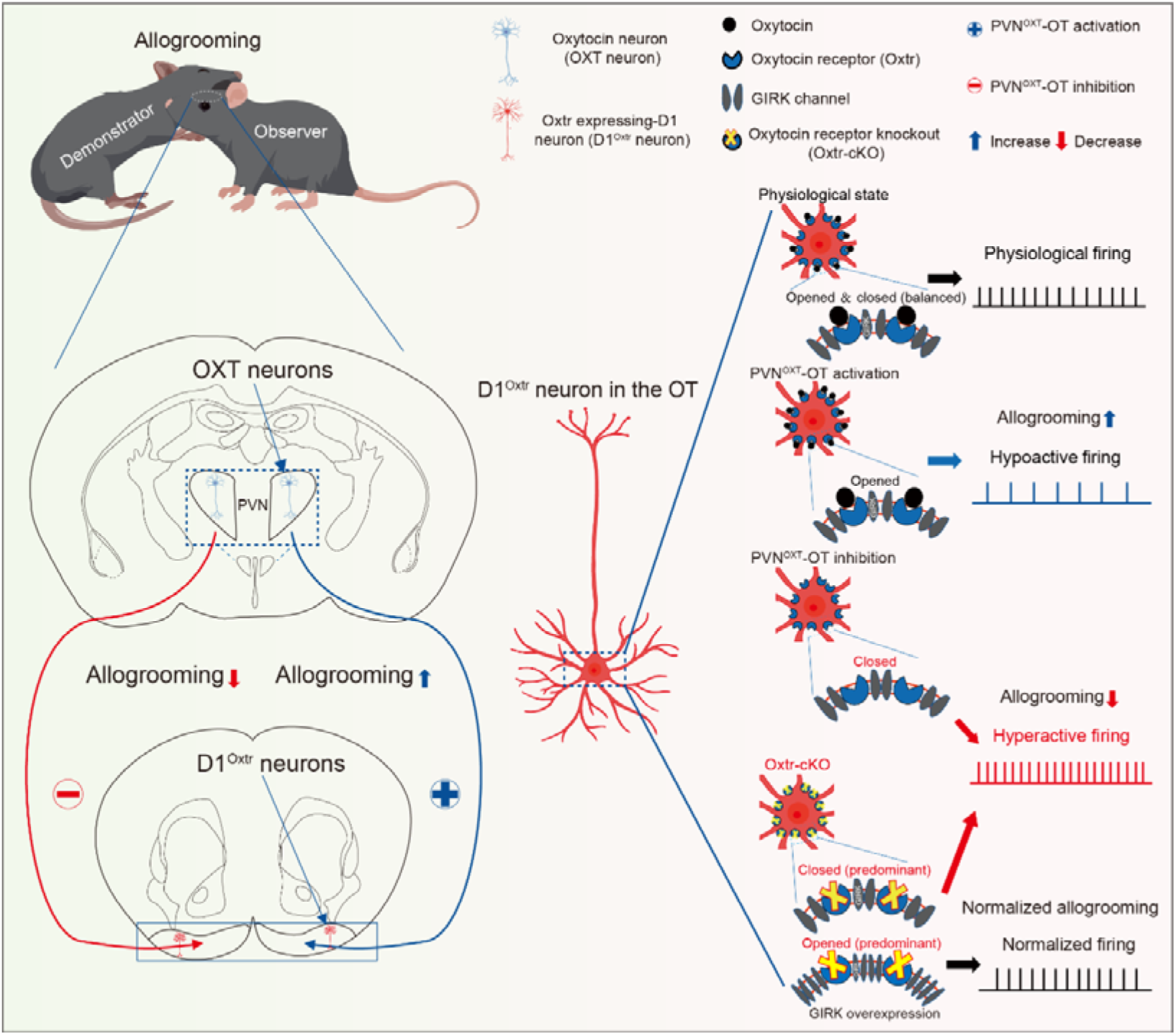
Proposed model: the PVN^OXT^-OT pathway bidirectionally modulates allogrooming via GIRK channels in D1Oxtr neurons. This model posits that oxytocin (OXT) release from the paraventricular nucleus (PVN^OXT^) to the olfactory tubercle (OT) orchestrates allogrooming by regulating GIRK channels on OT dopamine receptor 1-expressing neurons that contain the OXT receptor (D1^Oxtr^). Inhibition of this pathway decreases, while activation increases, allogrooming toward anesthetized conspecifics. OXTR signaling within OT D1 neurons is essential for this process. Under physiological conditions, OXT binding to OXTR maintains tonic GIRK activity via intracellular signaling, ensuring appropriate D1 neuron firing and baseline allogrooming. Pathway activation increases synaptic OXT, enhancing GIRK opening, suppressing D1 neuron activity (hypoactive firing), and promoting allogrooming. Conversely, pathway inhibition reduces OXT availability, decreasing GIRK activity, leading to D1 neuron hyperexcitability and impaired allogrooming. Conditional Oxtr knockout in D1 neurons (D1^Oxtr-cKO^) ablates this regulation, causing reduced GIRK activity, neuronal hyperexcitability, and allogrooming deficits. Overexpression of GIRK channels in the D1^Oxtr-cKO^ background restores normal GIRK function, rescuing both the aberrant neuronal firing and the allogrooming impairment.

## Discussion

Allogrooming, as one subtype of rescue-like behavior or reviving-like prosocial behavior, is evolutionarily conserved across species^4,8–14^. However, the neural substrates responsible for regulating allogrooming still remain incompletely understood. Herein, taking advantage of the sodium pentobarbital-induced anesthesia mouse model to mimic mouse’s unconscious and unresponsive state and by virtue of optogenetic, chemogenetic, and pharmacological manipulations, genetic ablation, electrophysiological recordings, as well as behavioral assays, we interrogated the role of PVN^OXT^-OT pathway in the orchestration of allogrooming. It denoted that observer exhibited substantial allogrooming toward the anesthetized demonstrators, which could effectively mitigate anesthesia-induced anxiety-like behaviors in the demonstrators (**Figs. 1, S1**). Moreover, the main olfactory system was critically essential for the observer’s biased allogrooming toward an anesthetized demonstrator versus an awake one (**Fig. 2**). Loss-of-function of the PVN^OXT^-OT pathway reduced, while activation of it increased, observer’s allogrooming toward the anesthetized demonstrator (**Figs. 3-4**). OXTR signaling that involved G protein-gated inwardly rectifying K^+^ (GIRK) channels in D1 neurons in the OT was indispensable for this process (**Figs. 6-7, S12, S15-17**). Further fiber photometry recordings demonstrated that the PVN^OXT^-OT pathway as well as OXT in the OT showed time-locked allogrooming-related activities/changes (**Figs. 5, S8**). Collectively, the current study indicated a critical and bidirectional role of the PVN-OT oxytocinergic pathway in modulating allogrooming. Our study provides more evidence for neural underpinnings of OXT’s role in modulating allogrooming in mice and deepens the understanding of oxytocin as a prosocial neuropeptide during social interactions in mice.

Allogrooming confers significant emotional benefits to recipients across multiple vertebrate species, reducing stress and tension in both mammalian and avian taxa^78–80^. Our result showed that observer’s allogrooming mitigated demonstrator’s distress state (anxiety-like behaviors) (**Fig. S1**). Similarly, a recent study reveals that observer’s rescue-like behaviors including allogrooming promotes the demonstrator’s awakening^4^. This cross-species evidence supports the notion that allogrooming serves a conserved, stress-buffering function, and are and ecologically meaningful. Intriguingly, in our study, observer, instead of showing any obvious distressed phenotypes, exhibited hyperactivity upon sensing the anesthetized state of the demonstrator regardless of physical contact or not. This is in contrast to a finding that observer became distress when encountering anesthetized conspecifics and reduced its own stress levels following allogrooming its distressed conspecifics^4^. This discrepancy is potentially reasonable due to different mice species recruited and distinct experimental settings.

The cooperation of different sensory systems to perceive and process corresponding cues guarantees the fulfillment of appropriate behaviors^81^, including allogrooming, among animals, however, information regarding which sensory cues contribute to allogrooming, is still incomplete^4,8,10^. Our study indicates that observer spatially preferred an anesthetized demonstrator over an awake one, regardless of visual cues (**Fig. 2a-d**), suggesting visual cues are not indispensable for observer’s allogrooming. Notably, this biased spatial preference was not detected in methimazole-treated observers, and these observers also exhibited significantly reduced allogrooming toward the anesthetized demonstrator compared to saline-treated observers (**Fig. 2e-j**). These evidence denotes that the main olfactory system responsible for olfactory cues processing is necessary during observer’s allogrooming toward anesthetized demonstrator. Herein, the bidirectional manipulating the PVN^OXT^-OT pathway could modulate allogrooming via OXT signaling in the OT, a key component of the main olfactory system (**Figs. 3-5, S8, S12**). Moreover, the endogenous OXT system contributes greatly and is crucial in the modulation of many social behaviors that depend on sensory cues including olfaction^34,82,83^. It is likely that the PVN^OXT^-OT pathway-controlled OXT changes in the OT influence olfactory cues processing, which eventually affects observer’s biased allogrooming toward anesthetized peers.

A notable dissociation emerged from our circuit manipulations: inhibition of the PVN^OXT^-OT pathway reduced both allogrooming and sociability preference (**Fig. 3**), whereas activation of the same pathway selectively enhanced allogrooming without affecting sociability (**Fig. 4**). This functional dissociation is intriguing and suggests that the PVN^OXT^-OT pathway may exert distinct, context-dependent influences on social behavior. One possible interpretation is that allogrooming and sociability preference, though both prosocial in nature, engage separable circuit mechanisms. Allogrooming represents a targeted, action-oriented form of prosocial interaction that requires precise motor coordination and sensory integration, whereas sociability preference reflects a more general motivational drive to approach and investigate a conspecific. The observation that chemogenetic inhibition of PVN^OXT^-OT projections to the anterior AON reduced sociability but spared allogrooming (**Fig. S5**) further supports the idea that these two behavioral domains are mediated by distinct, yet overlapping, oxytocinergic pathways. This is consistent with the emerging view that OXT circuits are organized in a projection-specific manner, with different terminal fields governing distinct aspects of social behavior^4,31,36^. For instance, PVN^OXT^ projections to the central amygdala (CeA) and dorsal bed nucleus of the stria terminalis (dBNST) have been shown to separately coordinate the emotional and motor components of rescue-like behavior^4^. The bidirectional modulation we observed, where pathway activation selectively enhanced allogrooming but not sociability, may reflect a ceiling effect or, more interestingly, suggest that the pathway primarily encodes the execution of allogrooming once social motivation is already engaged. Conversely, the reduction in sociability following pathway inhibition may indicate that this circuit contributes to the motivational salience of social cues under certain conditions, a role that may be dispensable when allogrooming is pharmacologically or optogenetically driven. Collectively, these findings highlight the functional heterogeneity of the PVN^OXT^-OT pathway and underscore the importance of considering both the direction of circuit manipulation and the specific behavioral readout when interpreting oxytocinergic regulation of prosocial behavior. Future experiments employing projection-specific and activity-dependent labeling, combined with real-time behavioral tracking, will be necessary to dissect the precise contributions of this pathway to distinct facets of social interaction.

A particularly striking observation emerged from our loss-of-function experiments: OXTR ablation in the OT did not merely reduce allogrooming but instead shifted the observer’s allogrooming behavior from the anesthetized conspecific toward the toy mouse (**Fig. S12j**). This redirection of prosocial behavior is intriguing and suggests that OXTR signaling in the OT is not simply permissive for allogrooming but rather plays a critical role in guiding the target specificity of this behavior. One interpretation is that OXTR in the OT may be essential for the proper processing or valuation of social olfactory cues, enabling the observer to distinguish between a distressed conspecific and an inanimate object. Without this signaling, the motivational salience of social cues may be diminished, causing the observer to redirect grooming behavior toward an alternative, non-social target that nonetheless permits the expression of the motor program. This interpretation aligns with our finding that the main olfactory system is necessary for selective allogrooming toward anesthetized conspecifics (**Fig. 2**), and with evidence that OXT signaling modulates olfactory processing in other brain regions to guide social behavior^42,53,83^. Alternatively, the shift toward the toy mouse could reflect a disruption in the balance between social and non-social reward valuation. Given that the OT is a key node in the ventral striatum, a region critical for reward processing, OXTR deletion in this area may impair the assignment of reward value to social stimuli, leading to a compensatory increase in engagement with a non-social object that retains its incentive salience. This interpretation is consistent with studies showing that OXT signaling in the ventral striatum promotes social reward and facilitates social approach^84–86^, and that disruption of oxytocinergic modulation can lead to a relative preference for non-social stimuli^31,38^. Notably, this behavioral shift was specific to allogrooming, as OXTR ablation did not significantly alter social investigation or sociability preference in the same paradigm (**Fig. S12j**), suggesting that the effect is not simply due to a general reduction in social motivation. Rather, it points to a more nuanced role for OT OXTR signaling in orchestrating the appropriate allocation of specific prosocial actions toward their intended targets. Collectively, these findings underscore the importance of oxytocinergic modulation within the OT for ensuring that allogrooming, a behavior with critical adaptive value, is directed toward conspecifics in need, rather than toward irrelevant objects.

An interesting dissociation emerged from our loss-of-function experiments: while pharmacological antagonism of OXTR in the OT did not alter anxiety-like behavior (**Fig. S12e-f**), genetic deletion of OXTR in the same region induced significant anxiety-like phenotypes, as reflected by reduced center exploration in the open field test and increased time spent in the dark compartment in the light-dark transition test (**Fig. S13a-l**). This discrepancy between acute receptor blockade and chronic genetic ablation is noteworthy and warrants consideration. One plausible explanation is that the two manipulations operate on different timescales and engage distinct compensatory mechanisms. Acute antagonism with L□368899 likely provides a transient and partial blockade of OXTR signaling, sufficient to disrupt allogrooming but insufficient to induce lasting changes in emotional state. In contrast, constitutive OXTR deletion, even when restricted to the OT, may trigger developmental or homeostatic adaptations that extend beyond the acute loss of receptor function. For instance, chronic OXTR deficiency could lead to compensatory alterations in other neurotransmitter systems (e.g., dopamine, serotonin, or GABA) that modulate anxiety-related circuitry, or it may impair the development of stress-regulatory pathways that are normally shaped by oxytocinergic signaling during development^87–89^. Additionally, the OT itself has been implicated in emotional processing beyond its role in olfaction, and chronic disruption of OXTR signaling within this region may unmask latent contributions to anxiety regulation that are not apparent following acute blockade^57,58^. Notably, both pharmacological and genetic manipulations similarly impaired allogrooming and sociability, suggesting that these social behaviors are more directly dependent on ongoing OXTR signaling, whereas the effects on anxiety may require more sustained disruption to manifest. This dissociation highlights the importance of considering both the duration and mechanism of manipulation when interpreting oxytocinergic contributions to distinct behavioral domains. Future studies employing temporally controlled, inducible knockout strategies could help dissociate developmental from acute effects of OXTR loss in the OT on emotional regulation.

Herein, we observed that OXTR signaling specifically in D1 neurons, but not in D2 or D3 neurons, of the OT is required for allogrooming toward anesthetized conspecifics, highlighting the crucial role of OT D1 neurons in regulating social behavior. A recent study demonstrated that repetitive D1 receptor inhibition in the dorsomedial striatum (DMS) significantly improved valproic acid (VPA)-induced social deficits but not repetitive behaviors, whereas repetitive D2 receptor inhibition alleviated VPA-induced repetitive behaviors without affecting social deficits, suggesting distinct behavioral roles: D1-expressing neurons predominantly mediate social behaviors, while D2-expressing neurons are more involved in asocial behaviors^90^. Further evidence underscored the importance of D1 receptor-expressing neurons in the ventral striatum (including the NAc) in orchestrating social behaviors. For instance, D1, but not D2, neurons in the NAc shell drove isolation-induced sociability^91^. Additionally, both D1 receptors and OXTR have been implicated in responses to social loss in female prairie voles^86^, suggesting that the specific involvement of OXTR-expressing D1 neurons in allogrooming may stem from intrinsic differences in intracellular signaling between D1- and D2-like dopamine receptors upon OXT-OXTR interactions. Together, these findings emphasize the critical and selective role of D1 neurons, particularly those expressing OXTR, in modulating social behaviors such as allogrooming.

While our study identifies the PVN^OXT^-OT pathway as a novel neural substrate responsible for modulating allogrooming, several caveats warrant consideration. First, only male mice were recruited in this study. This decision was made both to adhere to animal welfare principles by minimizing animal use and to avoid potential confounds from hormonal fluctuations during the female estrous cycle. This approach is supported by the axonal projections of PVN^OXT^ neurons in mice do not exhibit obvious sex differences^44,45,84^. Furthermore, recent studies on allogrooming as a prosocial behavior suggest minimal sexual dimorphism; for instance, the amount of head grooming performed by observers was comparable regardless of the sex of either the demonstrator or the observer^10^, and sex combinations played only a minor role in regulating allogrooming toward unfamiliar peers^8^. However, it is important to acknowledge the broader literature documenting sex differences in the oxytocinergic regulation of social behavior. For example, sex-specific patterns of OXTR expression and signaling have been reported in the central amygdala and other limbic regions, contributing to divergent behavioral outcomes in stress and social contexts^85,92^. Moreover, while allogrooming itself may show minimal sex differences, the underlying neural circuit mechanisms, including the role of the PVN^OXT^-OT pathway, could be modulated by sex hormones that influence OXT neuron activity and OXTR function^87,89^. Given these well-documented nuances in oxytocin signaling, it remains to be established whether the specific circuit mechanism we describe-involving the PVN^OXT^-OT pathway and OXTR signaling on D1 neurons-is fully generalizable to female mice. Future studies incorporating both sexes will be essential to determine if this pathway operates similarly or employs distinct mechanisms to regulate allogrooming and other prosocial behaviors across the sexes. Second, OXT regulated social behavior through multiple brain regions beyond the OT, such as the ventral tegmental area (VTA) and nucleus accumbens (NAc)^88^. For instance, OXT release in the VTA orchestrated dopamine neuron activity to promote social reward ^93,94^, whereas interactions between OXT and dopamine signaling in the NAc modulated pair bonding in prairie voles^95^, and OXT-mediated serotonin release in the NAc promoted social reward in mice^96^. Given that the OT contains abundant dopamine receptors (e.g., D1, D2, D3) that receives dopaminergic inputs from regions such as the VTA, a key question remains whether OXTR signaling within the OT interacts with dopamine systems, or other neurotransmitters or neuropeptides, to regulate allogrooming, a mechanism that awaits further investigation.

## Acknowledgements

This work was supported by the National Natural Science Foundation of China (Grant No. 82371515), the National Key R&D Program of China (2024YFD1400500), the Initiative Scientific Research Program, Institute of Zoology, CAS (Grant No. 2025IOZ08), the State Key Laboratory of Animal Biodiversity Conservation and Integrated Pest Management (Grant No. SKLA2505), the Talent Initiation BaiRen Plan Start-up Funds (Grant No. E251F811), the State Key Laboratory of Integrated Management of Pest Insects and Rodents (Grant No. IPM2301), the Initiative Scientific Research Program, Institute of Zoology, CAS (2024IOZ0105), and the Open Project of Hebei Key Laboratory of Early Life Health Promotion (ELHP-KF202501).

## Author Contributions

Conceptualization, J.M., Y.Y, C.Y.-L. and Y.F.-Z.; Methodology and investigation, all authors; Formal analysis, data curation, and visualization, J.Y., Y.Z., Y.Q. and L.G.; Writing, J.Y., Y.Z., Y.Q., L.G., M.M, J.M., Y.Y, C.Y.-L. and Y.F.-Z.; Supervision and funding acquisition, Y.F.-Z.

## Supplementary Figures

**Figure S1.**
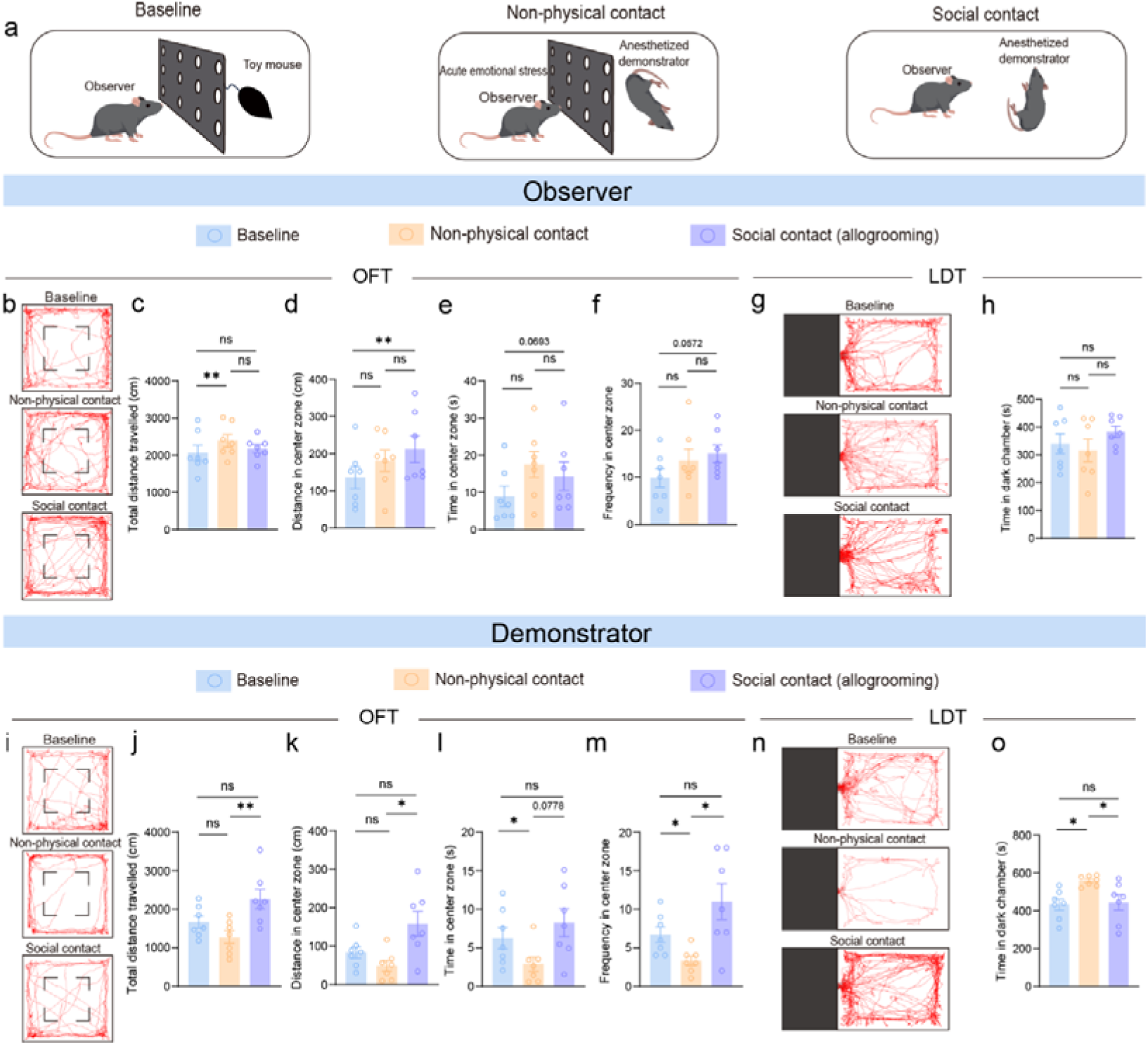
Observer-initiated allogrooming mitigates anesthesia-induced anxiety-like behaviors in demonstrator. **Related to Figure 1. a** Schematic of non-physical/social contact behavioral paradigm. **b-h** Observer behavior in open filed test (OFT) (**b-f**) and light-dark box transition test (LDT) (**g-h**) following interaction with demonstrators. **b, g** Representative locomotion traces in OFT (**b**) and LDT (**g**). **c** Total distance travelled. **d-f** Center zone activity: distance travelled (**d**), duration (**e**), and entry frequency (**f**). **h** Duration in dark chamber (LDT). **i-o** Demonstrator behavior in OFT (**i-m**) and LDT (**n-o**) following allogrooming by observers. **i, n** Locomotion traces (OFT: **i**; LDT: **n**). **j** Total distance travelled. **k-m** Center zone activity: distance (**k**), duration (**l**), entry frequency (**m**). **o** Duration in dark chamber. Data from consistent cohorts (observers: n = 7 [**b–h**]; demonstrators: n = 7 [**i–o**]). Data are presented as mean ± SEM. \**p*□<□0.05, \*\**p*<□0.01; ns, no significance. Statistical details see Table S1.

**Figure S2.**
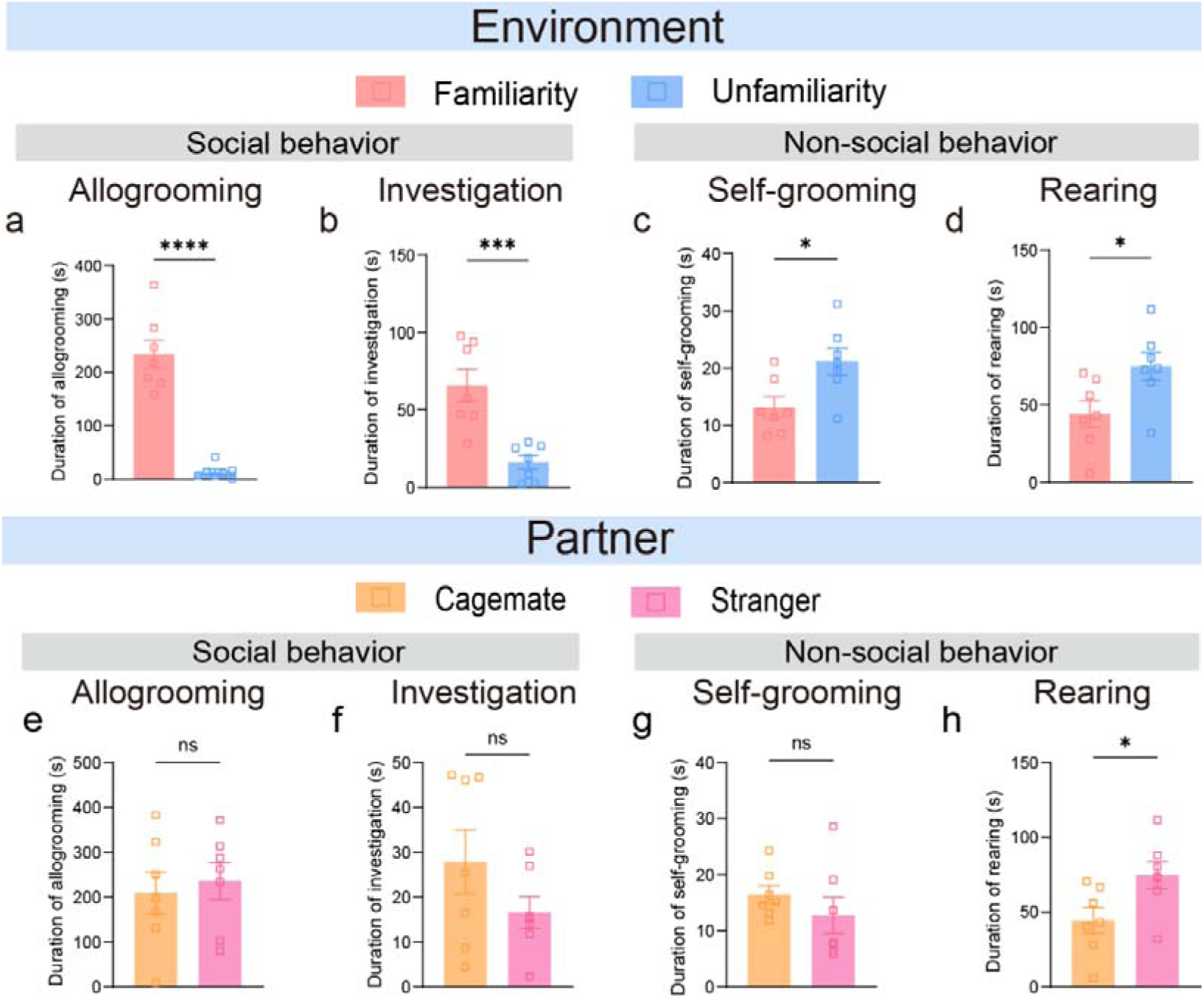
Familiarity with the environment, but not with the partner, influences observer’s allogrooming toward anesthetized demonstrator. **Related to Figure 1. a-d** Observer behavior in familiar versus unfamiliar environments. **a, b** Social behavior duration: allogrooming (**a**) and social investigation (**b**). **c, d** Non-social behavior duration: self-grooming (**c**) and rearing (**d**). **e-h** Observer behavior toward familiar versus unfamiliar partners. **e, f** Social behavior toward cagemate versus stranger: allogrooming (**e**) and social investigation (**f**). **g, h** Non-social behavior duration: self-grooming (g) and rearing (**h**). Observers: n = 7 (all panels). Demonstrators: cagemates (n = 7), strangers (n = 7). Data are presented as mean ± SEM. \**p*□<□0.05, \*\*\**p*□<□0.001, \*\*\*\**p*<□0.0001; ns, no significance. Statistical details see Table S1.

**Figure S3.**
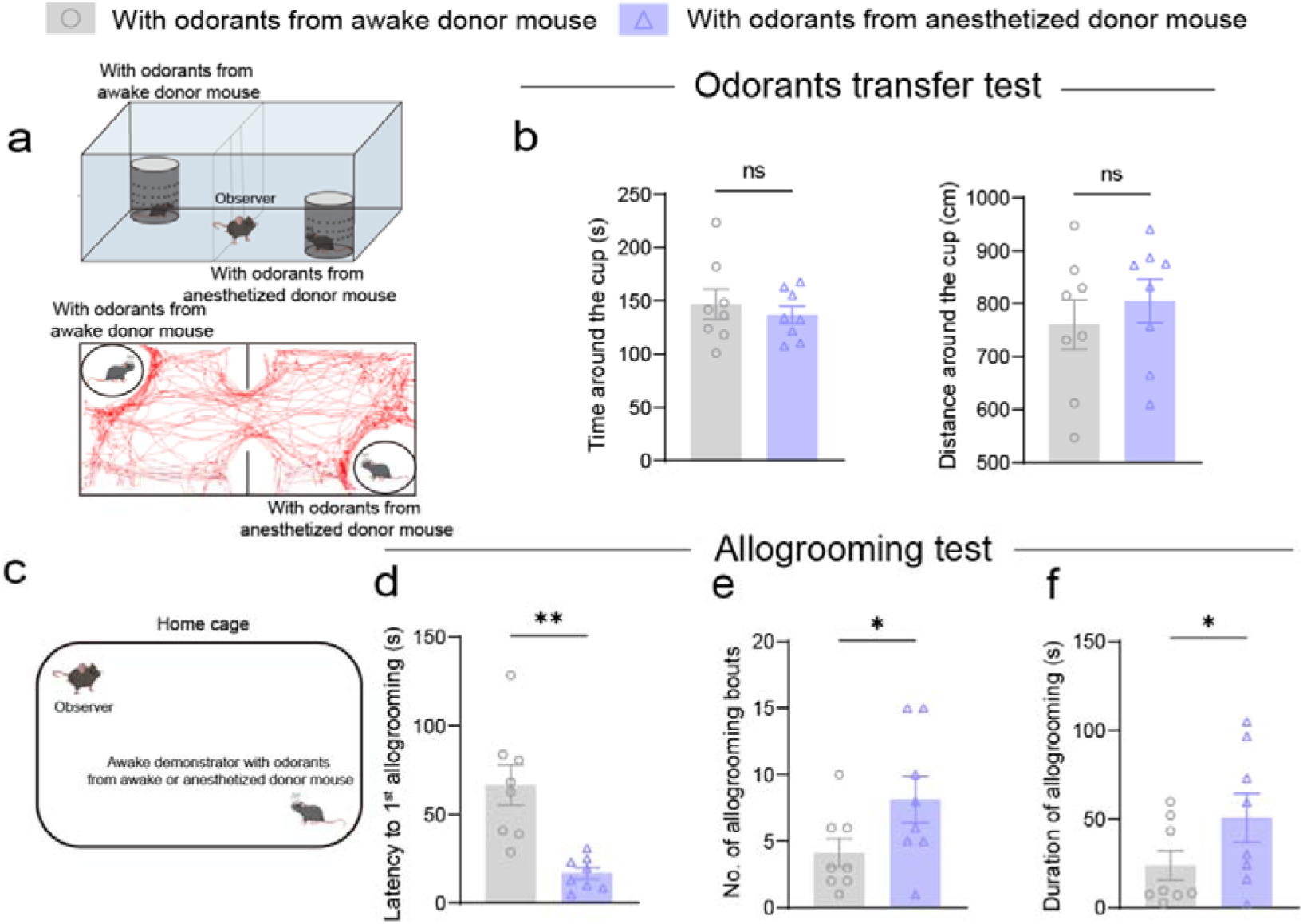
Observer directs preferential allogrooming toward unfamiliar conspecifics bearing odorants derived from anesthetized partner. **Related to Figure 2. a** Behavioral paradigm schematic (top) and representative observer locomotion tracks (bottom). **b** Duration of exploration (left) and distance travelled (right) around the stimulus cup. **c** Allogrooming test schematic. **d-f** Allogrooming metrics: latency to first event (**d**), event frequency (e) and total duration (**f**). Same cohort of observers (n = 8) and demonstrators (n = 16) in **b** and **d-f**. Data are presented as mean ± SEM. \**p*□<□0.05, \*\**p*<□0.01; ns, no significance. Statistical details see Table S1.

**Figure S4.**
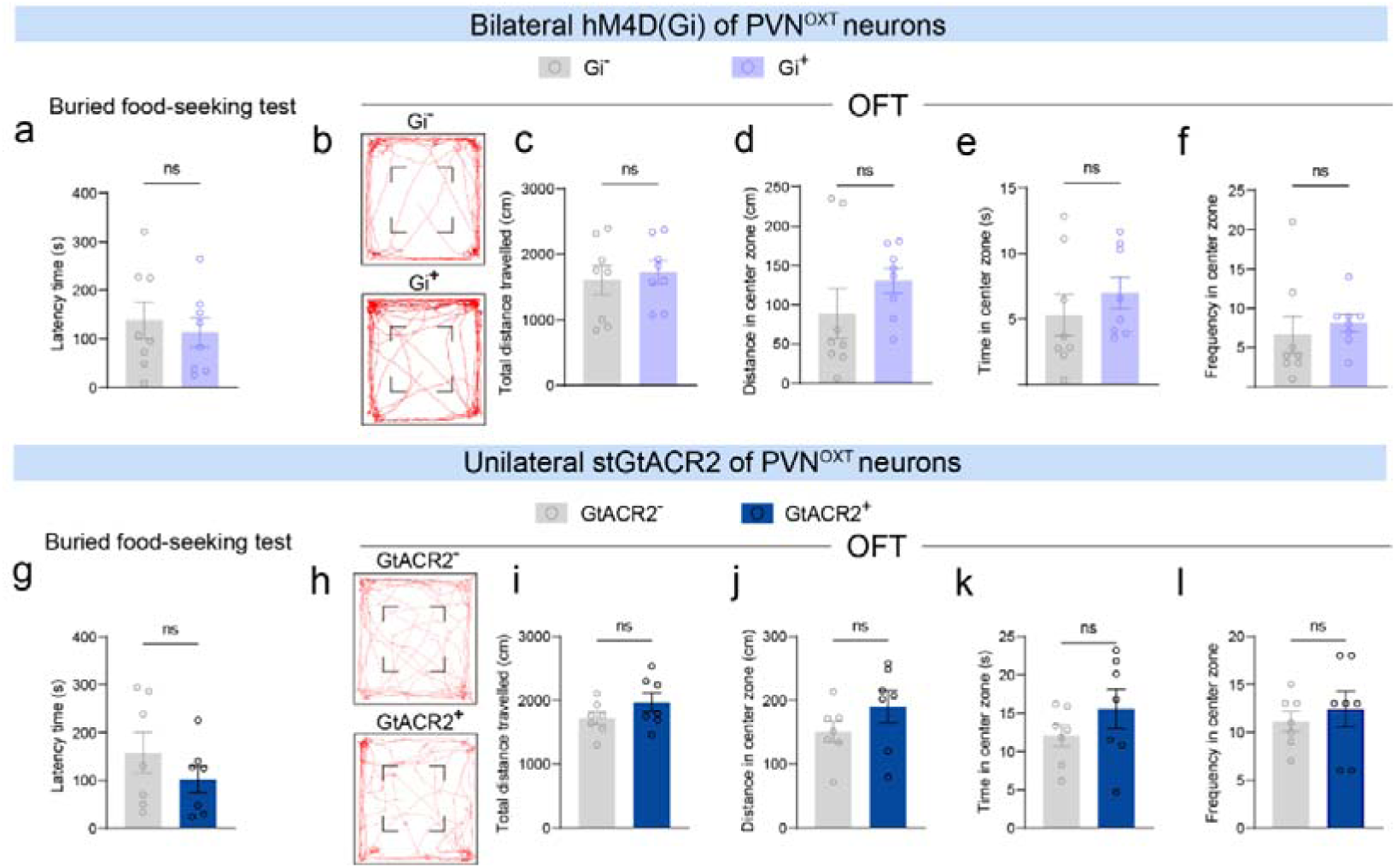
Inhibition of the PVN^OXT^-OT pathway does not alter olfactory perception or locomotion in observer. **Related to Figure 3.** Behavioral performance of observer following chemogenetic (**a-f**) or optogenetic (**g-l**) inhibition of the PVN^OXT^-OT pathway. **a, g** Latency to find the first food pellet in the buried food-seeking test. **b-f, h-l** Behavioral parameters measured in the open field test (OFT). **b, h** Representative locomotor tracks. **c, i** Total distance travelled. **d, j** Distance travelled in the center zone. **e, k** Time spent in the center zone. **f, l** Frequency of entries into the center zone. Same observers (n = 8) in **a-f** and another cohort of observers (n = 7) in **g-l**. ns, no significance. Statistical details see Table S1.

**Figure S5.**
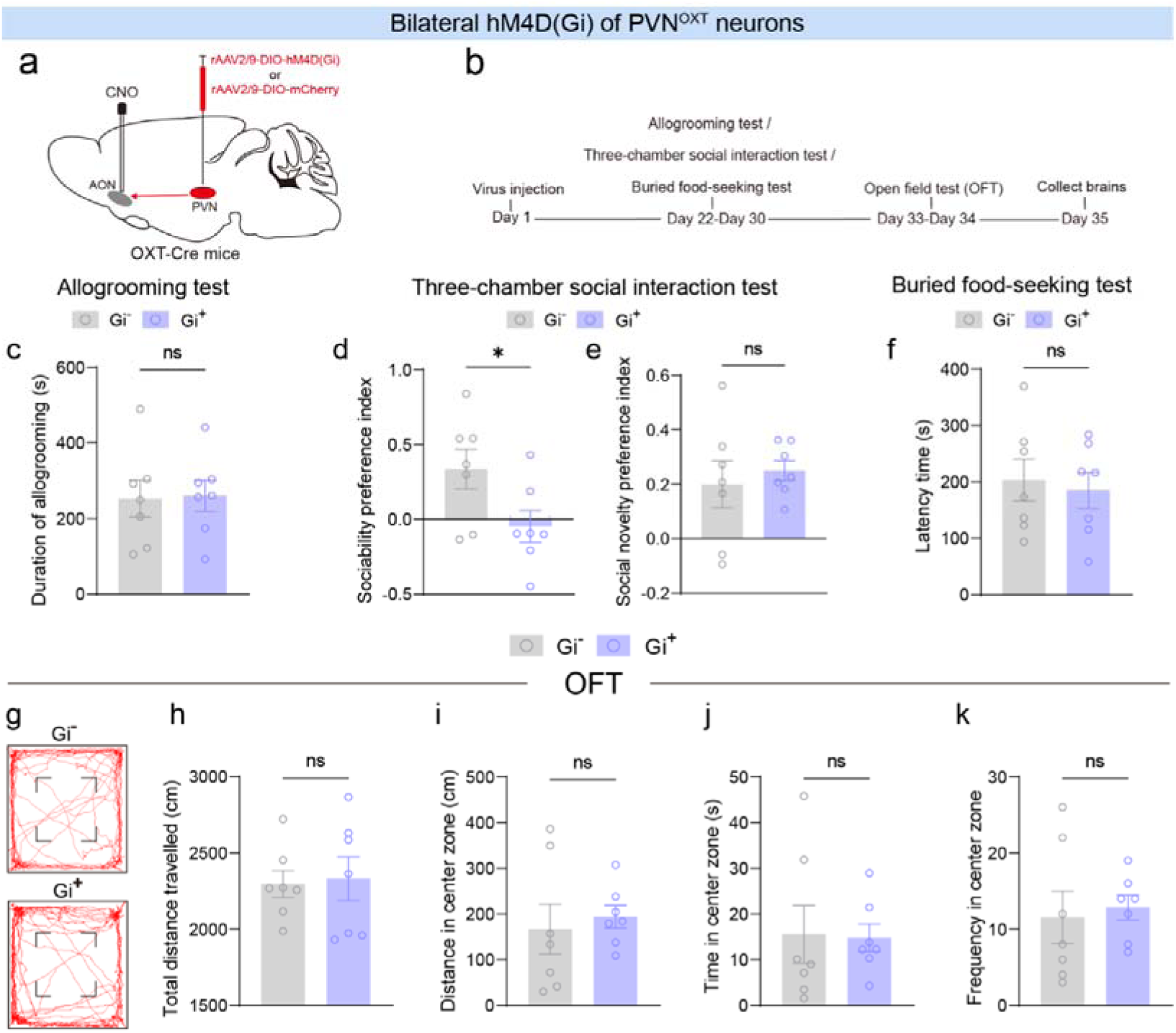
Chemeogenetic inhibition of the PVN^OXT^-AON pathway does not alter observer’s allogrooming toward anesthetized demonstrator. **Related to Figure 3. a** Schematic of viral injection strategy. **b** Timeline of behavioral assays. **c-f, h-i** Inhibition of PVN^OXT^ neuronal fibers in the anterior olfactory nucleus (AON) had no effect on observer allogrooming in the allogrooming test (**c**), olfactory perception (**f**) in the buried food-seeking test, or locomotion in the open field test (OFT) (**h, i**). However, it impaired sociability preference (**d**) without affecting social novelty preference (**e**) in the three-chamber social interaction test. Locomotor tracks (**g**), center zone time (**j**), and center zone entry frequency (**k**) in the OFT were unaffected by the PVN^OXT^-AON pathway inhibition. The same cohort of observers (n = 7) were tested in **c-k**. Demonstrators (n = 7) were used in **c**. Data are presented as mean ± SEM. \**p*□<□0.05; ns, no significance. Statistical details see Table S1.

**Figure S6.**
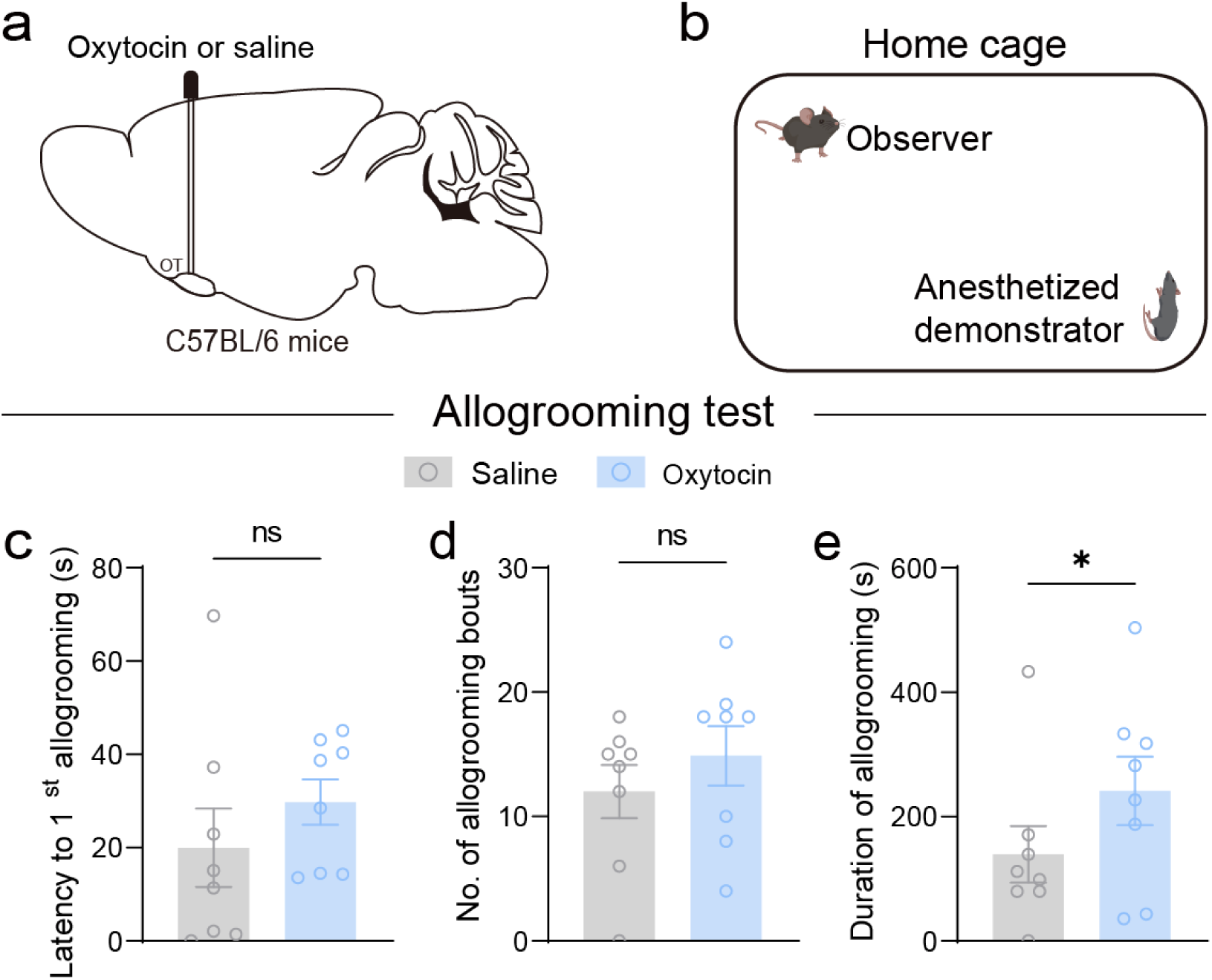
Bilateral oxytocin administration into the OT enhances observer’s allogrooming toward anesthetized demonstrator. **Related to Figure 4. a** Cannula implantation schematic and pharmacological strategy. **b** Behavioral paradigm schematic. **c-e** Allogrooming metrics: latency to first event (**c**), event frequency (**d**) and total duration (**e**). Same cohort of observers (n = 8) and demonstrators (n = 8) in **c-e**. Data are presented as mean ± SEM. \**p*□<□0.05; ns, no significance. Statistical details see Table S1.

**Figure S7.**
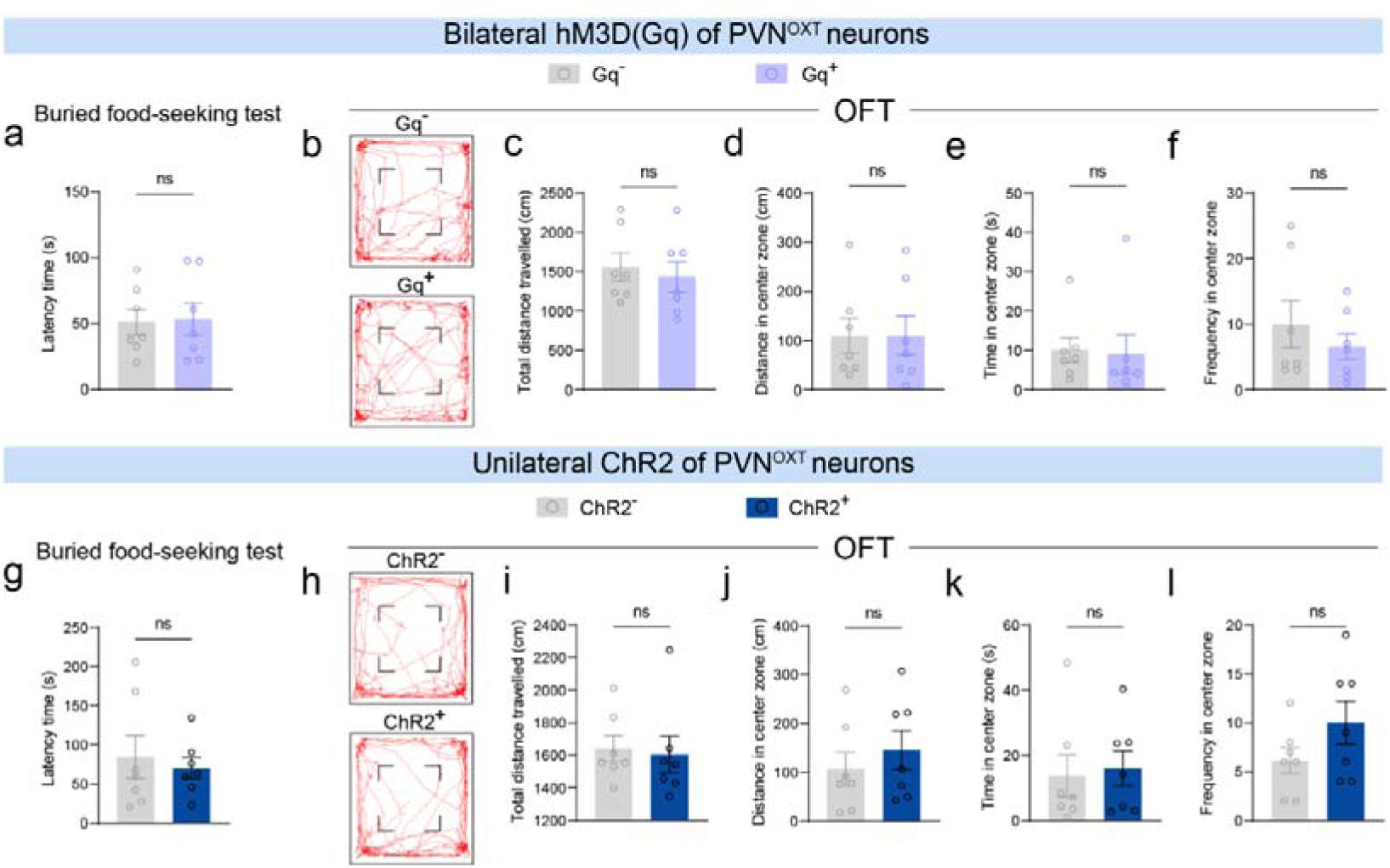
Activation of the PVN^OXT^-OT pathway does not alter olfactory perception or locomotion in observer. **Related to Figure 4.** Behavioral performance of observer following chemogenetic (**a-f**) or optogenetic (**g-l**) activation of PVN^OXT^-OT pathway. **a, g** Latency to find the first food pellet in the buried food-seeking test. **b-f, h-l** Behavioral parameters measured in the open field test (OFT). **b, h** Representative locomotor tracks. **c, i** Total distance travelled. **d, j** Distance travelled in the center zone. **e, k** Time spent in the center zone. **f, l** Frequency of entries into the center zone. Same observers (n = 7) in **a-f** and another cohort of observers (n = 7) in **g-l**. ns, no significance. Statistical details see Table S1.

**Figure S8.**
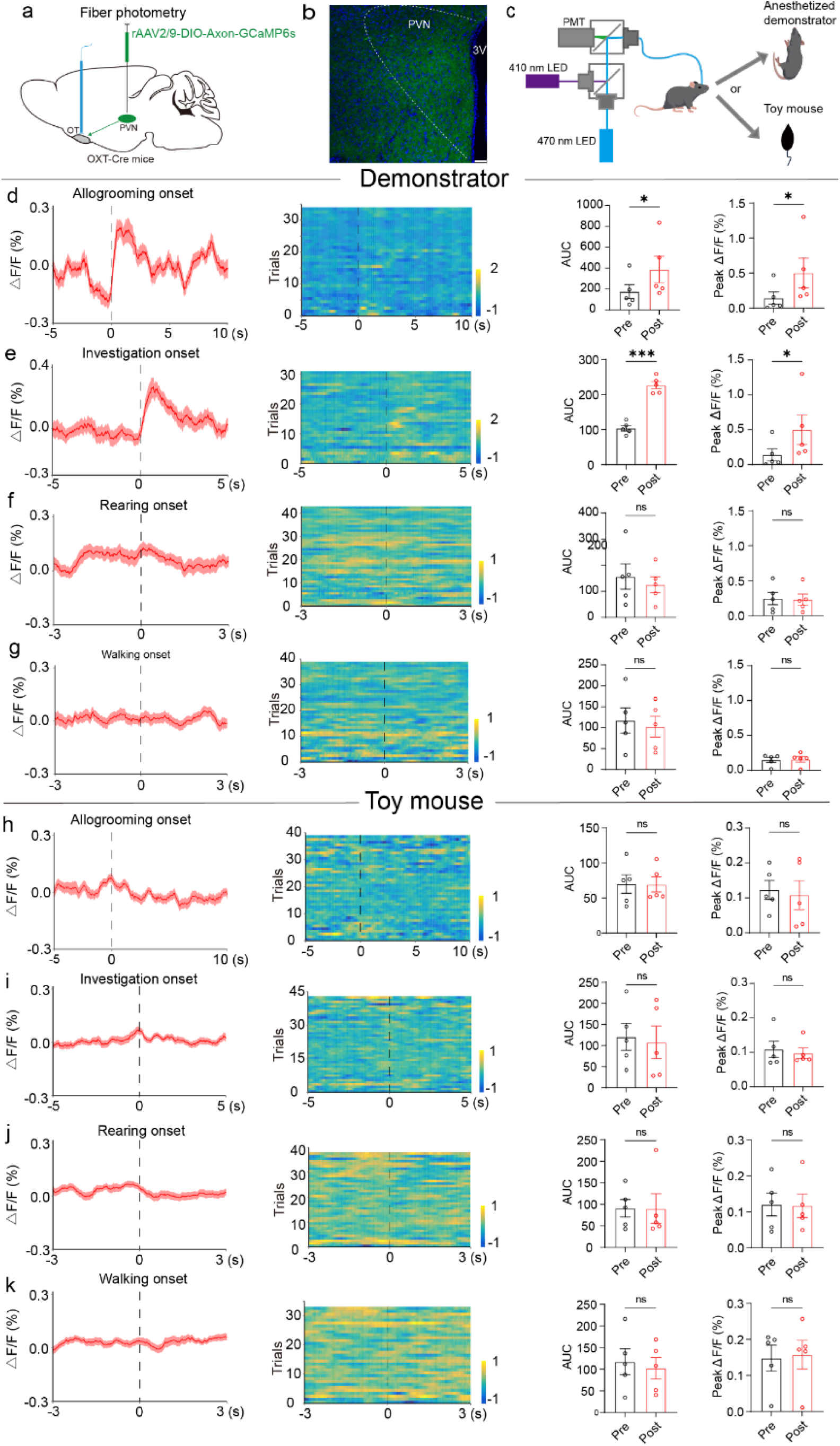
PVN^OXT^-OT neuronal fibers exhibit increased time-locked activity during observer’s allogrooming or social investigation of anesthetized demonstrator. **Related to Figure 3, 4. a** Schematic illustrating viral injection strategy. **b** Represent images showing GCaMP6s expression in the PVN. Scale bars: 100□μm. **c** Schematic of fiber photometry recordings from an observer during interacting with a demonstrator or toy mouse. **d-g** Peri-event averaged calcium response traces (left), heatmaps of individual trial calcium responses evoked in PVN^OXT^-OT neuronal fibers (middle), and quantifications of calcium-dependent fluorescence changes (far right panels; AUC, area under the curve; peak ΔF/F; pre, -3-0 s; post, 0-3 s) during observer’s allogrooming (**d**) and investigation (**e**) toward demonstrator, as well as rearing (**f**) and walking (**g**). **h-k** Corresponding analyses for observer interaction with a toy mouse (**h-k** mirror **d-g**, respectively). Total trials: 34, 32, 43, 39, 39, 43, 38, and 33 for **d-k**, respectively (6-10 trials/mouse from five OXT-Cre observer mice). Dashed lines indicate behavior onset, and shaded areas denote SEM. The same cohort of demonstrators (C57BL/6 mice, n = 5; **d-g**) were used. Data are presented as mean ± SEM. \**p*□<□0.05, \*\**p*□<□0.01; ns, no significance. Statistical details see Table S1.

**Figure S9.**
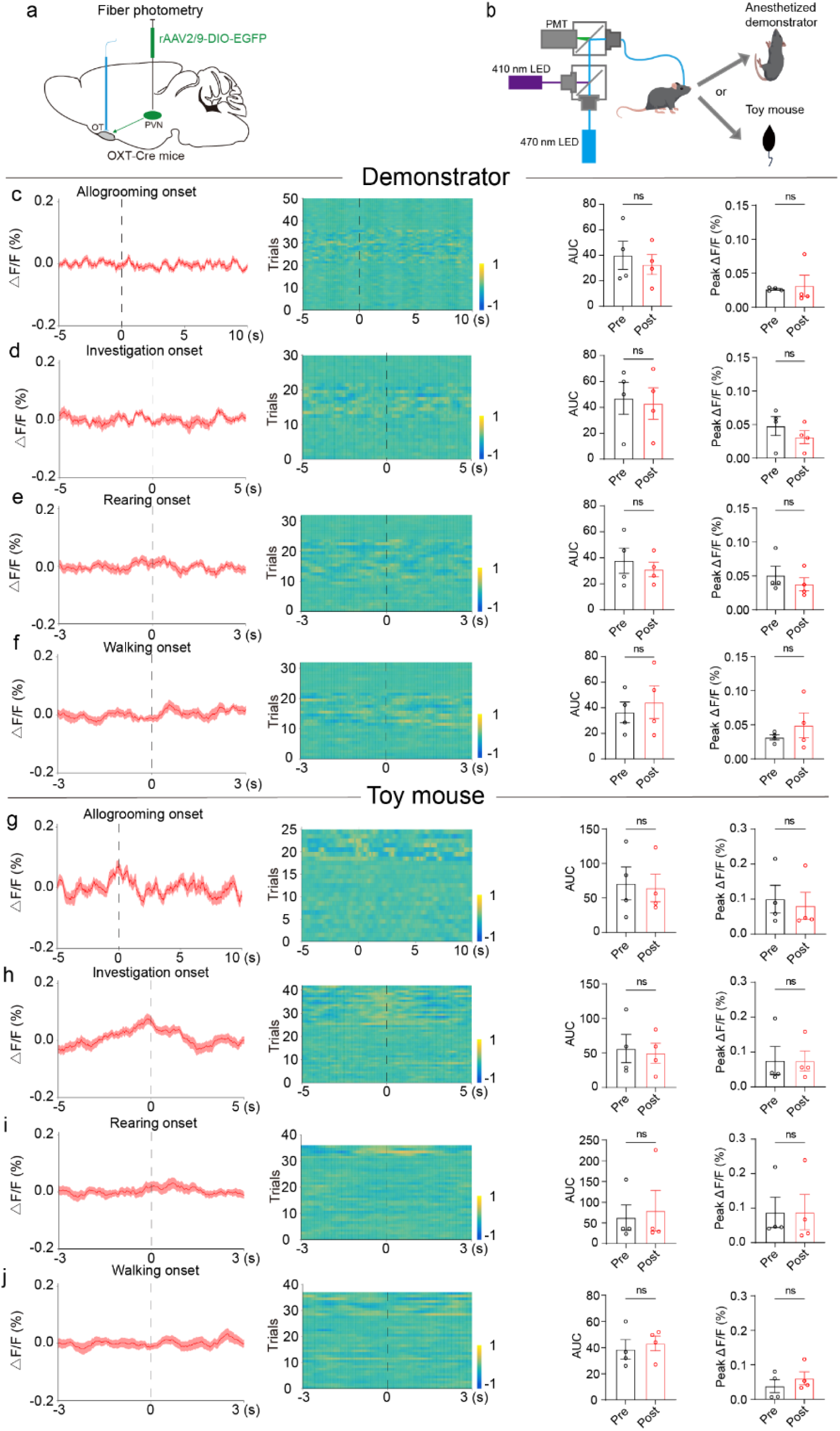
PVN^OXT^-OT neuronal fibers show no apparent time-locked responses to EGFP-expressing observer’s allogrooming or social investigation of anesthetized demonstrator. **Related to Figure 3, 4. a** Schematic illustrating viral injection strategy. **b** Schematic of fiber photometry recordings from an observer during interacting with demonstrator or toy mouse. **c-f** Peri-event averaged calcium response traces (left), heatmaps of individual trial calcium responses evoked in PVN^OXT^-OT neuronal fibers (middle), and quantifications of calcium-dependent fluorescence changes (far right panels; AUC, area under the curve; peak ΔF/F; pre, -3-0 s; post, 0-3 s) during observer’s allogrooming (**c**) and investigation (**d**) toward demonstrator, as well as rearing (**e**) and walking (f). **g-j** Corresponding analyses for observer interaction with a toy mouse (**g-j** mirror **c-f**, respectively). Total trials: 50, 30, 32, 32, 25, 42, 36, and 37 for c-j, respectively (6-15 trials/mouse from four OXT-Cre observer mice). Dashed lines indicate behavior onset, and shaded areas denote SEM. The same cohort of demonstrators (C57BL/6 mice, n = 4, **c-f**) were used. Data are presented as mean ± SEM. ns, no significance. Statistical details see Table S1.

**Figure S10.**
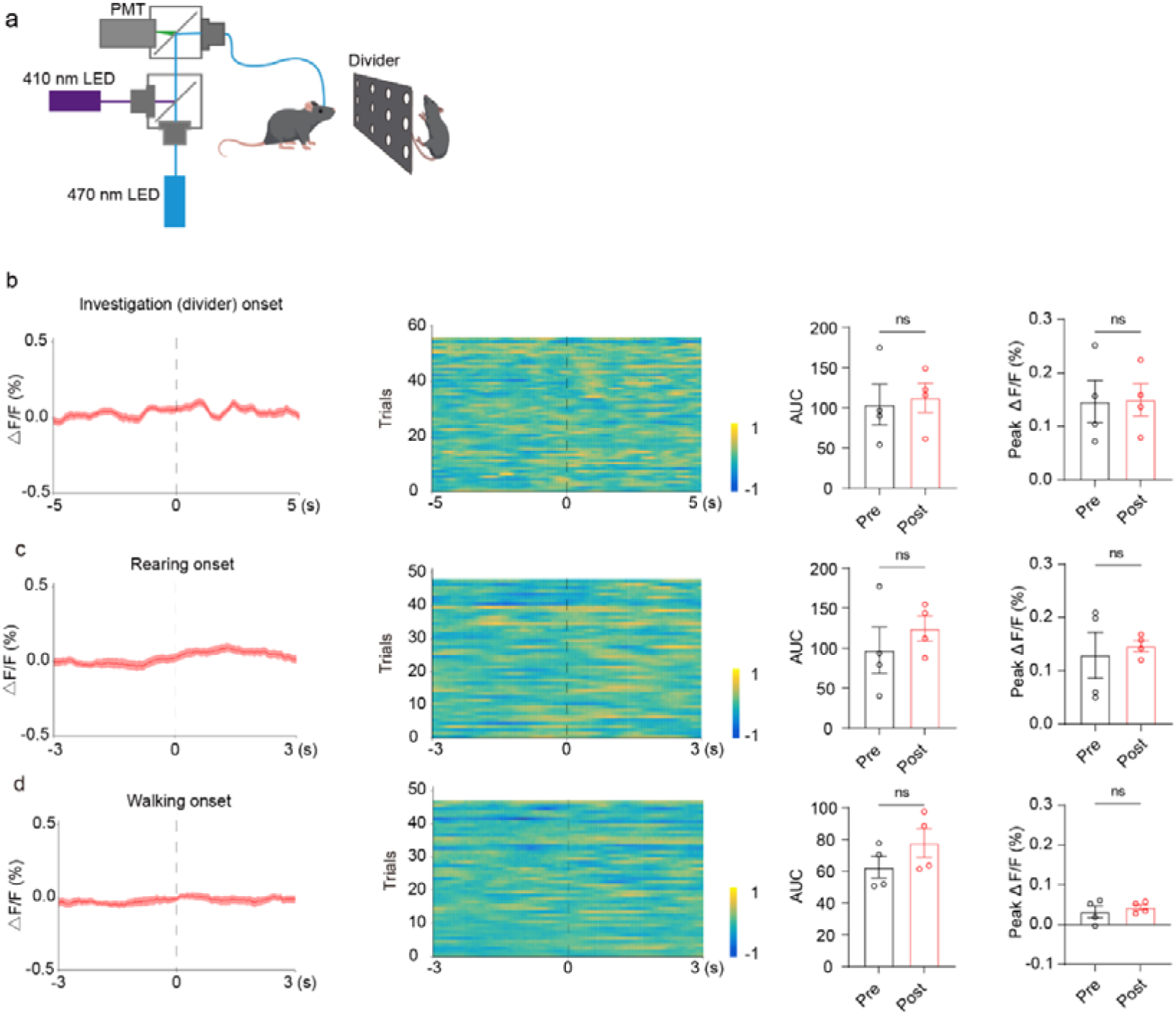
OXT release in the OT showed no detectable time-locked changes during observer’s non-contact social behaviors toward anesthetized demonstrator. **Related to Figure 5. a** Schematic of fiber photometry recordings from an observer during a non-contact social paradigm with a demonstrator. **b-d** Peri-event averaged OXT-dependent fluorescence traces (left), heatmaps of individual trial fluorescence changes evoked in the OT (middle), and quantifications of OXT-dependent fluorescence changes (far right panels; AUC, area under the curve; peak ΔF/F; pre, -3-0 s; post, 0-3 s) during observer’s investigation (**b**) toward divider, as well as rearing (**c**) and walking (**d**). Total trials: 55, 48, and 47 for **b-d**, respectively (10-15 trials/mouse from four C57BL/6 observer mice). Dashed lines indicate behavior onset, and shaded areas denote SEM. The same cohort of demonstrators (C57BL/6 mice, n = 4, **b-d**) were used. Data are presented as mean ± SEM. ns, no significance. Statistical details see Table S1.

**Figure S11.**
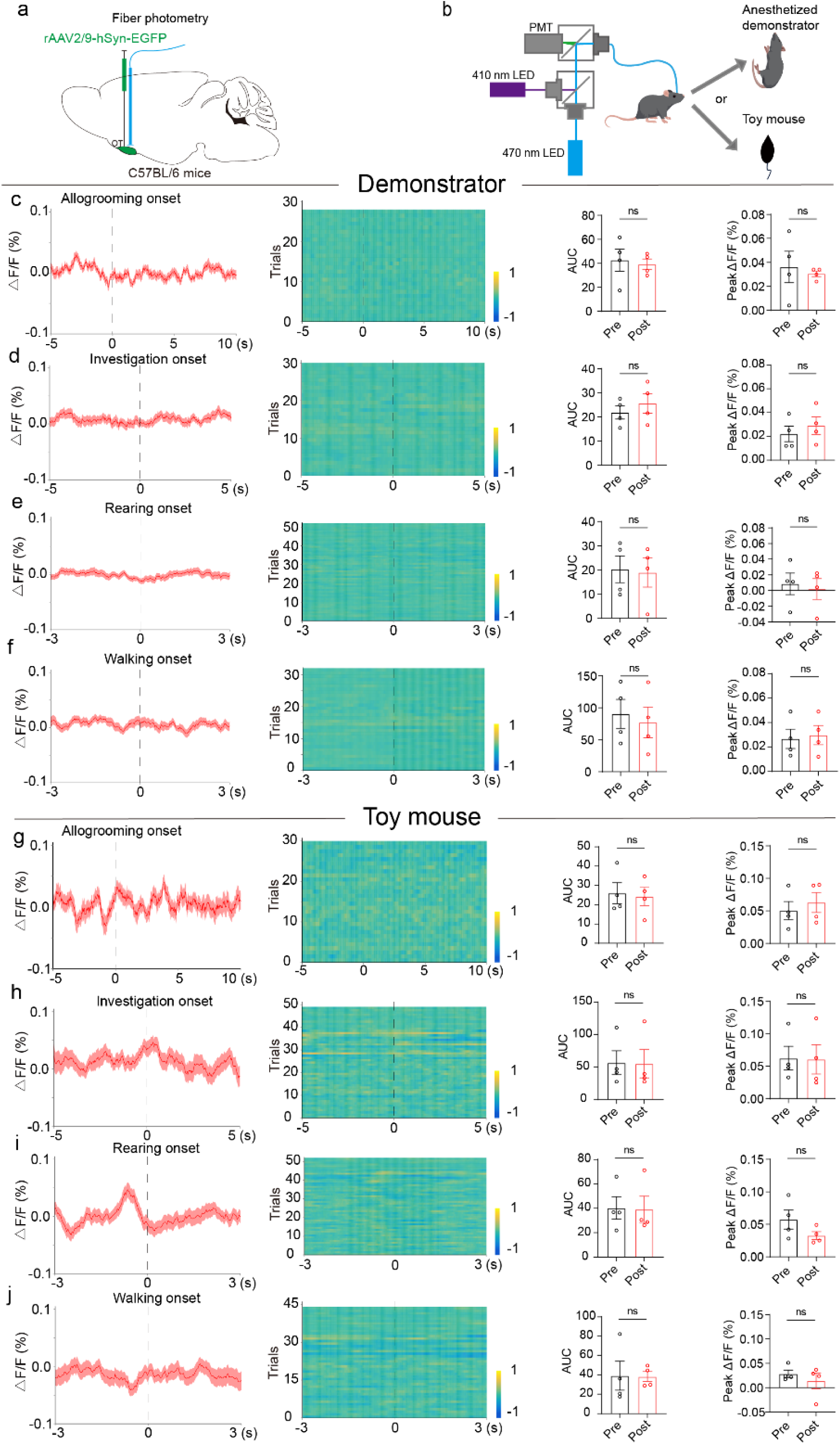
No apparent fluorescent changes were detected during EGFP-expressing observer’s allogrooming or social investigation of anesthetized demonstrator. **Related to Figure 5. a** Schematic illustrating viral injection strategy. **b** Schematic of fiber photometry recordings from an observer during interacting with demonstrator or toy mouse. **c-f** Peri-event averaged calcium response traces (left), heatmaps of individual trial fluorescence changes evoked in the OT (middle), and quantifications of calcium-dependent fluorescence changes (far right panels; AUC, area under the curve; peak ΔF/F; pre, -3-0 s; post, 0-3 s) during observer’s allogrooming (**c**) and investigation (**d**) toward demonstrator, as well as rearing (**e**) and walking (f). **g-j** Corresponding analyses for observer interaction with a toy mouse (**g-j** mirror **c-f**, respectively). Total trials: 28, 29, 52, 33, 30, 49, 52, and 44 for c-j, respectively (6-15 trials/mouse from four C57BL/6 observer mice). Dashed lines indicate behavior onset, and shaded areas denote SEM. The same cohort of demonstrators (C57BL/6 mice, n = 4, **c-f**) were used. Data are presented as mean ± SEM. ns, no significance. Statistical details see Table S1.

**Figure S12.**
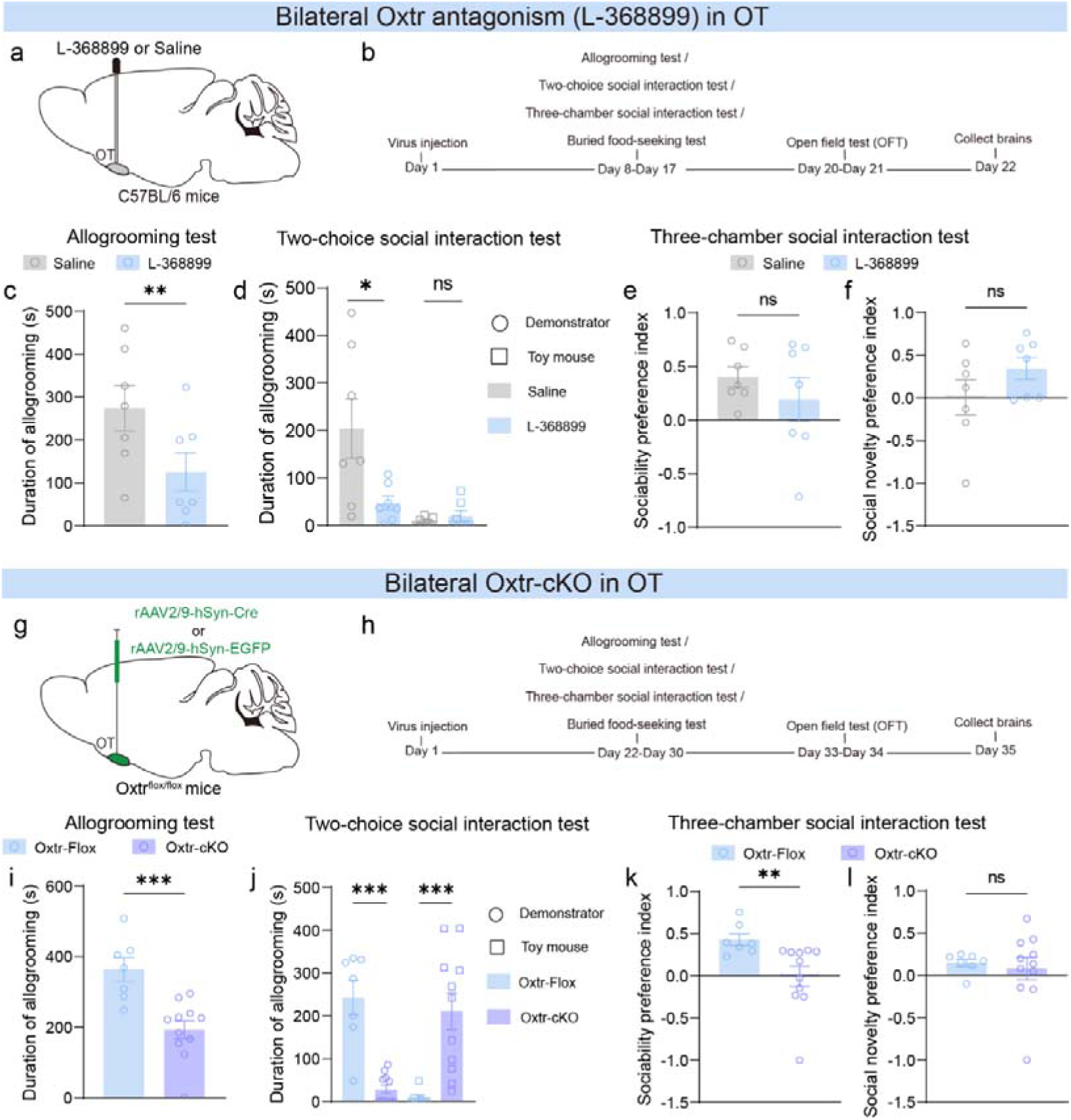
OXTR signaling in the OT is necessary for observer’s allogrooming toward anesthetized demonstrator. **Related to Figure 5. a** Schematic of viral injection strategy. **b** Behavioral assay timeline. **c-f** OXTR antagonism in the OT reduced observer allogrooming (**c**; allogrooming test) and decreased observer-biased allogrooming toward the demonstrator (**d**; two-choice test), but did not affect social behaviors (**e-f**; three-chamber test). **g** Schematic of viral injection strategy. **h** Behavioral assay timeline. **i-l** OXTR conditioned knockout (OXTR-cKO) in the OT reduced observer allogrooming (**i**; allogrooming test), decreased observer-biased allogrooming toward the demonstrator (**j**; two-choice test), and impaired social recognition but did not affect social novelty (**k-l**; three-chamber test). The antagonist-treated cohort (observers: n = 7; demonstrators: n = 7) was tested in **c-f**. The OXTR-cKO cohort (observers: n = 8 cKO, n = 7 Flox controls; demonstrators: n = 8) was tested in **i-l**. Data are presented as mean ± SEM. \**p*□<□0.05, \*\**p*□<□0.01, \*\*\**p*□<□0.001; ns, no significance. Statistical details see Table S1.

**Figure S13.**
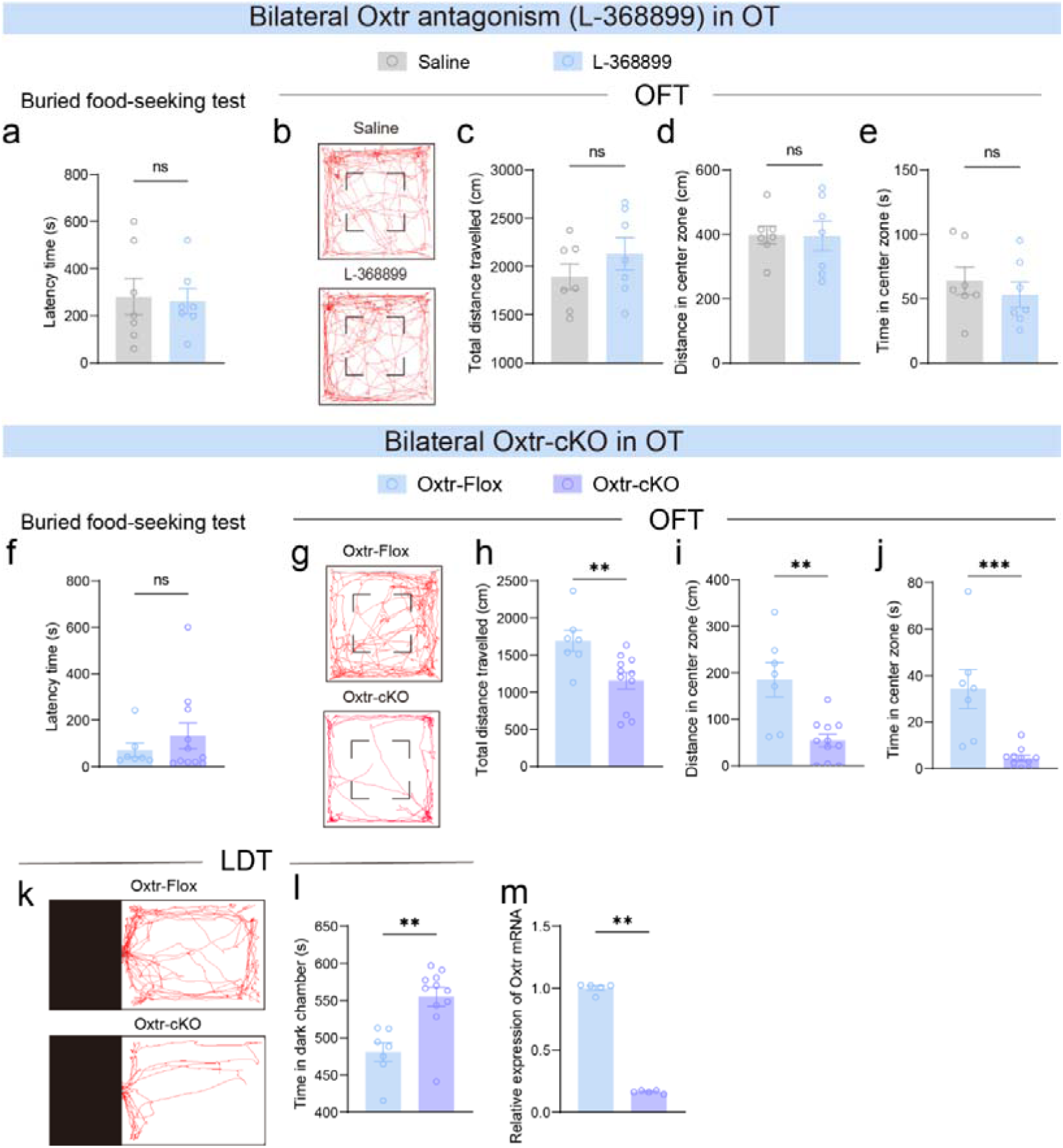
OXTR-cKO, but not OXTR antagonism, in the OT triggers anxiety-like behaviors in observer. **Related to Figure 5. a-g** Behavioral performance of OXTR-antagonized observers in the buried food-seeking test (**a**) and OFT (**b-e**). **a** Latency to find first food pellet. **B** Representative locomotor tracks. **c** Total distance travelled. **d, e** Distance travelled (**d**) and time spent (**e**) in center zone. **f-j** Parallel analyses for OXTR-cKO observers (**f-j** correspond to **a-e**, respectively). **k-l** Representative locomotion tracks (**k**) and time spent in dark chamber (**l**) of OXTR-cKO observers in LDT. **m** OXTR-cKO efficiency verification via qPCR. OXTR antagonism experiments (**a-e**): independent cohort (n = 7 observers). OXTR-cKO experiments (**f-l**): observers (n = 7 OXTR-Flox controls, n = 11 OXTR-cKO), with separate verification cohort (**m**; OXTR-Flox controls, n = 5; OXTR-Flox controls with AAV-Cre, n = 5). Data are presented as mean ± SEM. ** *p*□<□0.01, \*\*\**p*□<□0.001; ns, no significance. Statistical details see Table S1.

**Figure S14.**
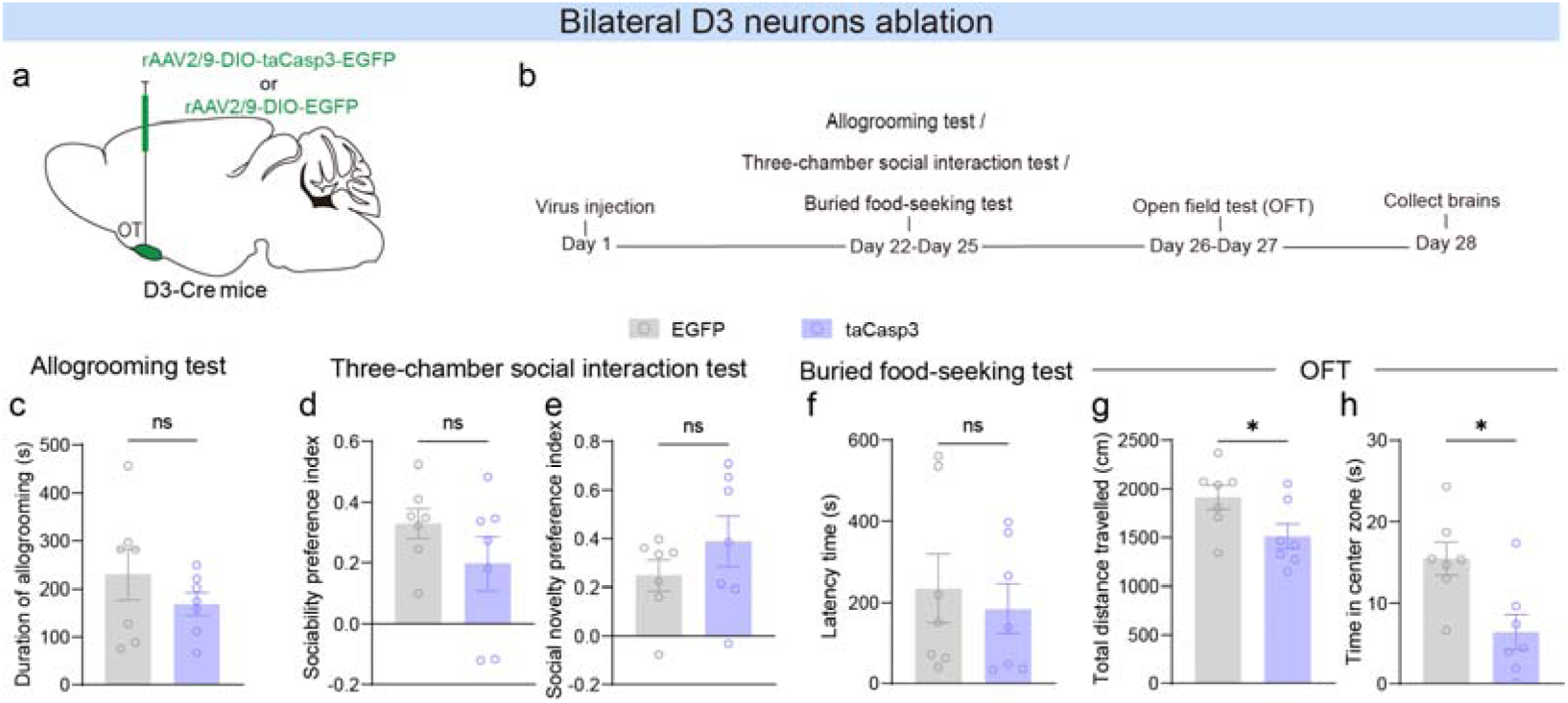
Ablation of D3 neurons in the OT does not affect observer’s allogrooming toward anesthetized demonstrator. **Related to Figure 6. a** Schematic of viral injection strategy. **b** Behavioral assay timeline. **c-h** Bilateral ablation D3 neurons has no significant effects on allogrooming (**c**; allogrooming test), sociability and social novelty preference (**d-e**; three-chamber test), as well as olfactory perception (**f**; buried food-seeking test), but decreasing locomotion (**g-h**; open field test, OFT). In **c-h**, observers: D3-Cre mice controls (n = 7), D3-Cre mice controls with AAV-DIO-taCasp3 (n = 7); Demonstrators C57BL/6 (n = 8, **c-e**). Data are presented as mean ± SEM. \**p*□<□0.05; ns, no significance. Statistical details see Table S1.

**Figure S15.**
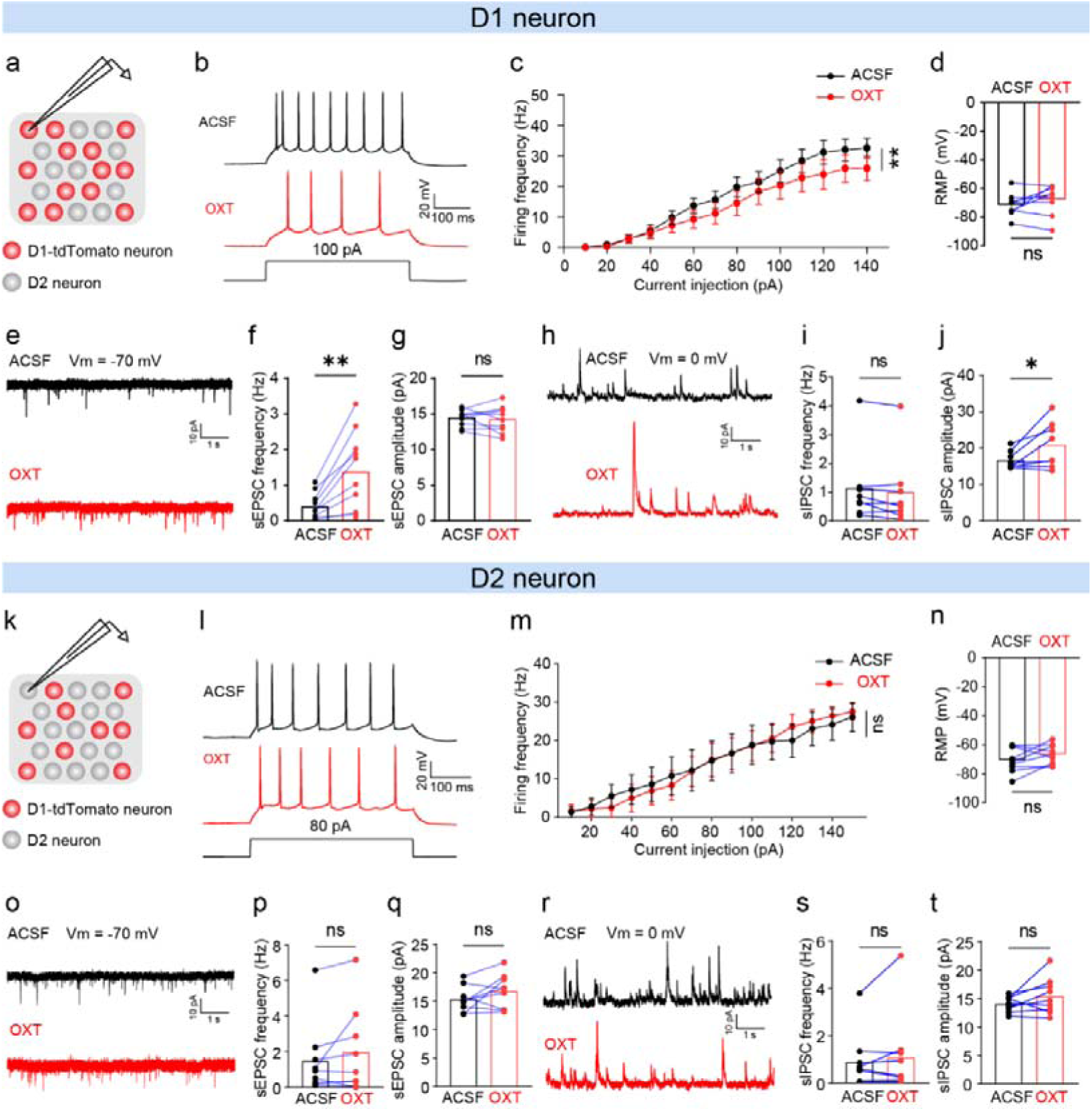
OXT induces hypoexcitability in D1 neurons but does not influence of D2 neural excitability in the OT. **Related to Figure 6. a** Schematic showing patch clamp recordings on D1-tdTomato neurons and D2 neurons. **b-d** Representative traces showing the firing of a D1 neuron pre- versus post-OXT treatment (**b**) and comparison of firing frequency (**c**) and resting membrane potential (RMP) between the two groups (**d**). **e-j** Representative sEPSCs (**e**) and sIPSCs (**h**) traces of D1 neurons pre- versus post-OXT treatment and comparison of corresponding frequency (**f, i**) and amplitude (**g, j**) between the two groups. **k-t** Same as **a-j**, respectively, but for D2 neurons. D1 neurons (n = 9 for **c-d**; n = 10 for **g-h**; n = 8 for **i-j**) and D2 neurons (n = 9) were recorded from three D1-tdTomato mice. Data are presented as mean ± SEM. \**p*□<□0.05, \*\**p*□<□0.01; ns, no significance. Statistical details see Table S1.

**Figure S16.**
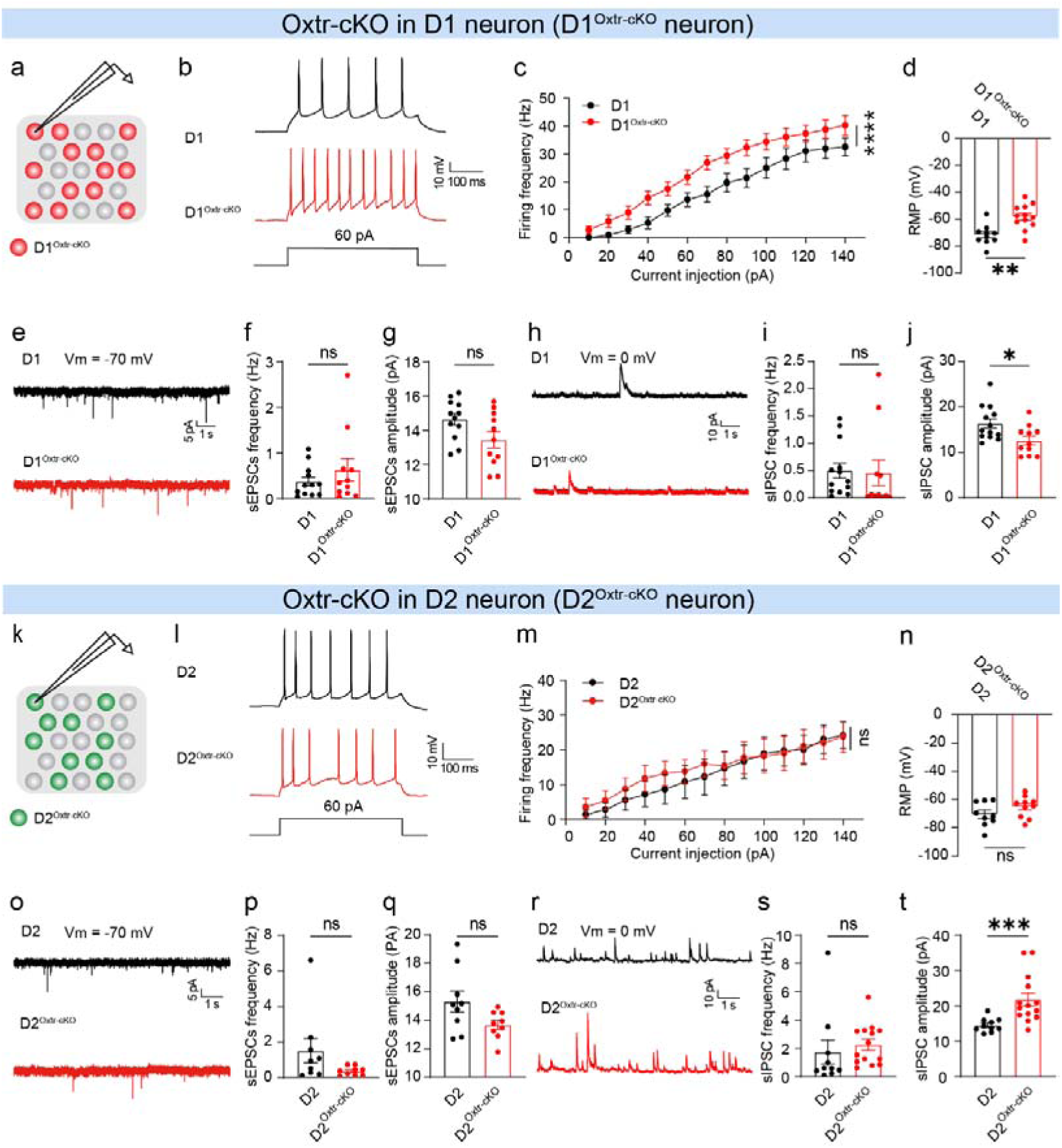
Oxtr-cKO induces hyperexcitability in D1 neurons but does not influence D2 neural excitability in the OT. **Related to Figure 6. a** Schematic showing patch clamp recordings on D1-EGFP neurons with Oxtr-cKO (D1^Oxtr-cKO^ neurons). **b-d** Representative traces showing firing of a D1 and a D1^Oxtr-cKO^ neuron (**b**) and comparison of firing frequency (**c**) and resting membrane potential (RMP) between the two groups (**d**). **e-j** Representative sEPSCs (**e**) and sIPSCs (**h**) traces of D1 neurons and D1^Oxtr-cKO^ neurons and comparison of corresponding frequency (**f, i**) and amplitude (**g, j**) between the two groups. **k-t** Same as **a-g**, respectively, but for D2^Oxtr-cKO^ neurons. D1 neurons (n = 10 for **c-d**; n = 12 for **f-g**; n = 13 for **i-j**); D1^Oxtr-cKO^ neurons (n = 12 for **c-d**; n = 11 for **f-g** and **i-j**); D2 neurons (n = 9 for **m-n** and **p-q**; n = 10 for **s-t**); D2^Oxtr-cKO^ neurons (n = 10 for **m-n**; n = 9 for **p-q**; n = 14 for **s-t**). n = 3 mice per group. The datasets of D1 and D2 neurons were from Figure S12. Data are presented as mean ± SEM. \**p*□<□0.05, \*\**p*□<□0.01; ns, no significance. Statistical details see Table S1.

**Figure S17.**
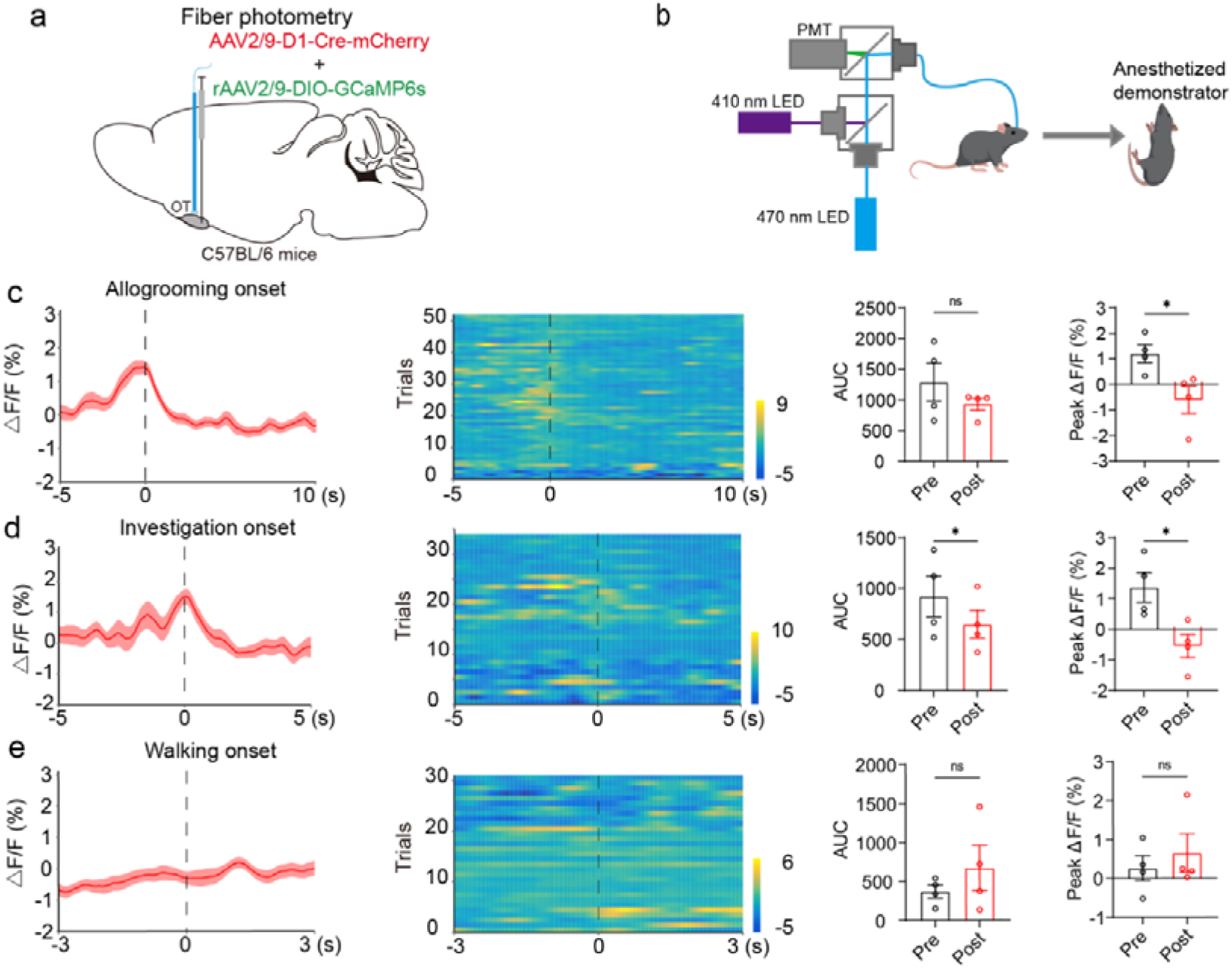
GCaMP6s-expressing D1 neurons in the OT respond to observer’s allogrooming or social investigation toward anesthetized demonstrator. **Related to Figure 6. a** Schematic illustrating viral injection strategy. **b** Schematic of fiber photometry recordings from an observer during interacting with a demonstrator. **c-e** Peri-event averaged calcium response traces (left), heatmaps of individual trial calcium responses evoked in D1 neurons (middle), and quantifications of calcium-dependent fluorescence changes (far right panels; AUC, area under the curve; peak ΔF/F; pre, -5-0 s; post, 0-5 s) during observer’s allogrooming (**c**) and investigation (**d**) toward demonstrator, as well as walking (**e**). Total trials: 53, 34, and 31 for c-e, respectively (7-14 trials/mouse from four C57BL/6 observer mice). Dashed lines indicate behavior onset, and shaded areas denote SEM. The same cohort of demonstrators (C57BL/6 mice, n = 4; **c-e**) were used. Data are presented as mean ± SEM. \**p*□<□0.05; ns, no significance. Statistical details see Table S1.

**Figure S18.**
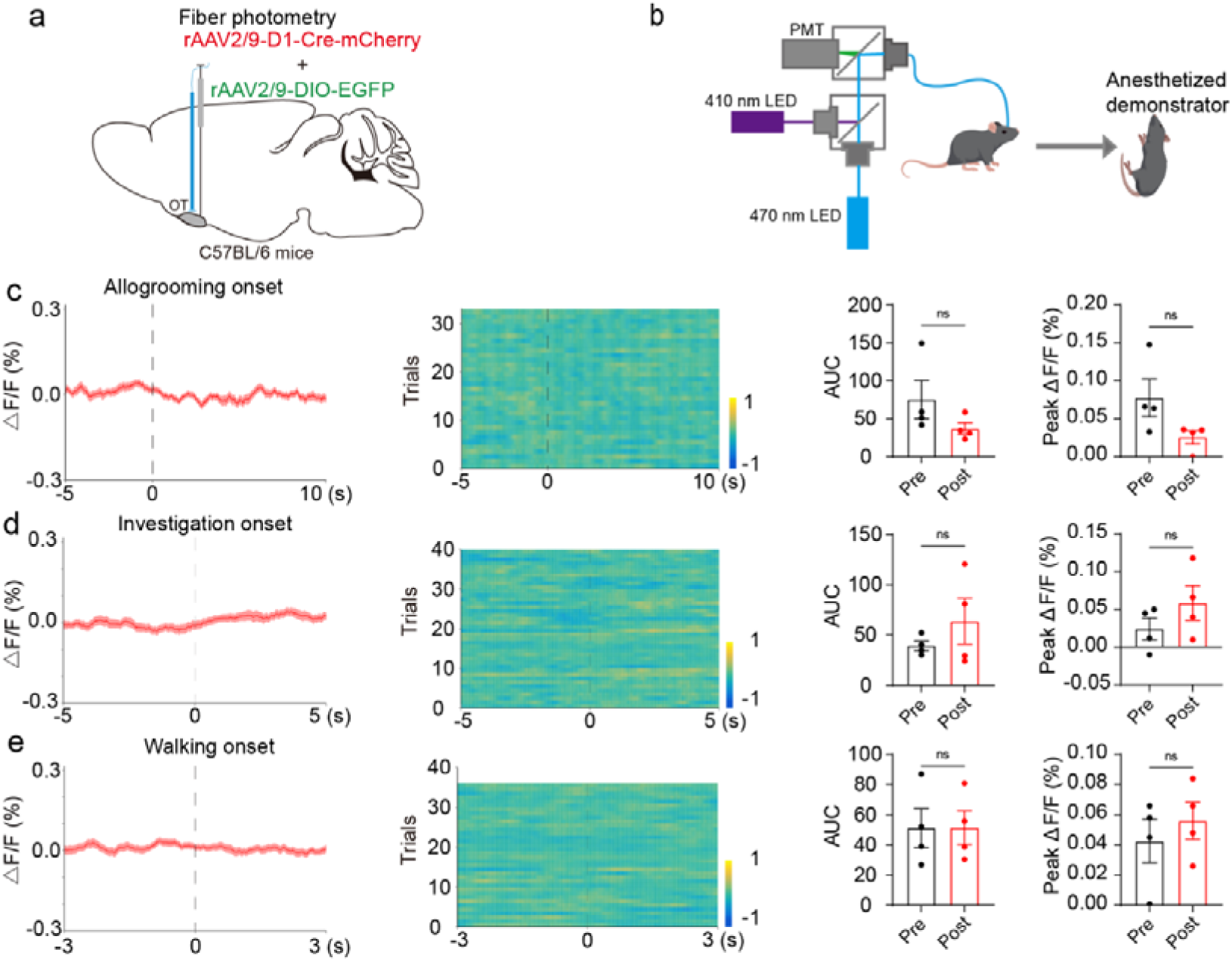
EGFP-expressing D1 neurons in the OT do not respond to observer’s allogrooming or social investigation toward anesthetized demonstrator. **Related to Figure 6. a** Schematic illustrating viral injection strategy. **b** Schematic of fiber photometry recordings from an observer during interacting with a demonstrator. **c-e** Peri-event averaged calcium response traces (left), heatmaps of individual trial calcium responses evoked in D1 neurons (middle), and quantifications of calcium-dependent fluorescence changes (far right panels; AUC, area under the curve; peak ΔF/F; pre, -5-0 s; post, 0-5 s) during observer’s allogrooming (**c**) and investigation (**d**) toward demonstrator, as well as walking (**e**). Total trials: 34, 40, and 36 for **c-e**, respectively (8-12 trials/mouse from four C57BL/6 observer mice). Dashed lines indicate behavior onset, and shaded areas denote SEM. The same cohort of demonstrators (C57BL/6 mice, n = 4; **c-e**) were used. Data are presented as mean ± SEM. ns, no significance. Statistical details see Table S1.

**Figure S19.**
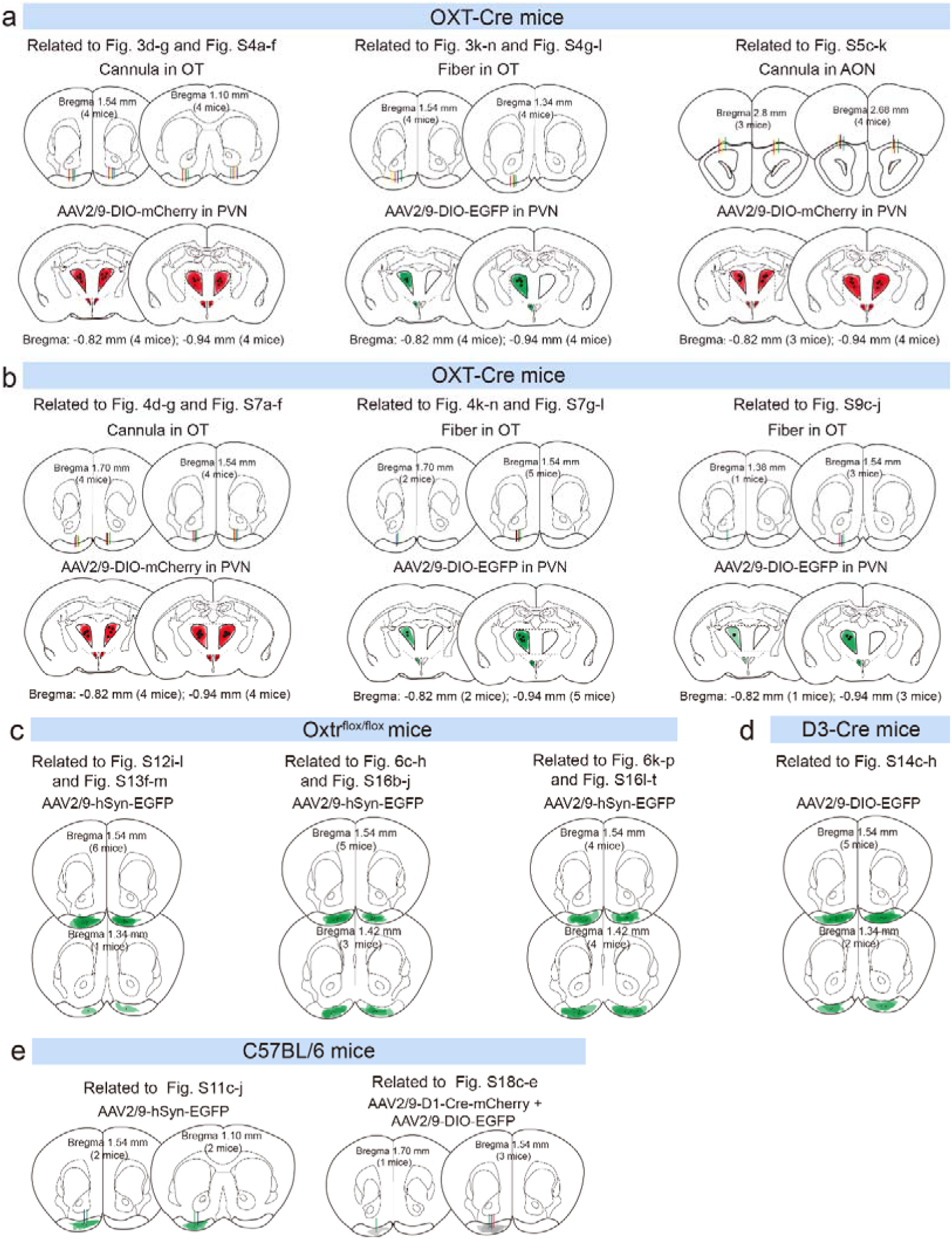
Viral injection/expression sites and optical fiber/cannula placements for experiments in Figures 3, 4, 6 and Figures S4, S5, S7, S9, S11-14, S16, S18 regarding control groups. Coronal brain sections at the bregma levels show viral injection sites (black dots), expression areas (red/green shadowed areas), and/or optical fiber/cannula tracts (vertical bars) for the included mice. Dotted rectangle areas are enlarged views of the PVN region. **a** OXT-Cre mice: loss-of-function of PVNOXT-OT pathway (mCherry, Fig. 3d-g, Fig. S4a-f an Fig. S5c-k; EGFP, Fig. 3k-n and Fig. S4g-l). **b** OXT-Cre mice: gain-of-function of PVNOXT pathway (mCherry, Fig. 4d-g and Fig. S7a-f; EGFP, Fig. 4k-n and Fig. S7g-l); Fiber photometry experiments (EGFP; Fig. S9c-j). **c** Oxtrflox/flox mice: EGFP expression in the OT (Fig. S12i-l and Fig. S13f-m); EGFP expression in OT D1 or D2 SPNs (D1, Fig. 6c-h and Fig. S16b-j; D2, Fig. 6k-p and Fig. S16l-t). **d** D3-Cre mice (EGFP; Fig. S14c-h). **e** C57BL/6 mice: Fiber photometry experiments (EGFP; Fig. S11c-j and Fig. S18c-e). Each dot or line represents one animal. Brain atlas images were adapted from the Allen Mouse Brain Common Coordinate Framework version 3 (https://scalablebrainatlas.incf.org/mouse/ABA_v3).

**Figure S20.**
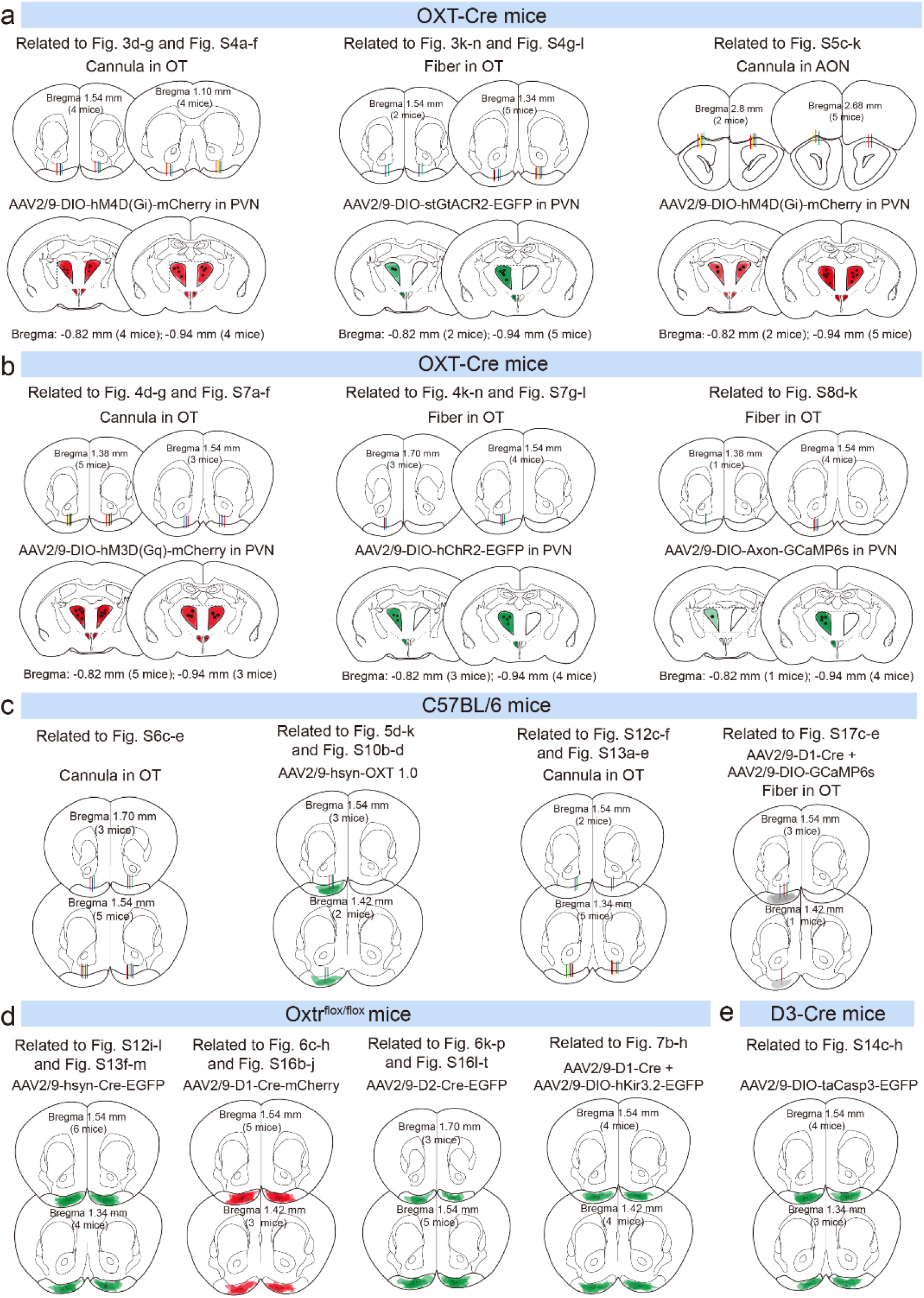
Viral injection/expression sites and optical fiber/cannula placements for experiments in Figures 3-7 and Figures S4-17 regarding experimental groups. Coronal brain sections at the bregma levels show viral injection sites (black dots), expression areas (red/green shadowed areas), and/or optical fiber/cannula tracts (vertical bars) for the included mice. Dotted rectangle areas are enlarged views of the PVN region. **a** OXT-Cre mice: loss-of-function of PVN^OXT^-OT pathway (Fig. 3d-g, k-n, Fig. S4 and Fig. S5c-k). **b** OXT-Cre mice: gain-of-function of PVN^OXT^-OT pathway (Fig. 4d-g, k-n and Fig. S7); Fiber photometry experiments (GCaMP6s; Fig. S8d-k). **c** C57BL/6 mice: bilateral OXT infusion into OT (Fig. S6c-e); Fiber photometry experiments (OXT sensor; Fig. 5d-k and Fig.S10b-d); Loss-of-function of OXTR in the OT (Pharmacology; Fig. S12c-f and Fig. S13a-e). Fiber photometry experiments (GCaMP6s; Fig. S17c-e). **d** Oxtr^flox/flox^ mice: Loss-of-function of OXTR in the OT (Genetic ablation; Fig. S12i-l and Fig. S13f-m); Loss-of-function of OXTR in D1 or D2 SPNs (D1, Fig. 6c-h and Fig. S16b-j; D2, Fig. 6k-p and Fig. S16l-t). GIRK channel overexpression (Fig. 7b-h). **e** D3-Cre mice: D3 neurons ablation (Fig. S14c-h). Each dot or line represents one animal. Brain atlas images were adapted from the Allen Mouse Brain Common Coordinate Framework version 3 (https://scalablebrainatlas.incf.org/mouse/ABA_v3).

**Table S1.**
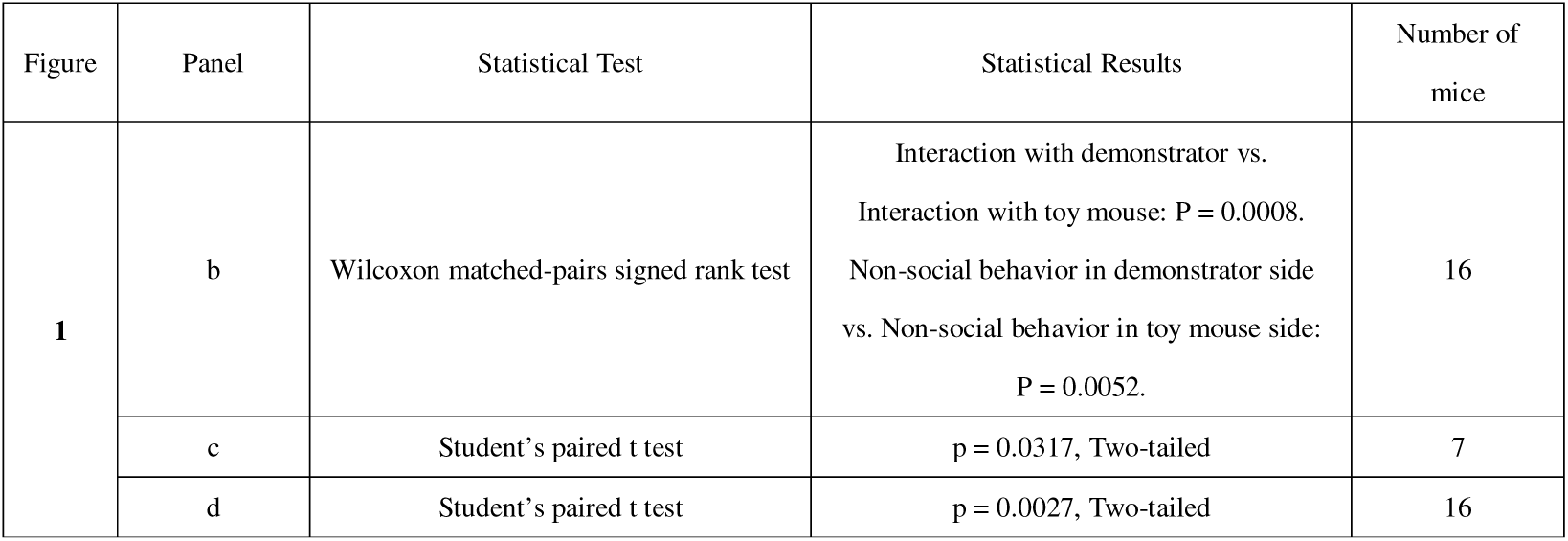
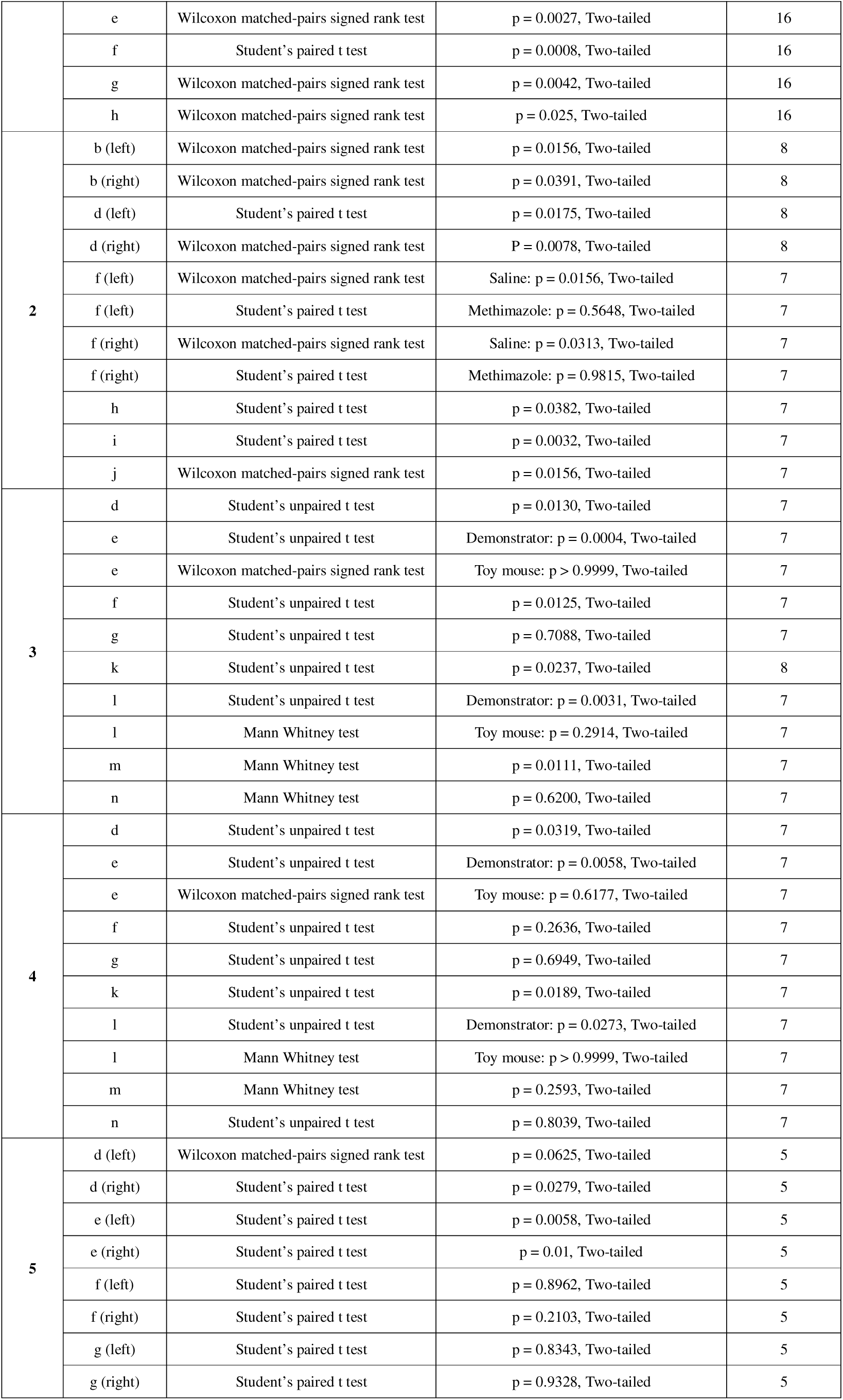
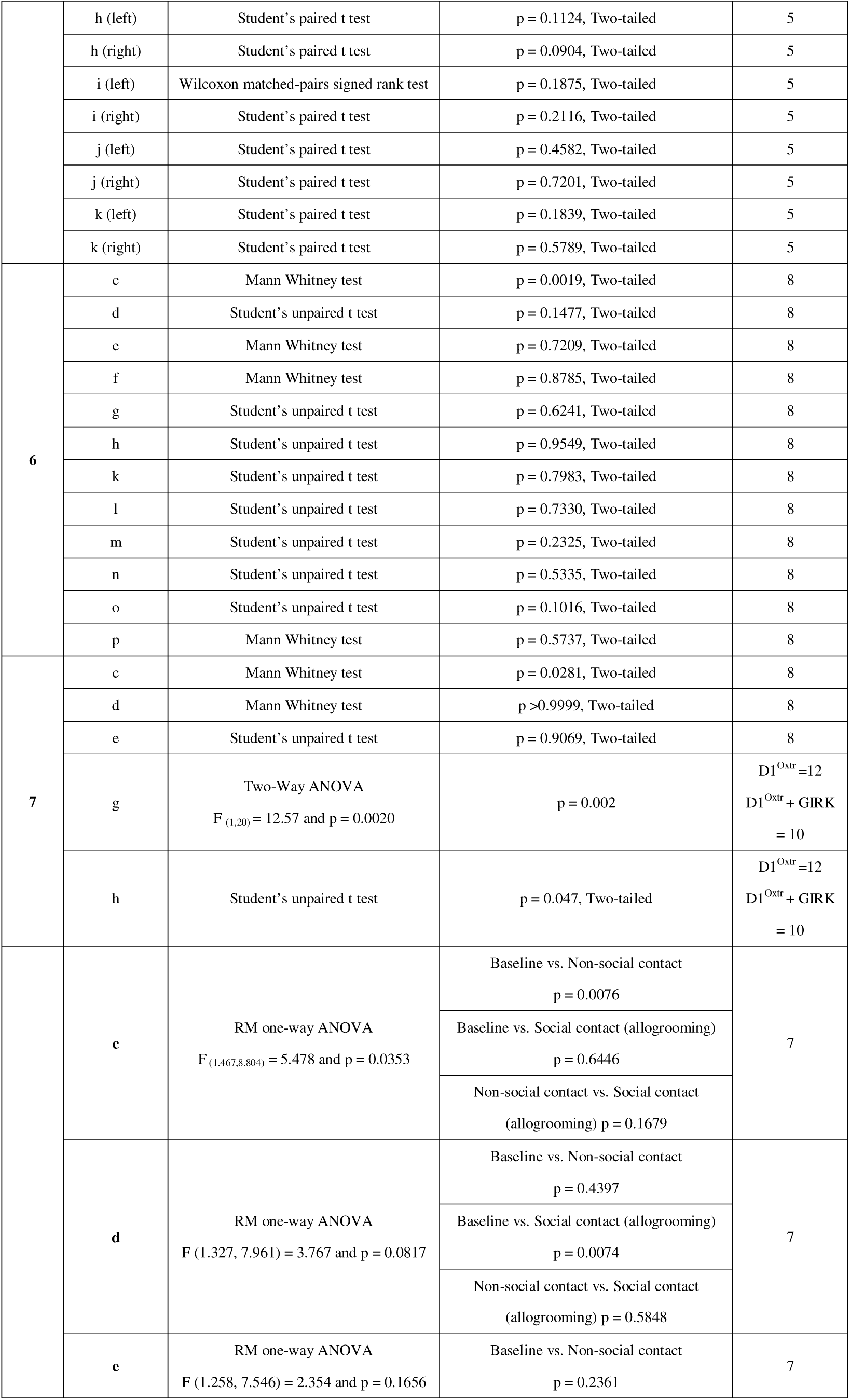
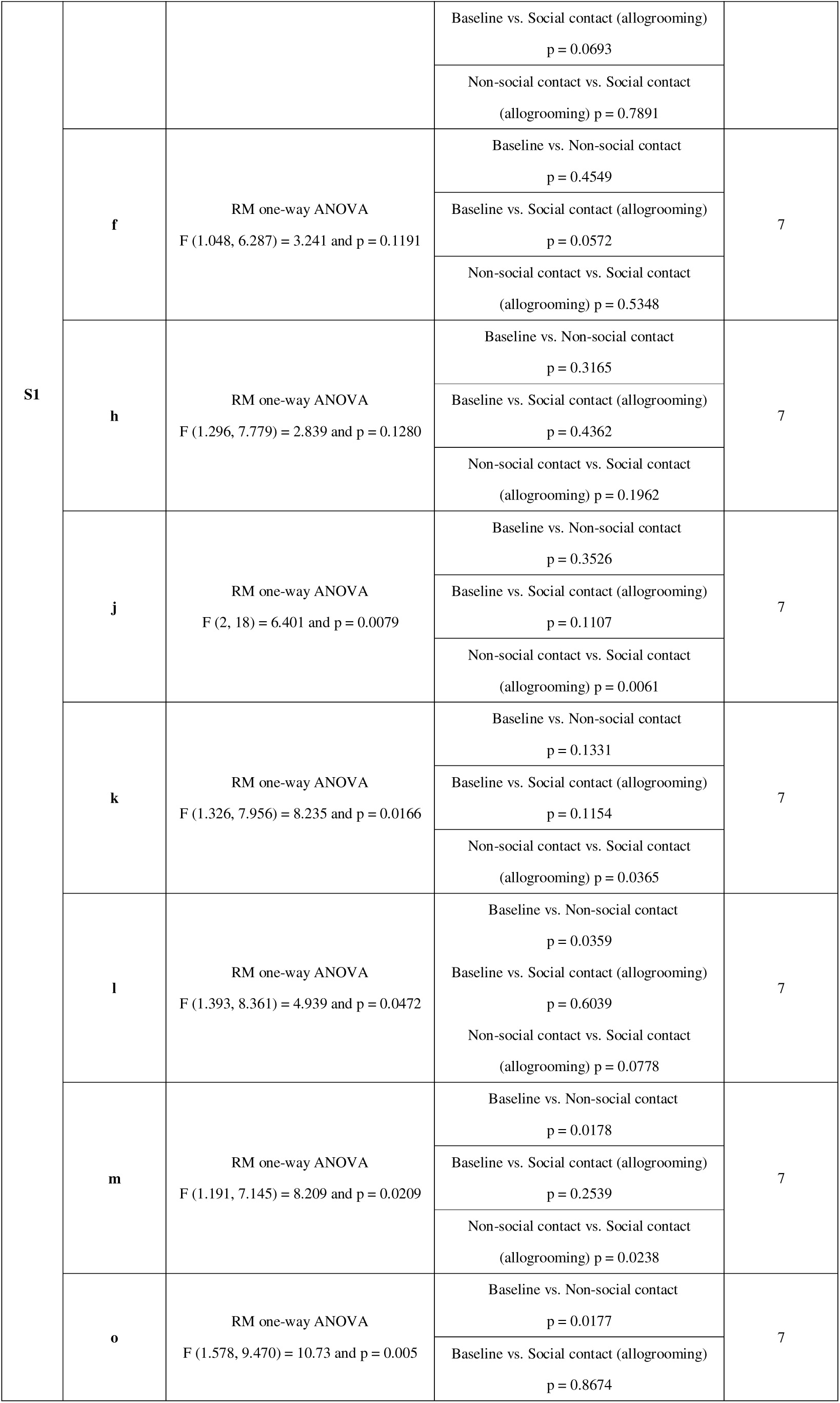
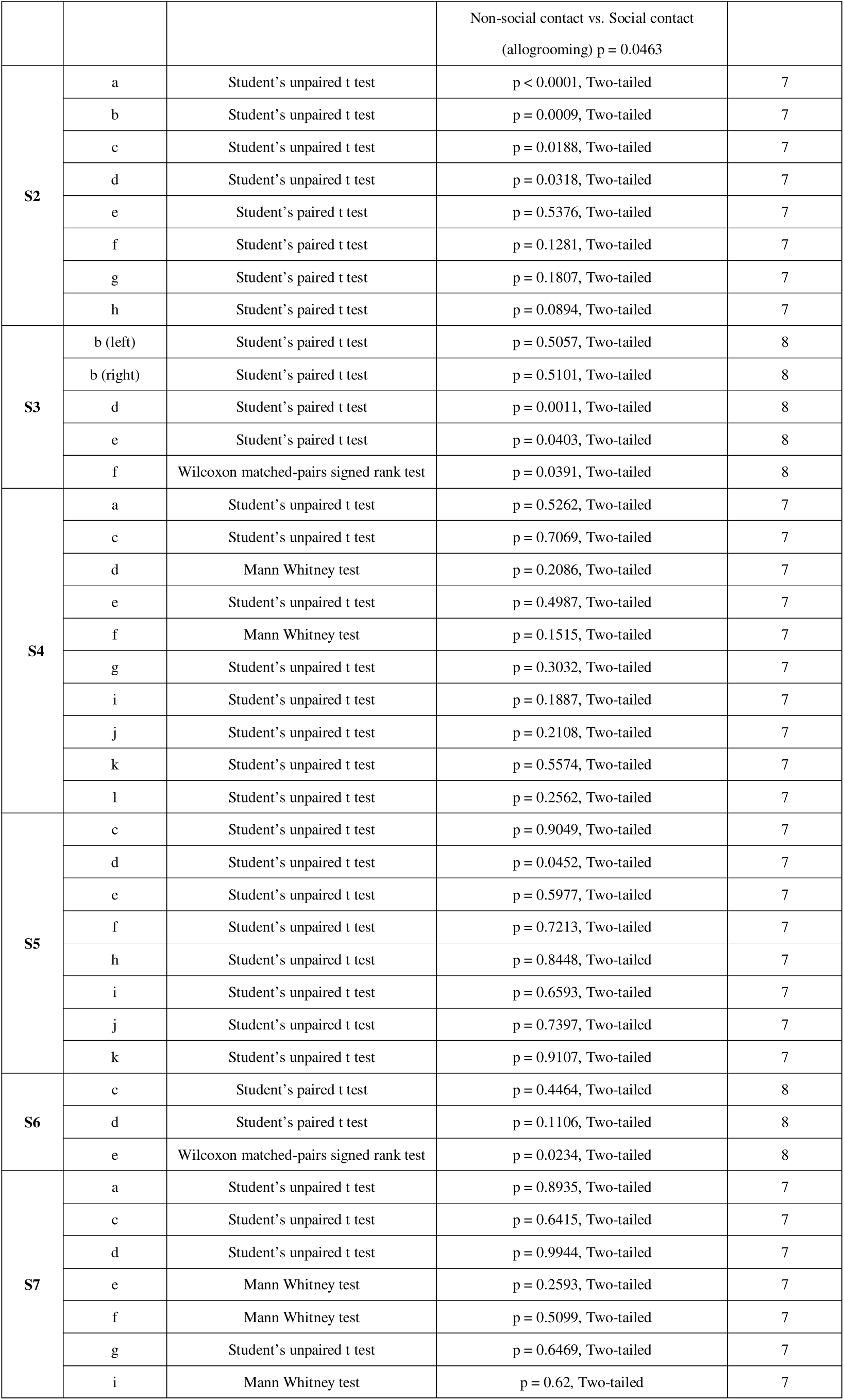
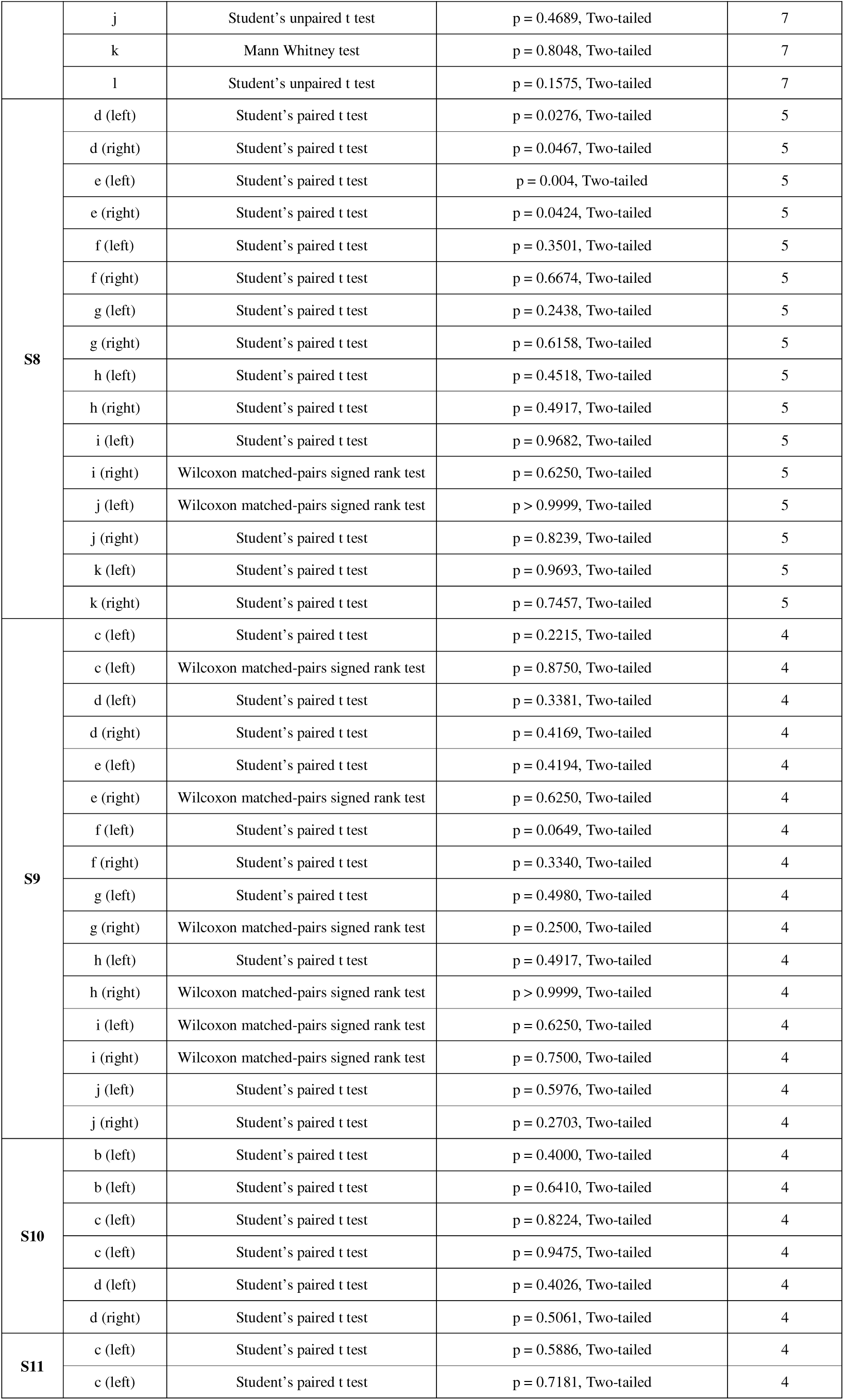
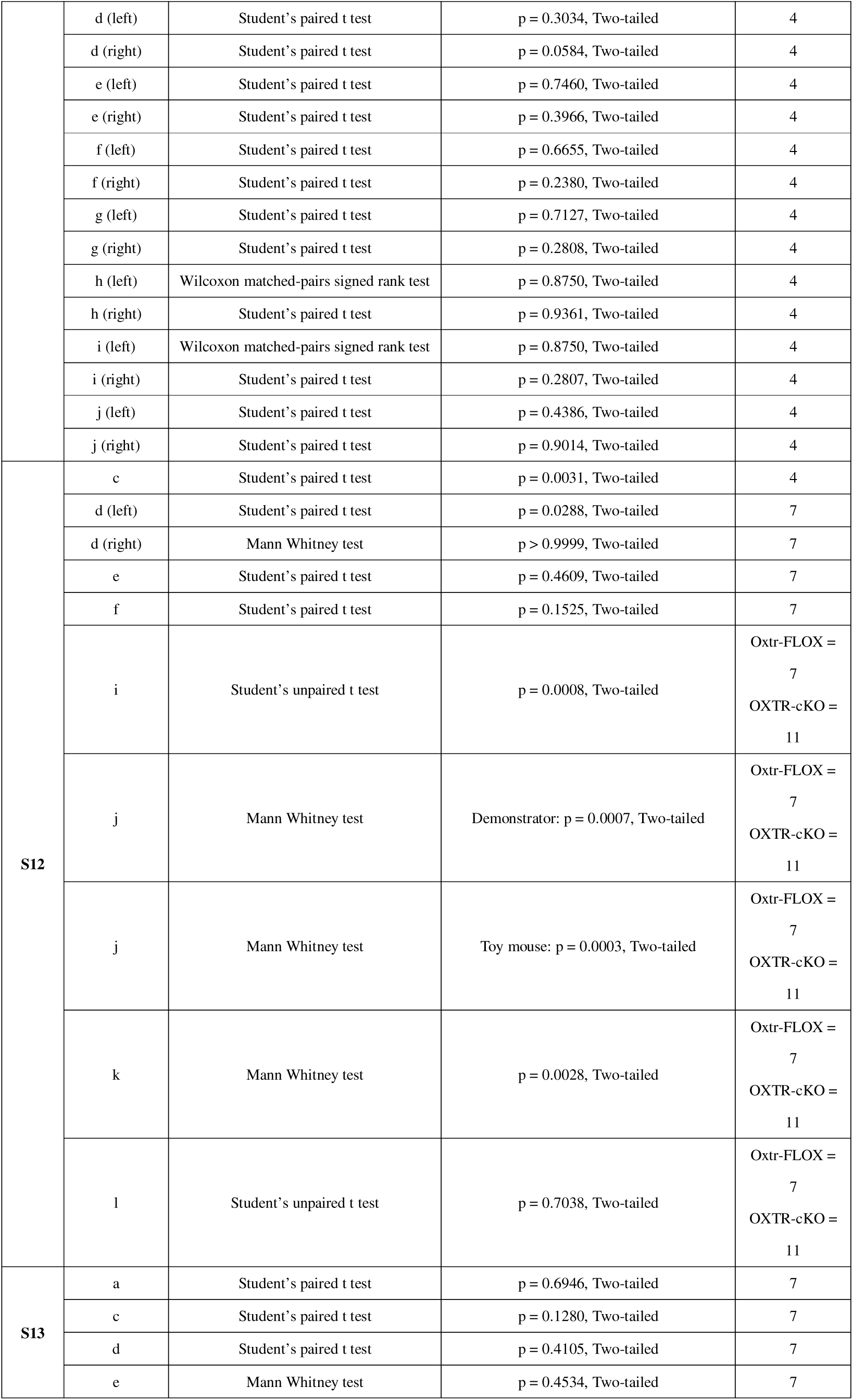
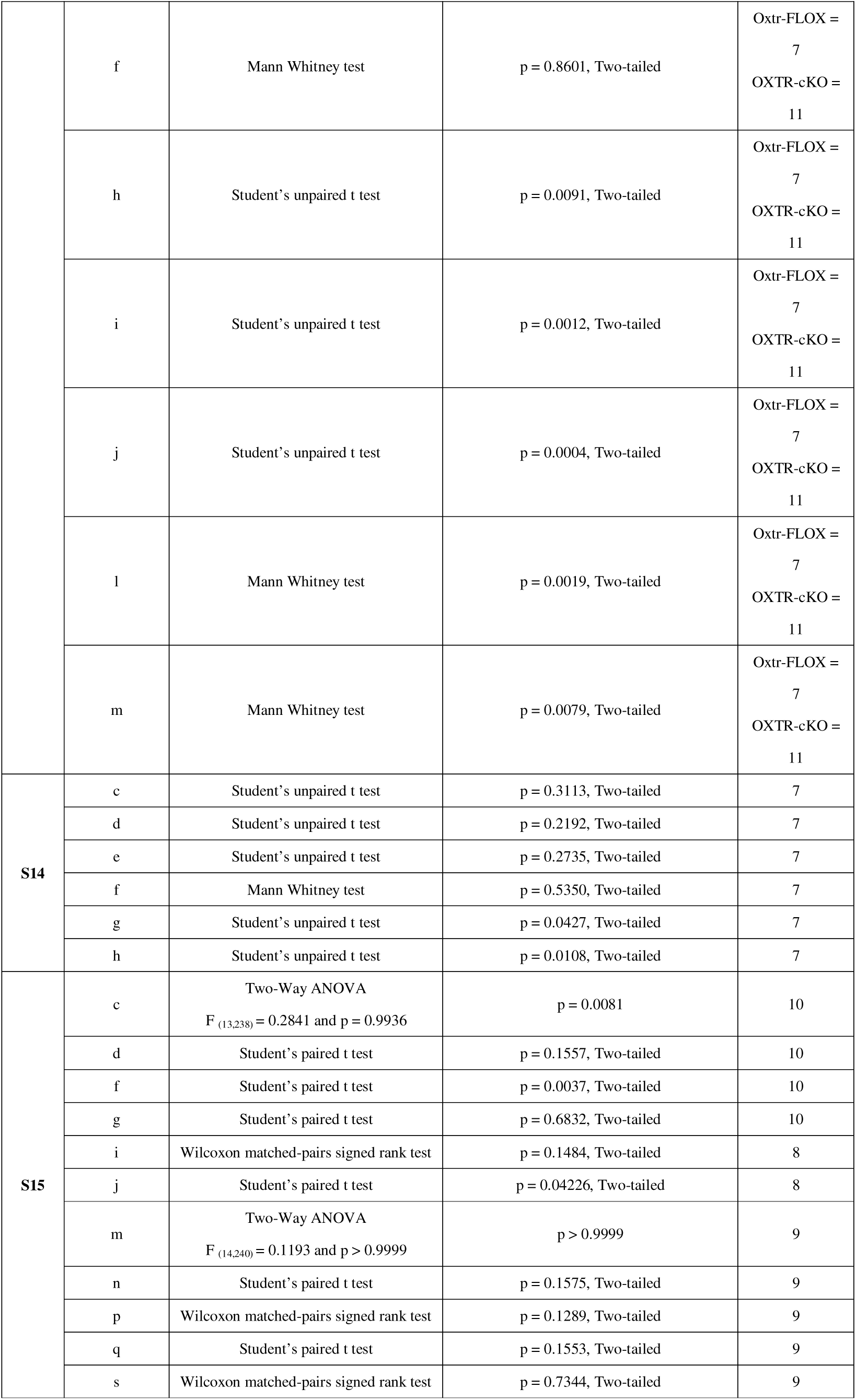
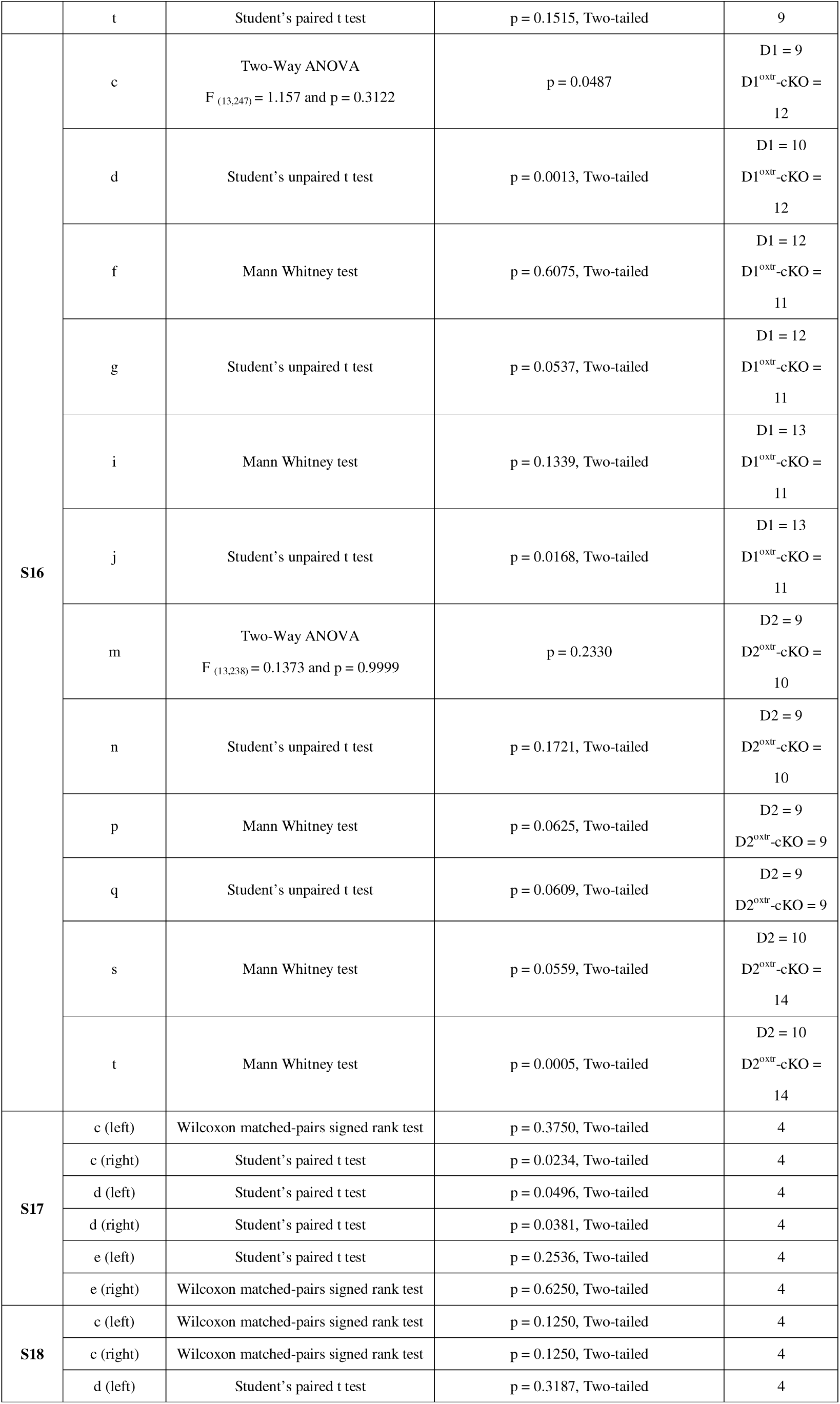
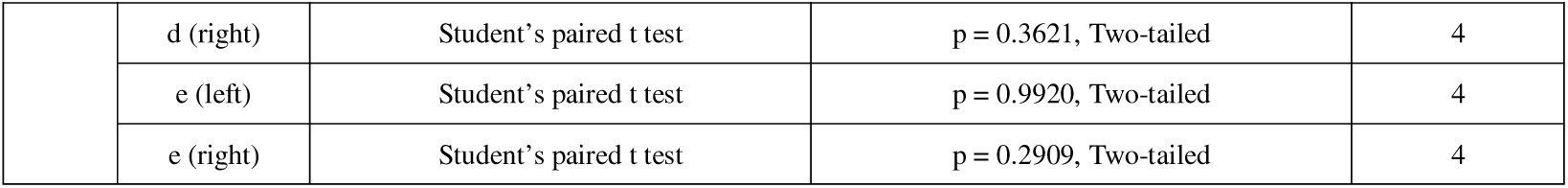
Statistical details.

## References

1. Scholl, W. (2013). The socio-emotional basis of human interaction and communication: How we construct our social world. Social Science Information 52, 3–33.

2. Bardis, P.D. (1979). Social interaction and social processes. Social Science 54, 147–167.

3. Quarantelli, E.L. (1960). Images of withdrawal behavior in disasters: Some basic misconceptions. Social Problems 8, 68–79.

4. Zhang, F.R., Liu, J., Wen, J., Zhang, Z.Y., Li, Y., Song, E., Hu, L., and Chen, Z.F. (2025). Distinct oxytocin signaling pathways synergistically mediate rescue-like behavior in mice. Proc Natl Acad Sci U S A 122, e2423374122. 10.1073/pnas.2423374122.

5. Decety, J., Bartal, I.B.-A., Uzefovsky, F., and Knafo-Noam, A. (2016). Empathy as a driver of prosocial behaviour: highly conserved neurobehavioural mechanisms across species. Philosophical Transactions of the Royal Society B: Biological Sciences 371, 20150077.

6. De Waal, F.B., and Preston, S.D. (2017). Mammalian empathy: behavioural manifestations and neural basis. Nature Reviews Neuroscience 18, 498–509.

7. De Waal, F.B. (2008). Putting the altruism back into altruism: the evolution of empathy. Annu. Rev. Psychol. 59, 279–300.

8. Sun, W., Zhang, G.W., Huang, J.J., Tao, C., Seo, M.B., Tao, H.W., and Zhang, L.I. (2025). Reviving-like prosocial behavior in response to unconscious or dead conspecifics in rodents. Science 387, eadq2677. 10.1126/science.adq2677.

9. Cao, P., Liu, Y., Ni, Z., Zhang, M., Wei, H.-R., Liu, A., Guo, J.-R., Yang, Y., Xu, Z., and Guo, Y. (2025). Rescue-like behavior in a bystander mouse toward anesthetized conspecifics promotes arousal via a tongue-brain connection. Science Advances 11, eadq3874.

10. Sun, F., Wu, Y.E., and Hong, W. (2025). A neural basis for prosocial behavior toward unresponsive individuals. Science 387, eadq2679. 10.1126/science.adq2679.

11. Schweinfurth, M.K., and Taborsky, M. (2018). Reciprocal trading of different commodities in Norway rats. Current Biology 28, 594–599. e593.

12. Hong, M., Zou, L., and Hu, R.K. (2025). Unraveling rescue-like behaviors: Neural circuits driving prosocial aid in mice. Neuron 113, 2034–2036. 10.1016/j.neuron.2025.06.004.

13. Dunbar, R.I. (2010). The social role of touch in humans and primates: behavioural function and neurobiological mechanisms. Neuroscience & Biobehavioral Reviews 34, 260–268.

14. Russell, Y.I., and Phelps, S. (2013). How do you measure pleasure? A discussion about intrinsic costs and benefits in primate allogrooming. Biol Philos 28, 1005–1020. 10.1007/s10539-013-9372-4.

15. Dunbar, R.I. (1991). Functional significance of social grooming in primates. Folia primatologica 57, 121–131.

16. Pfoh, R., Tiddi, B., Di Bitetti, M.S., and Agostini, I. (2021). Grooming site preferences in black capuchin monkeys: Hygienic vs. social functions revisited. American Journal of Primatology 83, e23336.

17. Schino, G., and Aureli, F. (2008). Grooming reciprocation among female primates: a meta-analysis. Biology Letters 4, 9–11.

18. Silk, J.B. (2007). The adaptive value of sociality in mammalian groups. Philosophical Transactions of the Royal Society B: Biological Sciences 362, 539–559.

19. Grebe, N.M., Sheikh, A., Ohannessian, L., and Drea, C.M. (2023). Effects of oxytocin receptor blockade on dyadic social behavior in monogamous and non-monogamous Eulemur. Psychoneuroendocrinology 150, 106044. 10.1016/j.psyneuen.2023.106044.

20. Laister, S., Stockinger, B., Regner, A.-M., Zenger, K., Knierim, U., and Winckler, C. (2011). Social licking in dairy cattle—Effects on heart rate in performers and receivers. Applied Animal Behaviour Science 130, 81–90.

21. Schino, G., Scucchi, S., Maestripieri, D., and Turillazzi, P.G. (1988). Allogrooming as a tension-reduction mechanism: A behavioral approach. Am J Primatol 16, 43–50. 10.1002/ajp.1350160106.

22. Burkett, J.P., Andari, E., Johnson, Z.V., Curry, D.C., de Waal, F.B., and Young, L.J. (2016). Oxytocin-dependent consolation behavior in rodents. Science 351, 375–378.

23. Tang, Y., Benusiglio, D., Lefevre, A., Hilfiger, L., Althammer, F., Bludau, A., Hagiwara, D., Baudon, A., Darbon, P., Schimmer, J., et al. (2020). Social touch promotes interfemale communication via activation of parvocellular oxytocin neurons. Nat Neurosci 23, 1125–1137. 10.1038/s41593-020-0674-y.

24. Knobloch, H.S., Charlet, A., Hoffmann, L.C., Eliava, M., Khrulev, S., Cetin, A.H., Osten, P., Schwarz, M.K., Seeburg, P.H., Stoop, R., and Grinevich, V. (2012). Evoked axonal oxytocin release in the central amygdala attenuates fear response. Neuron 73, 553–566. 10.1016/j.neuron.2011.11.030.

25. Borie, A.M., Young, L.J., and Liu, R.C. (2022). Sex-specific and social experience-dependent oxytocin-endocannabinoid interactions in the nucleus accumbens: implications for social behaviour. Philos Trans R Soc Lond B Biol Sci 377, 20210057. 10.1098/rstb.2021.0057.

26. Froemke, R.C., and Young, L.J. (2021). Oxytocin, Neural Plasticity, and Social Behavior. Annu Rev Neurosci 44, 359–381. 10.1146/annurev-neuro-102320-102847.

27. Yoshihara, C., Numan, M., and Kuroda, K.O. (2018). Oxytocin and Parental Behaviors. Curr Top Behav Neurosci 35, 119–153. 10.1007/7854_2017_11.

28. Cui, X., and Xiao, L. (2025). Complexity of the Hypothalamic Oxytocin System and its Involvement in Brain Functions and Diseases. Neurosci Bull 41, 1267–1288. 10.1007/s12264-025-01424-1.

29. Althammer, F., and Grinevich, V. (2018). Diversity of oxytocin neurones: Beyond magno-and parvocellular cell types? Journal of Neuroendocrinology 30, e12549.

30. Walum, H., and Young, L.J. (2018). The neural mechanisms and circuitry of the pair bond. Nat Rev Neurosci 19, 643–654. 10.1038/s41583-018-0072-6.

31. Jurek, B., and Neumann, I.D. (2018). The Oxytocin Receptor: From Intracellular Signaling to Behavior. Physiol Rev 98, 1805–1908. 10.1152/physrev.00031.2017.

32. Lee, H.J., Macbeth, A.H., Pagani, J.H., and Young, W.S., 3rd (2009). Oxytocin: the great facilitator of life. Prog Neurobiol 88, 127–151. 10.1016/j.pneurobio.2009.04.001.

33. Qu, Y., Zhang, L., Hou, W., Liu, L., Liu, J., Li, L., Guo, X., Li, Y., Huang, C., He, Z., and Tai, F. (2024). Distinct medial amygdala oxytocin receptor neurons projections respectively control consolation or aggression in male mandarin voles. Nat Commun 15, 8139. 10.1038/s41467-024-51652-8.

34. Marlin, B.J., Mitre, M., D’Amour J, A., Chao, M.V., and Froemke, R.C. (2015). Oxytocin enables maternal behaviour by balancing cortical inhibition. Nature 520, 499–504. 10.1038/nature14402.

35. Yu, H., Miao, W., Ji, E., Huang, S., Jin, S., Zhu, X., Liu, M.Z., Sun, Y.G., Xu, F., and Yu, X. (2022). Social touch-like tactile stimulation activates a tachykinin 1-oxytocin pathway to promote social interactions. Neuron 110, 1051–1067 e1057. 10.1016/j.neuron.2021.12.022.

36. Zhang, B., Qiu, L., Xiao, W., Ni, H., Chen, L., Wang, F., Mai, W., Wu, J., Bao, A., Hu, H., et al. (2021). Reconstruction of the Hypothalamo-Neurohypophysial System and Functional Dissection of Magnocellular Oxytocin Neurons in the Brain. Neuron 109, 331–346 e337. 10.1016/j.neuron.2020.10.032.

37. Wolf, D., Hartig, R., Zhuo, Y., Scheller, M.F., Articus, M., Moor, M., Grinevich, V., Linster, C., Russo, E., Weber-Fahr, W., et al. (2024). Oxytocin induces the formation of distinctive cortical representations and cognitions biased toward familiar mice. Nat Commun 15, 6274. 10.1038/s41467-024-50113-6.

38. Ferguson, J.N., Young, L.J., Hearn, E.F., Matzuk, M.M., Insel, T.R., and Winslow, J.T. (2000). Social amnesia in mice lacking the oxytocin gene. Nat Genet 25, 284–288. 10.1038/77040.

39. Liu, S., Yang, Q., Zhu, P., Liu, X., Lu, Q., Yang, J., Gao, J., Han, H., Zhang, Z., Gu, N., et al. (2025). Precise Magnetic Stimulation of the Paraventricular Nucleus Improves Sociability in a Mouse Model of ASD. Neurosci Bull. 10.1007/s12264-025-01444-x.

40. Zelmanoff, D.D., Bornstein, R., Kaufman, M., Dine, J., Wietek, J., Litvin, A., Abraham, S., Cohen, S., Atzmon, A., Porat, I., and Yizhar, O. (2025). Oxytocin signaling regulates maternally directed behavior during early life. Science 389, eado5609. 10.1126/science.ado5609.

41. Russell, Y.I. (2022). Allogrooming. In Encyclopedia of Animal Cognition and Behavior, (Springer), pp. 183–186.

42. Choe, H.K., Reed, M.D., Benavidez, N., Montgomery, D., Soares, N., Yim, Y.S., and Choi, G.B. (2015). Oxytocin mediates entrainment of sensory stimuli to social cues of opposing valence. Neuron 87, 152–163.

43. Mitre, M., Marlin, B.J., Schiavo, J.K., Morina, E., Norden, S.E., Hackett, T.A., Aoki, C.J., Chao, M.V., and Froemke, R.C. (2016). A Distributed Network for Social Cognition Enriched for Oxytocin Receptors. J Neurosci 36, 2517–2535. 10.1523/JNEUROSCI.2409-15.2016.

44. Li, H., Jiang, T., An, S., Xu, M., Gou, L., Ren, B., Shi, X., Wang, X., Yan, J., and Yuan, J. (2024). Single-neuron projectomes of mouse paraventricular hypothalamic nucleus oxytocin neurons reveal mutually exclusive projection patterns. Neuron 112, 1081–1099. e1087.

45. Son, S., Manjila, S.B., Newmaster, K.T., Wu, Y.T., Vanselow, D.J., Ciarletta, M., Anthony, T.E., Cheng, K.C., and Kim, Y. (2022). Whole-Brain Wiring Diagram of Oxytocin System in Adult Mice. J Neurosci 42, 5021–5033. 10.1523/JNEUROSCI.0307-22.2022.

46. Cui, X., and Xiao, L. (2025). Complexity of the Hypothalamic Oxytocin System and its Involvement in Brain Functions and Diseases: X. Cui et al.: Diversity of Hypothalamic Oxytocin System. Neuroscience Bulletin, 1–22.

47. Wesson, D.W. (2020). The Tubular Striatum. J Neurosci 40, 7379–7386. 10.1523/JNEUROSCI.1109-20.2020.

48. Giessel, A.J., and Datta, S.R. (2014). Olfactory maps, circuits and computations. Curr Opin Neurobiol 24, 120–132. 10.1016/j.conb.2013.09.010.

49. Ikemoto, S. (2010). Brain reward circuitry beyond the mesolimbic dopamine system: a neurobiological theory. Neurosci Biobehav Rev 35, 129–150. 10.1016/j.neubiorev.2010.02.001.

50. Xiong, A., and Wesson, D.W. (2016). Illustrated Review of the Ventral Striatum’s Olfactory Tubercle. Chem Senses 41, 549–555. 10.1093/chemse/bjw069.

51. Sun, C., Yin, Z., Li, B.Z., Du, H., Tang, K., Liu, P., Hang Pun, S., Lei, T.C., and Li, A. (2021). Oxytocin modulates neural processing of mitral/tufted cells in the olfactory bulb. Acta Physiologica 231, e13626.

52. Oettl, L.-L., Ravi, N., Schneider, M., Scheller, M.F., Schneider, P., Mitre, M., da Silva Gouveia, M., Froemke, R.C., Chao, M.V., and Young, W.S. (2016). Oxytocin enhances social recognition by modulating cortical control of early olfactory processing. Neuron 90, 609–621.

53. Wolf, D., Hartig, R., Zhuo, Y., Scheller, M.F., Articus, M., Moor, M., Grinevich, V., Linster, C., Russo, E., and Weber-Fahr, W. (2024). Oxytocin induces the formation of distinctive cortical representations and cognitions biased toward familiar mice. Nature Communications 15, 6274.

54. Zhang, Y.F., Vargas Cifuentes, L., Wright, K.N., Bhattarai, J.P., Mohrhardt, J., Fleck, D., Janke, E., Jiang, C., Cranfill, S.L., Goldstein, N., et al. (2021). Ventral striatal islands of Calleja neurons control grooming in mice. Nat Neurosci 24, 1699–1710. 10.1038/s41593-021-00952-z.

55. Zhang, Z., Zhang, H., Wen, P., Zhu, X., Wang, L., Liu, Q., Wang, J., He, X., Wang, H., and Xu, F. (2017). Whole-Brain Mapping of the Inputs and Outputs of the Medial Part of the Olfactory Tubercle. Front Neural Circuits 11, 52. 10.3389/fncir.2017.00052.

56. Warfvinge, K., Krause, D., and Edvinsson, L. (2020). The distribution of oxytocin and the oxytocin receptor in rat brain: relation to regions active in migraine. The journal of headache and pain 21, 1–13.

57. Gimpl, G., and Fahrenholz, F. (2001). The oxytocin receptor system: structure, function, and regulation. Physiological reviews 81, 629–683.

58. Yoshimura, R., Kiyama, H., Kimura, T., Araki, T., Maeno, H., Tanizawa, O., and Tohyama, M. (1993). Localization of oxytocin receptor messenger ribonucleic acid in the rat brain. Endocrinology 133, 1239–1246. 10.1210/endo.133.3.8396014.

59. Lee, H.J., Caldwell, H.K., Macbeth, A.H., Tolu, S.G., and Young, W.S., 3rd (2008). A conditional knockout mouse line of the oxytocin receptor. Endocrinology 149, 3256–3263. 10.1210/en.2007-1710.

60. Gong, S., Doughty, M., Harbaugh, C.R., Cummins, A., Hatten, M.E., Heintz, N., and Gerfen, C.R. (2007). Targeting Cre recombinase to specific neuron populations with bacterial artificial chromosome constructs. J Neurosci 27, 9817–9823. 10.1523/JNEUROSCI.2707-07.2007.

61. Chung, W., Choi, S.Y., Lee, E., Park, H., Kang, J., Park, H., Choi, Y., Lee, D., Park, S.G., Kim, R., et al. (2015). Social deficits in IRSp53 mutant mice improved by NMDAR and mGluR5 suppression. Nat Neurosci 18, 435–443. 10.1038/nn.3927.

62. Jamain, S., Radyushkin, K., Hammerschmidt, K., Granon, S., Boretius, S., Varoqueaux, F., Ramanantsoa, N., Gallego, J., Ronnenberg, A., Winter, D., et al. (2008). Reduced social interaction and ultrasonic communication in a mouse model of monogenic heritable autism. Proc Natl Acad Sci U S A 105, 1710–1715. 10.1073/pnas.0711555105.

63. Zhang, Y.F., Wu, J., Wang, Y., Johnson, N.L., Bhattarai, J.P., Li, G., Wang, W., Guevara, C., Shoenhard, H., Fuccillo, M.V., et al. (2023). Ventral striatal islands of Calleja neurons bidirectionally mediate depression-like behaviors in mice. Nat Commun 14, 6887. 10.1038/s41467-023-42662-z.

64. Wang, W., Zhong, Y., Tan, R., Wang, M., Liu, J., Wang, D., Wang, H., Li, Y., Li, G., Yang, J., et al. (2025). A midbrain-to-ventral-striatum dopaminergic pathway orchestrates odor-guided insect predation in mice. Proc Natl Acad Sci U S A 122, e2514847122. 10.1073/pnas.2514847122.

65. Panksepp, J.B., and Lahvis, G.P. (2011). Rodent empathy and affective neuroscience. Neurosci Biobehav Rev 35, 1864–1875. 10.1016/j.neubiorev.2011.05.013.

66. Ueno, H., Suemitsu, S., Murakami, S., Kitamura, N., Wani, K., Okamoto, M., Matsumoto, Y., Aoki, S., and Ishihara, T. (2018). Empathic behavior according to the state of others in mice. Brain Behav 8, e00986. 10.1002/brb3.986.

67. Gonzalez-Liencres, C., Juckel, G., Tas, C., Friebe, A., and Brune, M. (2014). Emotional contagion in mice: the role of familiarity. Behav Brain Res 263, 16–21. 10.1016/j.bbr.2014.01.020.

68. Kim, S.W., Kim, M., and Shin, H.S. (2021). Affective empathy and prosocial behavior in rodents. Curr Opin Neurobiol 68, 181–189. 10.1016/j.conb.2021.05.002.

69. Wu, Y.E., and Hong, W. (2022). Neural basis of prosocial behavior. Trends Neurosci 45, 749–762. 10.1016/j.tins.2022.06.008.

70. Suzukawa, K., Kondo, K., Kanaya, K., Sakamoto, T., Watanabe, K., Ushio, M., Kaga, K., and Yamasoba, T. (2011). Age-related changes of the regeneration mode in the mouse peripheral olfactory system following olfactotoxic drug methimazole-induced damage. J Comp Neurol 519, 2154–2174. 10.1002/cne.22611.

71. Zhang, Y.F., Janke, E., Bhattarai, J.P., Wesson, D.W., and Ma, M. (2022). Self-directed orofacial grooming promotes social attraction in mice via chemosensory communication. iScience 25, 104284. 10.1016/j.isci.2022.104284.

72. Barabas, A.J., Aryal, U.K., and Gaskill, B.N. (2019). Proteome characterization of used nesting material and potential protein sources from group housed male mice, Mus musculus. Sci Rep 9, 17524. 10.1038/s41598-019-53903-x.

73. Yoshimura, R., Kimura, T., Watanabe, D., and Kiyama, H. (1996). Differential expression of oxytocin receptor mRNA in the developing rat brain. Neurosci Res 24, 291–304. 10.1016/0168-0102(95)01003-3.

74. Schwartz, J.-C., Diaz, J., Bordet, R., Griffon, N., Perachon, S., Pilon, C., Ridray, S., and Sokoloff, P. (1998). Functional implications of multiple dopamine receptor subtypes: the D1/D3 receptor coexistence. Brain research reviews 26, 236–242.

75. Ridray, S., Griffon, N., Mignon, V., Souil, E., Carboni, S., Diaz, J., Schwartz, J.C., and Sokoloff, P. (1998). Coexpression of dopamine D1 and D3 receptors in islands of Calleja and shell of nucleus accumbens of the rat: opposite and synergistic functional interactions. European Journal of Neuroscience 10, 1676–1686.

76. Hu, B., Boyle, C.A., and Lei, S. (2020). Oxytocin receptors excite lateral nucleus of central amygdala by phospholipase Cbeta- and protein kinase C-dependent depression of inwardly rectifying K(+) channels. J Physiol 598, 3501–3520. 10.1113/JP279457.

77. Karschin, C., Dissmann, E., Stuhmer, W., and Karschin, A. (1996). IRK(1-3) and GIRK(1-4) inwardly rectifying K+ channel mRNAs are differentially expressed in the adult rat brain. J Neurosci 16, 3559–3570. 10.1523/JNEUROSCI.16-11-03559.1996.

78. Radford, A.N. (2012). Post-allogrooming reductions in self-directed behaviour are affected by role and status in the green woodhoopoe. Biology letters 8, 24–27.

79. Yates, K., Stanley, C.R., and Bettridge, C.M. (2022). The effects of allogrooming and social network position on behavioural indicators of stress in female lion-tailed macaques (Macaca silenus). Behav Process 202, 104740.

80. Lim, K.Y., and Hong, W. (2023). Neural mechanisms of comforting: Prosocial touch and stress buffering. Hormones and behavior 153, 105391.

81. Chen, P., and Hong, W. (2018). Neural Circuit Mechanisms of Social Behavior. Neuron 98, 16–30. 10.1016/j.neuron.2018.02.026.

82. Grinevich, V., and Stoop, R. (2018). Interplay between Oxytocin and Sensory Systems in the Orchestration of Socio-Emotional Behaviors. Neuron 99, 887–904. 10.1016/j.neuron.2018.07.016.

83. Oettl, L.L., Ravi, N., Schneider, M., Scheller, M.F., Schneider, P., Mitre, M., da Silva Gouveia, M., Froemke, R.C., Chao, M.V., Young, W.S., et al. (2016). Oxytocin Enhances Social Recognition by Modulating Cortical Control of Early Olfactory Processing. Neuron 90, 609–621. 10.1016/j.neuron.2016.03.033.

84. Freda SN, P.M., Badong D, Xiao L, Liu Y, Kozorovitskiy Y (2022). Brainwide Input-Output Architecture of Paraventricular Oxytocin and Vasopressin Neurons. bioRxiv. 10.1101/2022.01.17.476652.

85. Tost, H., Kolachana, B., Hakimi, S., Lemaitre, H., Verchinski, B.A., Mattay, V.S., Weinberger, D.R., and Meyer-Lindenberg, A. (2010). A common allele in the oxytocin receptor gene (OXTR) impacts prosocial temperament and human hypothalamic-limbic structure and function. Proc Natl Acad Sci U S A 107, 13936–13941. 10.1073/pnas.1003296107.

86. Kirckof, A., Kneller, E., Vitale, E.M., Johnson, M.A., and Smith, A.S. (2025). The effects of social loss and isolation on partner odor investigation and dopamine and oxytocin receptor expression in female prairie voles. Neuropharmacology 267, 110298. 10.1016/j.neuropharm.2025.110298.

87. Narita, K., Murata, T., and Matsuoka, S. (2016). The ventromedial hypothalamus oxytocin induces locomotor behavior regulated by estrogen. Physiol Behav 164, 107–112. 10.1016/j.physbeh.2016.05.047.

88. Menon, R., and Neumann, I.D. (2023). Detection, processing and reinforcement of social cues: regulation by the oxytocin system. Nat Rev Neurosci 24, 761–777. 10.1038/s41583-023-00759-w.

89. Schumacher, M., Coirini, H., Flanagan, L.M., Frankfurt, M., Pfaff, D.W., and McEwen, B.S. (1992). Ovarian steroid modulation of oxytocin receptor binding in the ventromedial hypothalamus. Ann N Y Acad Sci 652, 374–386. 10.1111/j.1749-6632.1992.tb34368.x.

90. Zheng, Q., Zhao, Y., Cheng, Q., Wang, H., Liu, F., Lai, J., Liu, Y., Zhang, X., Kang, Y., Li, Z., et al. (2025). Potentiation of nigra-striatal dopaminergic projection underpins core autism-like behaviors in valproate-exposed mice. J Neurosci. 10.1523/JNEUROSCI.0382-25.2025.

91. Choi, J.E., Choi, D.I., Lee, J., Kim, J., Kim, M.J., Hong, I., Jung, H., Sung, Y., Kim, J.I., Kim, T., et al. (2022). Synaptic ensembles between raphe and D1R-containing accumbens shell neurons underlie postisolation sociability in males. Science Advances 8. ARTN eabo7527 10.1126/sciadv.abo7527.

92. Meyer, M.A.A., Anstotz, M., Ren, L.Y., Fiske, M.P., Guedea, A.L., Grayson, V.S., Schroth, S.L., Cicvaric, A., Nishimori, K., Maccaferri, G., and Radulovic, J. (2020). Stress-related memories disrupt sociability and associated patterning of hippocampal activity: a role of hilar oxytocin receptor-positive interneurons. Transl Psychiatry 10, 428. 10.1038/s41398-020-01091-y.

93. Xiao, L., Priest, M.F., Nasenbeny, J., Lu, T., and Kozorovitskiy, Y. (2017). Biased Oxytocinergic Modulation of Midbrain Dopamine Systems. Neuron 95, 368–384 e365. 10.1016/j.neuron.2017.06.003.

94. Hung, L.W., Neuner, S., Polepalli, J.S., Beier, K.T., Wright, M., Walsh, J.J., Lewis, E.M., Luo, L., Deisseroth, K., Dolen, G., and Malenka, R.C. (2017). Gating of social reward by oxytocin in the ventral tegmental area. Science 357, 1406–1411. 10.1126/science.aan4994.

95. Young, L.J., and Wang, Z. (2004). The neurobiology of pair bonding. Nat Neurosci 7, 1048–1054. 10.1038/nn1327.

96. Dolen, G., Darvishzadeh, A., Huang, K.W., and Malenka, R.C. (2013). Social reward requires coordinated activity of nucleus accumbens oxytocin and serotonin. Nature 501, 179–184. 10.1038/nature12518.

